# Antiviral State of CD1c+ cDC contributes to Increased Maturation and Activation of Cytotoxic Natural Killer cells in Sjögren’s Syndrome

**DOI:** 10.1101/2022.03.13.484063

**Authors:** I Sánchez-Cerrillo, M Calvet-Mirabent, A Triguero-Martínez, D Calzada-Fraile, C Delgado-Arévalo, M Valdivia-Mazeyra, M Ramírez-Huesca, E Vázquez de Luis, A Benguría-Filippini, R Moreno-Vellisca, M Adrados de Llano, H de la Fuente, Ilya Tsukalov, E Roy-Vallejo, A Ramiro, F Sánchez-Madrid, A Dopazo, I González Álvaro, S Castañeda, E Martin-Gayo

## Abstract

**Objective:** Primary Sjögren’s syndrome (pSS) is an inflammatory autoimmune disorder characterized by damage of exocrine glands and linked to IFN responses and the induction of autoreactive adaptive immune cells. However, the role of innate immune cells in pSS pathology remains understudied.

**Methods:** We studied differential phenotypical characteristics of different NK cell, conventional dendritic cell (cDC) and monocyte subsets in the blood and salivary glands from pSS individuals. Transcriptional patterns of circulating cDC and Mo from pSS and healthy controls were also compared. Finally, *in vivo* alterations in these cell populations in the salivary gland were investigated in a mouse model.

**Results:** Here, we identified CD16+ CD56hi NK cells enriched in pSS patients which associates with higher natural cytotoxic function and increased proportions of circulating CD64+ CD1c+ cDC exhibiting antiviral transcriptional IFN signatures. CD64hi cDC and NK cell were detected infiltrated into the salivary glands from pSS patients and a murine SS model. CD1c+ cDC from patients with pSS expressed high levels of ligands for activating NK receptors and increased ability to activate NK cells *ex vivo*. Finally, the antiviral RIG-I and DDX60 sensors regulated the expression of NK cell receptor ligands on CD1c+ cDC.

**Conclusions:** Therefore, the interplay of CD1c+ cDCs and NK cells could contribute to pSS pathology.

## INTRODUCTION

Primary Sjögren’s syndrome (pSS) is a complex autoimmune disorder usually characterized by inflammation of salivary and/or lacrimal glands, and in some cases extra-glandular manifestations, ^1^. Altered adaptive immune responses mediated by Th17 cells, ^2-4^ CD8+ T cells, ^5, 6^ and autoreactive B cells have been well characterized and associated with glandular tissue destruction, ^7, 8^. However, the activation of interferon (IFN) responses in pSS patients suggests that innate immunity plays an active role during the development and/or progression of the pathology, ^9-13^. Innate immune cell types such as monocytes (Mo), dendritic cells (DC) and Natural Killer (NK) cells might be altered in pSS patients, ^2, 14-16^ but the phenotypical and functional alterations of specific subsets and their contribution to pathology remain unexplored. Inflammatory CD16+ monocytes (Mo), plasmacytoid DCs (pDC) and CD11c+ conventional DCs (cDC) are recruited to the salivary glands (SGs) in pSS patients, especially at initial stages of the disease, ^17-22^. cDC is a heterogeneous group comprised by CD1c+ and CD141+ cDC subsets, with specific functional and inflammatory properties induced by different types of PAMPs, ^23-25^. They can induce the activation and maturation of NK cells, ^26, 27^ which might contribute to autoimmunity, ^28^. Recognition of endogenous RNA and DNA can aberrantly activate myeloid cells and could play a role in pSS, ^10, 29^. In the present study, we assessed alterations in subset distribution and functional properties of NK cells and cDCs from pSS patients. Here, we provide novel insights on innate immune cellular mechanisms potentially driving typical histological disturbances in pSS and identifying new potential targets for future therapies.

## METHODS

### Study Participants

The current study included n=38 primary Sjögren’s syndrome (pSS) and n=43 healthy donors (HD). All pSS patients assessed in a single cross-sectional visit and all of them fulfilled the 2016 ACR-EULAR classification criteria for pSS, ^30^. Biological samples were collected at the visit and stored at the Instituto de Investigación Sanitaria La Princesa (IIS-IP) Biobank. To evaluate the disease activity, recruited pSS patients were classified according to the (ESSDAI) index, ^31, 32^. The clinical characteristics and treatments of pSS patients are summarized in Supplemental Table 1. PBMC samples from HD were isolated from Buffy coats obtained from the Centro de Transfusiones Comunidad de Madrid and used as controls for comparison purposes.

### Ethic statement

All subjects participating in the study gave written informed consent, and the study was approved by the Institutional Review Board of Hospital Universitario de La Princesa (Protocol Number #3515) and following the Helsinki declaration. For *in vivo* experiments, mice were housed at the animal facility from Centro Nacional de Investigaciones Cardiovasculares. Animal experiments were approved by the local ethics committee and are in agreement with the EU Directive 86/609/EEC, Recommendation 2007/526/EC and Real Decreto 53/2013.

### Patient involvement

Patients recruited for our study were only informed about the objective and outcomes of the research, but they were not involved in the design, conduct, reporting or dissemination plan of the study.

### Flow cytometry analysis and cell sorting

*Ex vivo* and cultured PBMC were stained with APC-H7 (Tonbo Biosciences, San Diego, CA) or Brilliant Violet 405 (Molecular Probes, Eugene, OR) viability dye in the presence of different panels of monoclonal antibodies directed to human CD3, CD19, CD20, CD56, CD14, CD16, HLA-DR, CD11c, CD1c, CD141 and CD64 (Biolegend, San Diego, CA). Additional panels using anti-human CD56, CD16, SLAMF7, PCNA, MICAB, ULBP1, NKG2D, CD107a, TNFα, IFNγ (BioLegend, San Diego, CA) and NKG2A (RyD systems, Minneapolis, MN) were used for validation purposes (see Supplemental Table 3). For *in vivo* experiments, anti-mouse CD3, CD11b, NK1.1 (Tonbo Biosciences), CD27, XCR1, CD107a (BioLegend), CD45.2, IFNγ, CD11b (eBioscience), CD64, CD11c, MCH-II, Ly6C, CD3, CD45.2 (BD Bioscience) and NKG2D (ThermoFisher) (see Supplemental Table 3). Samples were analyzed on a Fortessa cytometer (BD Biosciences, San Jose, CA) at Centro Nacional de Investigaciones Cardiovasculares (CNIC, Madrid, Spain). Analysis of individual and multiparametric flow cytometry data was performed using FlowJo software (Tree Star). Human CD1c+ cDC, CD141+ cDC, Mo and CD56+ NK cells were sorted from PBMCs, using a FACS Aria II sorter (BD Biosciences), from pSS patients (untreated) and HD following the next gating strategy: viable human Lin^-^ (CD3, CD19, CD20, CD56), CD14^-^ CD11c^+^ HLADR^+^ CD1c+ cDCs; CD14^-^ CD11c^+^ HLADR^+^ CD141+ cDCs; total Lin^-^ CD14^+^. Samples from n=4 pSS patients and n=4 HD were used for transcriptional analyses and n=8 pSS patients for *ex vivo* functional analyses. For functional analysis of NK cells, a modified panel (Lineage cocktail containing only anti-CD3, CD19, CD20 mAbs) was used allowing also the separation of Lin- HLADR- CD56+.

### Longitudinal analysis of myeloid and NK subsets in vivo in a murine model of pSS

We used an *in vivo* model of Sjögren’s syndrome previously described, ^29, 33^ which is based on the intraperitoneal injection of poly(I:C). (See details on online Supplemental Methods).

### Gene expression analysis by RNA-Seq and Computational data analysis

Total RNA was isolated from sorted primary PB Mo, CD1c+ and CD141+ cDC subsets from n=4 untreated pSS patients and n=4 HD, using the RNeasy Micro Kit (Qiagen). Quality and integrity of each RNA sample was checked using a Bioanalyzer prior to proceeding to RNAseq protocol. Subsequently, selected RNAs from cDCs were used to amplify the cDNA using the SMART-Seq v4 Ultra Low Input RNA Kit (Clontech-Takara). Amplified (1 ng) cDNA was used to generate barcoded libraries using the Nextera XT DNA library preparation kit (Illumina, San Diego, CA). The size of the libraries was checked using the Agilent 2100 Bioanalyzer High Sensitivity DNA chip and their concentration was determined using the Qubit® fluorometer (ThermoFisher Scientific, Waltham, MA).

RNA from circulating Mo was processed as follows: 200 ng of total RNA were used to generate barcoded RNA-seq libraries using the NEBNext Ultra II Directional RNA Library Prep Kit (New England Biolabs Inc). Libraries were sequenced on a HiSeq 2500 (Illumina, San Diego, CA) and processed with RTA v1.18.66.3. FastQ files for each sample were obtained using bcl2fastq v2.20.0.422 software (Illumina, San Diego, CA). Sequencing reads were aligned to the human reference transcriptome (GRCh38 v91) and quantified with RSem v1.3.1 (34). Raw counts were normalized with TPM (Transcripts per Million) and TMM (Trimmed Mean of M-values) methods, transformed into log2 expression (log2(rawCount+1)) and compared to calculate fold-change and corrected p-value. Only those genes expressed with at least 1 count in a number of samples equal to the number of replicate samples of the condition with less replicates were taken into account. Gene expression changes were considered significant if associated to Benjamini and Hochberg adjusted p-value < 0.05.

Heatmaps were generated with Morpheus online tool from Broad Institute (https://software.broadinstitute.org/morpheus). Finally, pathway analysis and visualization of gene networks for each DEG list (FDR corrected p<0.05, considering a cutoff log2Fold change in expression in pSS vs HD >1.5 and < −1.5) was performed using Ingenuity Pathway Analysis (Qiagen) and NetworkAnalyst (35) Softwares.

### Validation of gene expression by RT-qPCR

RNA was isolated from sorted PB myeloid subsets using RNeasy Micro Kit (Qiagen) according to manufacturer’s instructions. cDNA was synthesized using the reverse transcription kit (Promega) and transcriptional expression of IFIT1 (Fw: 5’-GGAATACACAACCTACTAGCC-3’; Rv: 5’-CCAGGTCACCAGACTCCTCA-3’), IFIT3 (Fw: 5’-TGAGGAAGGGTGGACACAACTGAA-3’; Rv: 5’-AGGAGAATTCTGGGTTGTTGGGCT-3’), DDX60 (Fw 5’- AAGGTGTTCCTTGATGATCTCC-3’ Rv : 5’ -TGACAATGGGAGTTGATATTCC-3’) as analyzed by semiquantitative PCR using the SYBR Green assay GoTaq® qPCR Master Mix (Promega) with standardized primers (Metabion). TaqMan Gene expression Assays were also used to determine transcriptional expression of RIG-I (Hs01061436_m1) and β-Actin (Hs01060665_g1). qPCR amplifications were performed on a StepOne Real-Time PCR system (Applied Biosystems). Relative gene expression was normalized to β-Actin detection.

### siRNA-mediated gene knockdown

Gene knockdown of DDX60 and DDX58 (RIG-I) was performed by nucleofection of primary cDCs isolated from HD with specific siRNAs (SMART-pool, Horizon Discovery) or irrelevant scramble siRNA in an Amaxa4D-Nucleofector (Lonza) instrument using the CM120 protocol and following the manufacturer’s instructions. siRNA-mediated knockdown was confirmed after 24h by RT-qPCR analysis of mRNA levels for target genes.

### Immunomagnetic Isolation of primary cDCs and NK cells

Total CD1c+ cDCs were purified from total PBMC suspensions by negative immunomagnetic methods (purity >90%) using the Human Myeloid DC Enrichment Kit (STEMCELL). Untouched NK cells were also isolated by immunomagnetic selection from autologous PBMC samples using a negative selection kit (purity >90%) “EasySepTM Human NK Cell Isolation Kit (STEM CELL)”.

### Killing Assays

Natural cytotoxic function of circulating NK cells from HD and pSS patients was tested *in vitro* by coculture with the target Green Fluorescent Protein (GFP) expressing K562 cell line (NIH Reagent Program 116799). Briefly, NK cells were isolated by immunomagnetic selection as described above and labelled with Violet Cell Proliferation Tracker according to manufacter’s instructions (C34557, ThermoFisher) and subsequently cultured at a NK: GFP-K562 target ratio 5:1 for 16h. GFP-K562 targets were also cultured individually in the absence of NK cells as a control. Specific Killing was determined by flow cytometry by excluding Violet Cell Tracker + NK cells and determining the proportions of cells losing GFP expression and acquiring Live-Dead Viability Staining (see Supplemental Figure 2B).

### In vitro stimulation of DCs and functional assays

CD1c+ cDCs from PBMC of HD or pSS patients were cultured for 24h in the presence of media alone or 1.5μg/ml poly(I:C) (SIGMA) according to manufacturer’s instructions. After 24h, maturation of cDC was evaluated by FACS analysis of expression of ligands for activating (MICAB, SLAMF7) and inhibitory (PCNA) receptors in NK cells. For functional assays, *ex vivo* cDCs from HD and pSS individuals were co-cultured directly with autologous CD56+ NK cells for 24h at a 1:2 ratio in media supplemented with 50 IU/ml IL-2 (Prepotech). A negative control of NK cell activation cultured in media only was also included. After 24h of culture, expression of CD16, CD56 was analyzed on gated NK cells by flow cytometry. The same conditions were used in functional assays performed with cDCs nucleofected with siRNAs and exposed to Media or poly(I:C), as previously described. In some experiments, media was supplemented with 1.5 μg/ml Brefeldin A (SIGMA) and Monensin (SIGMA) and anti-human CD107a mAbs for subsequent analysis of degranulation and intracellular expression of IFNγ and TNFα by flow cytometry at the end of the assay.

### Histological analysis of pSS salivary glands by Immunofluorescence

Salivary gland biopsies from n=7 pSS patients were embedded in paraffin and segmented in 3 μm fragments. Tissue sections deparaffinization, hydration and target retrival was performed with a PT-LINK (Dako) before immune-staining. For immuno-staining of paraffin-preserved tissue we used goat anti-human CD56 (RyD Systems), mouse anti-human CD1c cDC (abcam), rat anti-human Granzyme B (eBioscience), rabbit anti-human HLA-DR (abcam), as primary antibodies; and donkey anti-rabbit AF488 (Invitrogen), donkey anti-rat AF594 (Jackson ImmunoResearch), donkey anti-goat AF568, donkey anti-goat AF647 (Invitrogen) and donkey anti-mouse AF647 (Thermo Fisher) were used as secondary antibodies. Images were taken with a Leica TCS SP5 confocal and processed with the LAS AF software. Detection of CD1c+ cDCs, HLADR+ cDCs, CD56+ NK cells and their co-localization with Granzyme B were analyzed with ImageJ software. In some cases, SG tissue sections were also stained with hematoxylin and eosin to discriminate extracellular matrix defining areas with different levels of infiltrates.

### Statistics

The statistical analysis to evaluate differences between the cells from different or within the same patient cohorts (pSS or HD) were assessed using Mann Whitney U or Wilcoxon matched-pairs signed-rank tests. For longitudinal *in vivo* studies, a Two-Way ANOVA test including correction for multiple comparisons was applied. Non-parametric Spearman correlation was performed to test association of mentioned parameters. A p value less than 0.05 was considered significant. Statistical analyses were performed using GraphPad Prism 9.0 software.

### Data sharing statement

RNA-seq data from the study has been deposited on the public Gene Expression Omnibus (GEO) repository, accession number: GSE157047 (https://www.ncbi.nlm.nih.gov/geo/query/acc.cgi?acc=GSE157047) and GSE194263 (https://www.ncbi.nlm.nih.gov/geo/query/acc.cgi?acc=GSE194263).

## RESULTS

### Altered distribution and increased cytotoxic function of NK cells from pSS patients

Compared to HD, in the blood of pSS patients CD56hi CD16+ NK cells were significantly increased, while significantly lower frequencies of CD56dim CD16-subset were detected (Figure 1A-B). The remaining NK subsets were not affected (Supplemental Figure 1A). In addition, while all CD16+ NK cell subsets from pSS expressed higher basal levels of the degranulation marker CD107a, only CD56hi CD16+ and CD56dim CD16+ NK cells specifically were enriched in significantly higher proportions of polyfunctional IFNγ+ CD107a+ cells TNFα+ cells compared to HD (Supplemental Figure 1B-C, Figure 1C). Interestingly, increased expression of the activating NK cell receptor NKG2D was observed in all NK subsets from blood of pSS, but less significant differences were observed in expression of the activating receptor SLAMF7 and the inhibitory receptor NKG2A (Supplemental Figure 2A). Furthermore, NK cells from pSS patients exhibited significantly higher cytotoxic function towards K562 target cells compared to cells from HD (Figure 1D; Supplemental Figure 2B). In addition, CD56+ NK cells from infiltrated areas in the SG from pSS patients co-expressed granzyme B (Figure 1E, Supplemental Figure 1D-E). Together, our data indicate that NK cells from pSS individuals are enriched on cell subsets characterized by a more mature and cytotoxic state.

**Figure 1.**
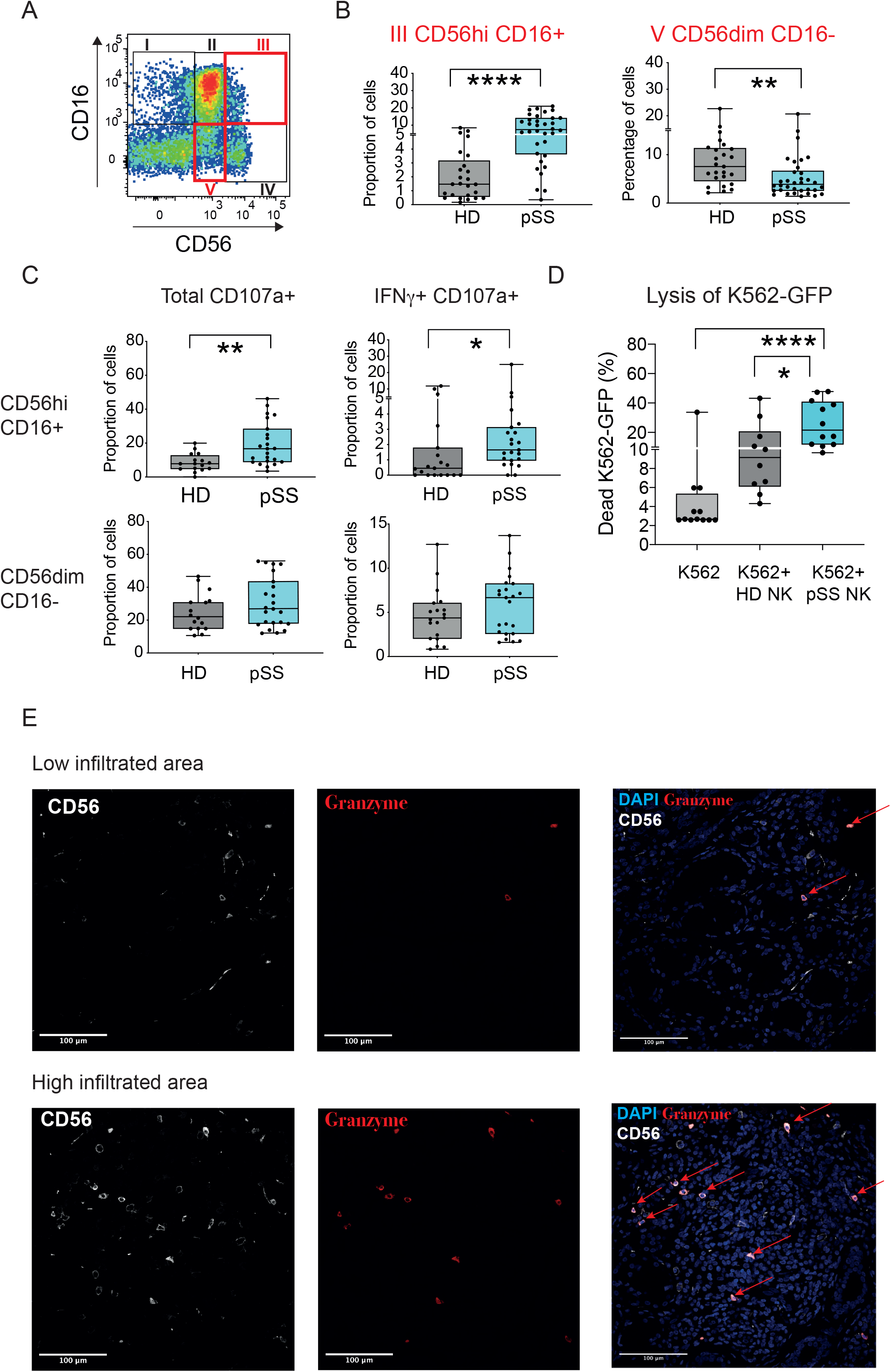
Proportions and phenotypical characteristics of natural killer cell subsets from pSS patients. (A): Representative flow cytometry dot plot showing analysis of NK cell subsets present in Lineage (CD3, CD19, CD14) negative, HLADR negative lymphocytes and defined by CD16 versus CD56 expression in the PB of our study cohort. (B): Summarized proportions of circulating CD56hi CD16+ (B, left panel) and CD56dim CD16-NK cells (B, right panel) in 25 healthy donors (HD; grey) and 34 primary Sjögren’s syndrome (pSS; blue) patients. (C): Proportions of cells expressing CD107a (left) and IFNγ+ CD107a+ (right) (C) within the indicated circulating NK cell subsets from n=16 HD (grey) and n=23 pSS patients (blue). Statistical significance was calculated using a two-tailed Mann Whitney test. *p<0.05; **p<0.01; ****p<0.0001. (D): Functional characteristics of NK cells from pSS patients. Summary of proportions of dead target K562-GFP cells cultured for 16h in the absence (light grey) or the presence of isolated circulating NK cells from HD or pSS patients. Statistical significance was calculated with a two-tailed Mann Whitney test. *p<0.05; ****p<0.0001. Statistical significance was calculated using a two-tailed Mann Whitney test. *p<0.05; **p<0.01; ****p<0.0001. (B-C-D). (E): Histological immunofluorescence analysis of expression of CD56 (white) and Granzyme B (red) on low (upper images) and highly (lower images) infiltrated glandular areas from the section of SG tissue from a representative pSS patient. Cell nuclei were stained with DAPI (blue). Cells co-expressing both markers are highlighted with red arrows. Original magnification 40X.

### Reduced frequencies and upregulation of CD64 in circulating CD1c+ cDCs from pSS patients

Next, we studied frequencies and phenotype of circulating pDCs, CD1c+ and CD141+ cDC subsets in the PB of pSS and HD (Supplemental Figure 3A). A non-significant trend to lower frequency of pDCs was observed in pSS patients compared to HD (Supplemental Figure 3B). However, proportions of circulating inflammatory CD14lo CD16hi non-classical (NC) Mo were significantly increased in PB of pSS patients, in agreement with previous studies, ^17^ while CD14+ CD16int transitional (T) and CD14+ CD16-classical (C) Mo were not significantly different (Figure 2A; Supplemental Figure 3A-C). In contrast, proportions of both CD1c+ and CD141+ cDC were markedly reduced in the PB of pSS individuals (Figure 2A). Moreover, HLADR+ cells with myeloid morphology and co-expressing CD1c could be found in infiltrated areas in proximity to CD56+ NK in the SGs of pSS (Figure 2B). In terms of phenotype, we observed that expression levels of CD64, associated with activation or response to IFN, ^36, 37^ and a marker for antiviral cDCs, ^25^ were significantly higher in Mo and CD1c+ cDCs from pSS PB (Figure 2C; Supplemental Figure 3D).

**Figure 2.**
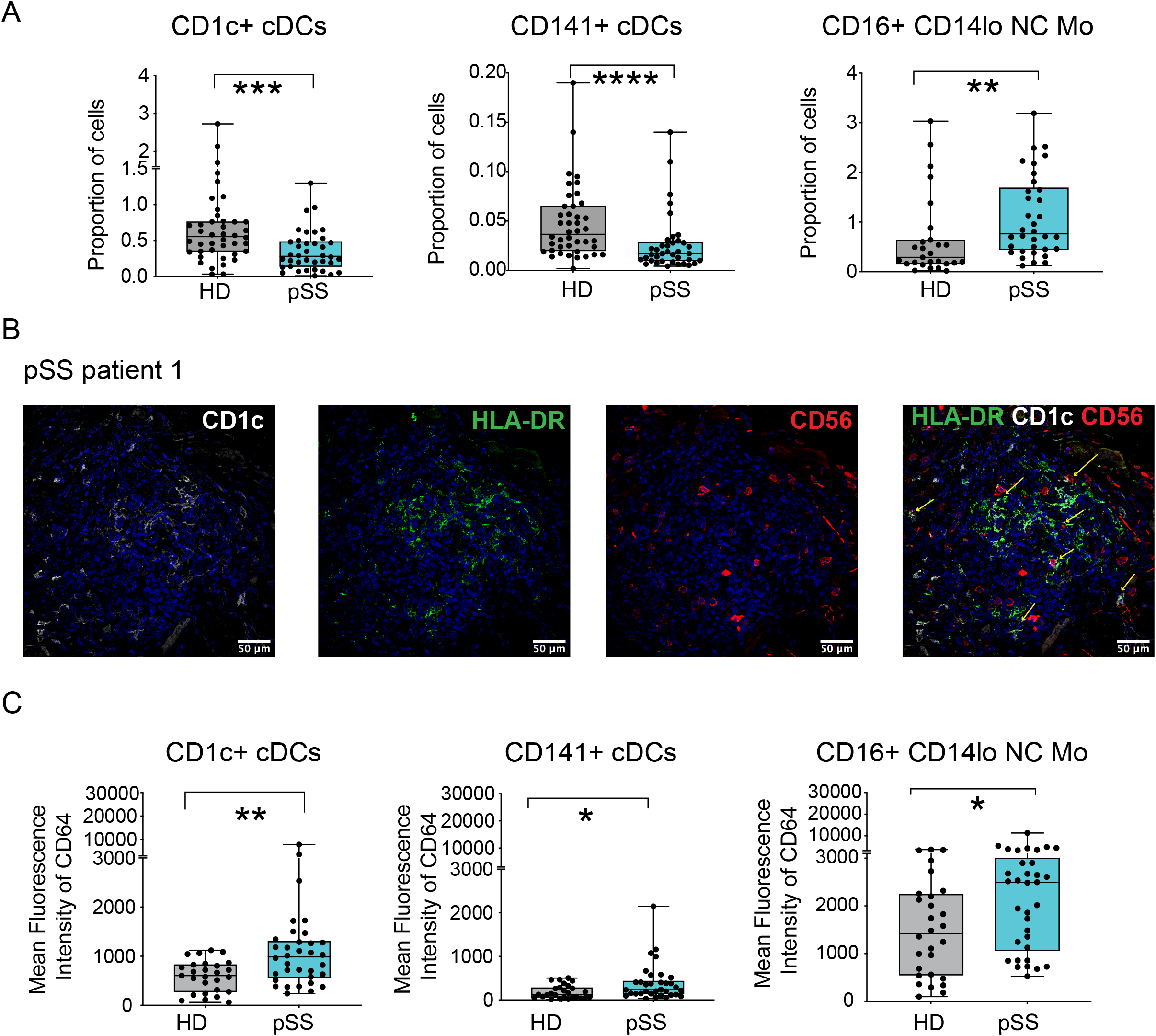
Frequencies and phenotypic characteristics of myeloid cell subsets from pSS patients. (A, C): Box and Whiskers plots showing proportions from live lymphocytes (A) and mean of fluorescence intensity of CD64 (C) of non-classical (NC) CD16+ CD14lo Mo, CD1c+ cDC and CD141+ cDC in the PB from (violet; n=43 for analysis of proportions and n=28 for analysis of CD64) and pSS patients (blue; n=38 for analysis of proportions and n=34 for analysis of CD64). Statistical significance was calculated using a two-tailed Mann Whitney test. *p<0.05; **p<0.01; ***p<0.001; ****p<0.0001. (B): Representative confocal microscopy (40x magnification) showing immunofluorescence analysis of CD1c (white), HLADR (green) and CD56 (red) expression on a SG tissue section from a representative pSS patient. Cell nuclei were stained with DAPI (blue). Cells co-expressing CD1c and HLADR markers close proximity to CD56 cells are highlighted with a yellow arrow.

### CD64+ cDC and activated NK cell subsets are induced in an *in vivo* SS mouse model

To determine whether specific cDC subsets and NK cell activation patterns might also occur *in vivo* in the SG, a murine *in vivo* SS-like model induced by intraperitoneal injections of poly(I:C) was analyzed, ^33, 38^ and compared with mice receiving PBS (Supplemental Figure 4A). Increased infiltration of CD45.2+ hematopoietic cells into the submandibular SG (SMSG) in mice injected with poly(I:C) was observed (Supplemental Figure 4B). We observed a significant increase in the proportions of CD11chi CD11b+ cDCs, the murine homologues of human CD1c+ cDC, which also exhibited the most significant increase in CD64 expression (Figure 3A-B; Supplemental Figure 4C). In contrast, proportions of Ly6C+ Mo and XCR1+ cDCs (homologues to human CD141+ cDC) tended to decrease in the SMSG from the poly(I:C)-injected group and CD64 expression was not as significantly affected in these subsets (Supplemental Figure 4D-4E). Of note, Ly6C-CD11clo/int cells increased in these animals (Supplemental Figure 4D).

**Figure 3.**
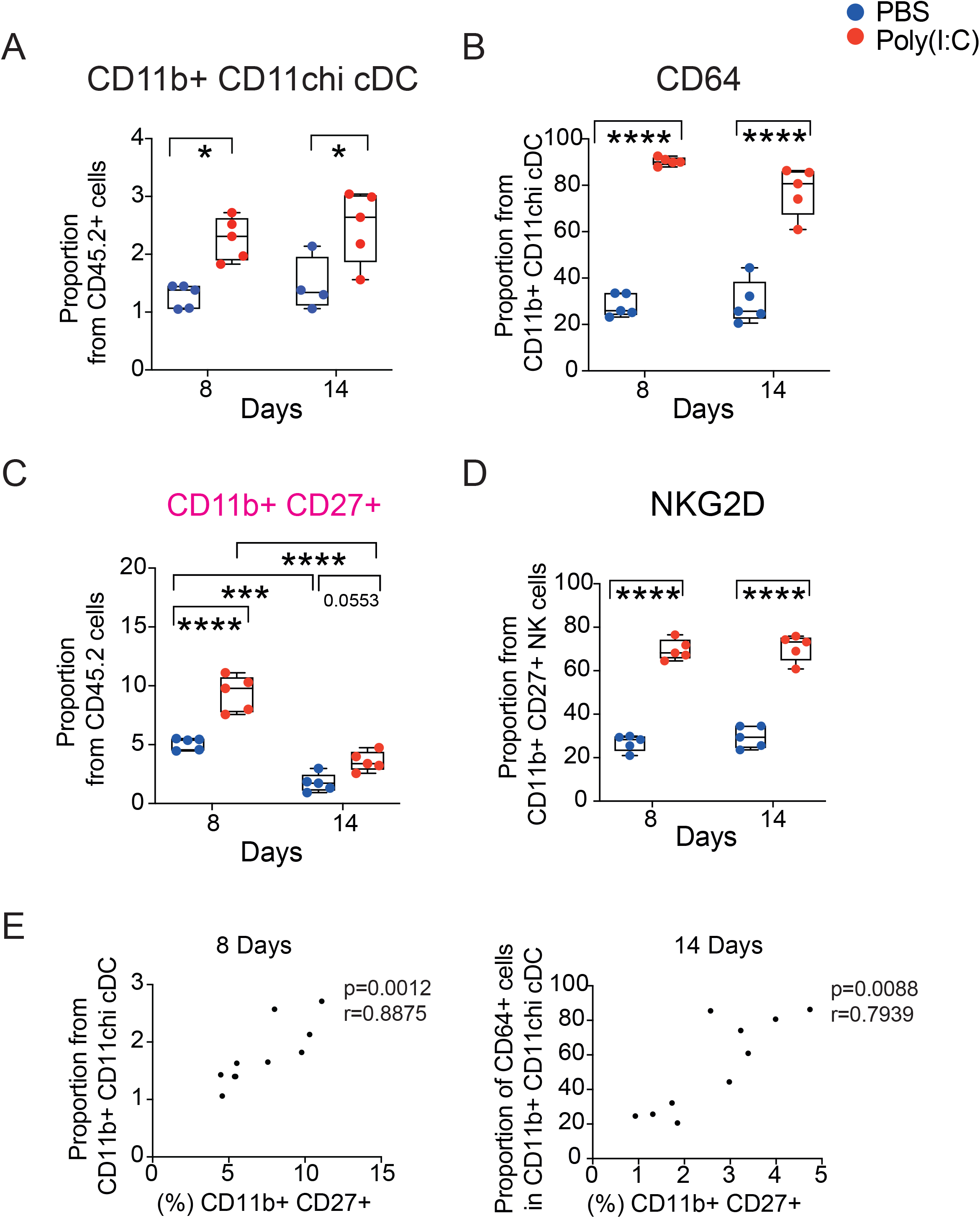
*In vivo* recruitment of activated cDC and NK cells subsets to the submandibular salivary gland in a Sjögren’s syndrome mouse model. (A): Proportions of CD11b+ CD11chi cDC in the submandibular salivary gland (SMSG) of mice injected with PBS (blue) or poly(I:C) (red) at 8 and 14 days after treatment initiation on a Sjögren’s syndrome mouse model. (B): Proportions of CD64+ cells included in the CD11b+ CD11chi cDC population infiltrated in the SMSG. (C-D): Frequencies of CD11b+CD27+ NK cell subset (C) and expression of NKG2D in this subset (D). (E): Spearman correlation between proportions of CD11b+ CD11chi cDC (left) and frequencies of CD64+ cells within this cDC subset (right) and CD11b+ CD27+ NK cell subset in the SMSG at day 8 and day 14, after treatment initiation, respectively. Spearman p- and r-values are detailed on top of each plot. These data are representative from one out of two independent experiments. Statistical significance was calculated with a two way ANOVA and Spearman test. *p<0.05; **p<0.01; ***p<0.001; ****p<0.0001.

Proportions of NK1.1+ NK cell subsets defined by differential profiles of CD27 and CD11b were assessed (Supplemental Figure 4C). Frequencies of transitional CD11blo CD27+ and CD11b+ CD27+ NK cell subsets increased in infiltrates at early time points after poly(I:C) injection but decreased at fourteen days p.i. (Figure 3C; Supplemental Figure 4F). Also, frequencies of transitional CD11b+ CD27+ NK cells were increased in the PB, similarly to NK cells from pSS patients (Supplemental Figure 4G). Moreover, proportions of infiltrated transitional CD11b+ CD27+ NK cells were positively correlated with early enrichment in CD11b+ cDC and subsequent increase on CD64 levels in these cells in the SMSG (Figure 3E). In addition, transitional murine CD11b+ CD27+ NK cells contained significantly higher levels of CD107a+ IFNγ+ cells in the PB and expressed significantly higher levels of NKG2D both in the PB and the SMSG (Figure 3D Supplemental Figure 4H). Together, our data validate the recruitment, interaction and phenotypical profiles of cDC and NK cell subsets in an *in vivo* model of SS.

### Specific antiviral transcriptional signatures in CD1c+ cDCs from pSS patients

We analyzed the differential transcriptional patterns of CD1c+ cDCs, CD141+ cDCs and Mo from pSS patients compared to HD by RNA-seq. Principal Component Analysis of detected genes indicated that each subset from pSS was transcriptionally different from the corresponding HD controls (Supplemental Figure 5A), identifying 988, 1137 and 754 significant differentially expressed genes in Mo, CD1c+ and CD141+ cDCs, respectively (Supplemental table 2). Canonical pathway analysis by IPA predicted higher upregulation of pathways associated with IFN signaling, activation of antiviral innate responses and RIG-I in CD1c+ cDC and Mo from pSS in comparison with the CD141+ cDC (Figure 4A). Interestingly, we found overlapping and non-overlapping DEG among the top three upregulated IFN-related pathways CD1c+ cDC and Mo (Figure 4A-C). Upregulated levels of RIG-I overlapped with IRF and PKR-antiviral pathways in CD1c+ cDC and Mo from pSS. Moreover, we also observed that the Interferon Stimulated Genes (ISG) IFIT1, IFIT3, BST2, MS4A4A and STAT2 were among the non-overlapping most significantly upregulated DEG in CD1c+ cDC and conformed an unique transcriptional signature more linked to IFN I (Figure 4B-D and Supplemental Figure 5B-C). Among these, IFIT1 is a downstream effector of the RIG-I pathway, ^39^ and has been linked to expression of ligands for NK cell receptors, ^40^. Importantly, a consistent and significant increase of IFIT1 transcription was confirmed in CD1c+ cDC from pSS by qPCR (Figure 4E; Supplemental Figure 5E). In contrast, ISG DEG more significantly upregulated in Mo were associated with both IFN I and II signaling (Figure 4B; Supplemental Figure 5D). Remarkably, a weak induction on the transcription of RIG-I but a significant increase in the helicase DDX60, known as a positive modulator of RIG-I, ^41^ and in IFIT3 was validated in both CD1c+ cDC and Mo from pSS (Figure 4E; Supplemental Figure 5E). Thus, selective activation of ISGs associated to the RIG-I pathway might be more active in CD1c+ cDC.

**Figure 4.**
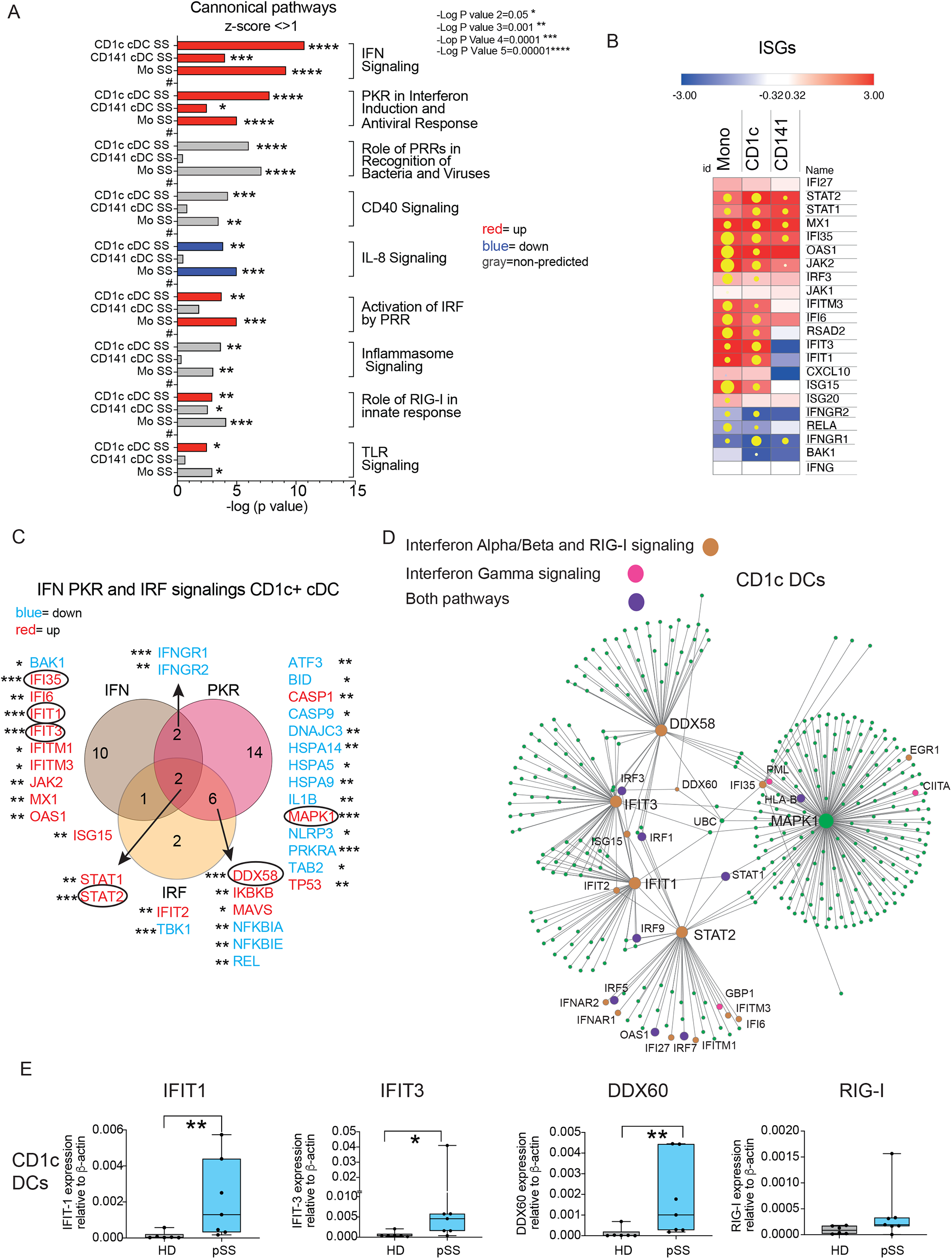
RNA-seq analysis and validation of differential transcriptional signatures in circulating Mo, CD1c+ and CD141+ cDCs from pSS patients. (A): Significance of selected upregulated (positive z-score>1; red), downregulated (negative z-score <-1; blue) canonical pathways predicted by Ingenuity Pathway Analysis (IPA) (full analysis shown in Supplemental Table 2) from significant differentially expressed genes (DEG p<0.05 after FDR correction, log2FC >1,5 and <-1,5) in Mo, CD1c+ cDCs and CD141+ cDCs from n=4 pSS vs n=4 HD. Pathways that did not have a Z-score or did not reach these mentioned value cut offs were labelled in grey. (B): Heatmaps representing log2-FC in transcription of selected Interferon Stimulated genes (ISG) or transcripts associated with the IFN pathway on each cell subset from the PB from pSS *versus* HD (red, upregulated; blue, downregulated). Yellow dots size is proportional to statistical significance of DEG expression (p<0.05 FDR corrected values). (C): Venn’s diagram of overlapping significant DEG included in selected IFN, PKR and IRF canonical pathways in PB CD1c+ cDCs from pSS patients. (D): Gene network including of DEG involved in IFNαβ and RIG-I pathways (orange), IFNγ (pink) or both Type I and II IFNs (purple) more significantly expressed in PB CD1c+ cDCs from pSS patients compared to HD. Genes characterized by higher connectivity values or assigned to the mentioned pathways are highlighted in bigger proportional sizes. (E): qPCR validation of the indicated transcript from sorted circulating CD1c+ cDCs from n=7 pSS patients (blue) versus n=6 HD (grey). Statistical significance was calculated using a two tailed U Mann Whitney test. *p<0.05; **p<0.01.

### IFN-signatures on CD1c+ cDCs from pSS patients associates with expression of ligands for NK cell receptors and enhanced NK cell maturation

We next investigated the expression of ligands for activating and inhibitory NK receptors on CD1c+ and CD141+ cDC and in Mo from pSS patients. Expression of SLAMF7 was significantly increased in all circulating myeloid subsets analyzed from pSS patients compared to HD. However, only circulating CD1c+ cDC from pSS patients expressed significantly higher levels of MICAB and ULBP1 (Figure 5A and Supplemental Figure 6A). In addition, expression of MICAB was confirmed on CD1c+ HLA-DR+ infiltrated in the SGs from pSS patients (Figure 5B). Interestingly, the expression of the inhibitory ligand PCNA tended to be lower in cDCs but increased in Mo (Supplemental Figure 6A). Therefore, we asked whether sorted circulating Mo, CD1c+ and CD141+ cDC from these patients could differ in their intrinsic abilities to stimulate autologous NK cells. Notably, CD1c+ cDC from pSS patients more efficiently increased higher proportions of CD56hi CD16+ NK cell subset enriched in PB from pSS (Figure 1B; Figure 5C). Moreover, this phenotypical maturation of NK cells was accompanied by the induction of higher proportions of CD107a+ cells (Figure 5C). Proportions of CD56dim CD16+ NK were also higher in the presence of CD1c+ cDCs and of Mo (Supplemental Figure 6B). CD141+ cDCs were unable to increase proportions of CD16+ CD56hi NK cells or CD107a+ cytotoxic cells (Figure 5C). However, they selectively induced expression of IFNg on NK cells, and an increase in the CD56dim CD16-NK subset, reduced in pSS patients (Supplemental Figure 6B-C; Figure 1B). Thus, CD1c+ cDC from pSS patients are characterized by enhanced abilities to induce maturation of cytotoxic NK cells.

**Figure 5.**
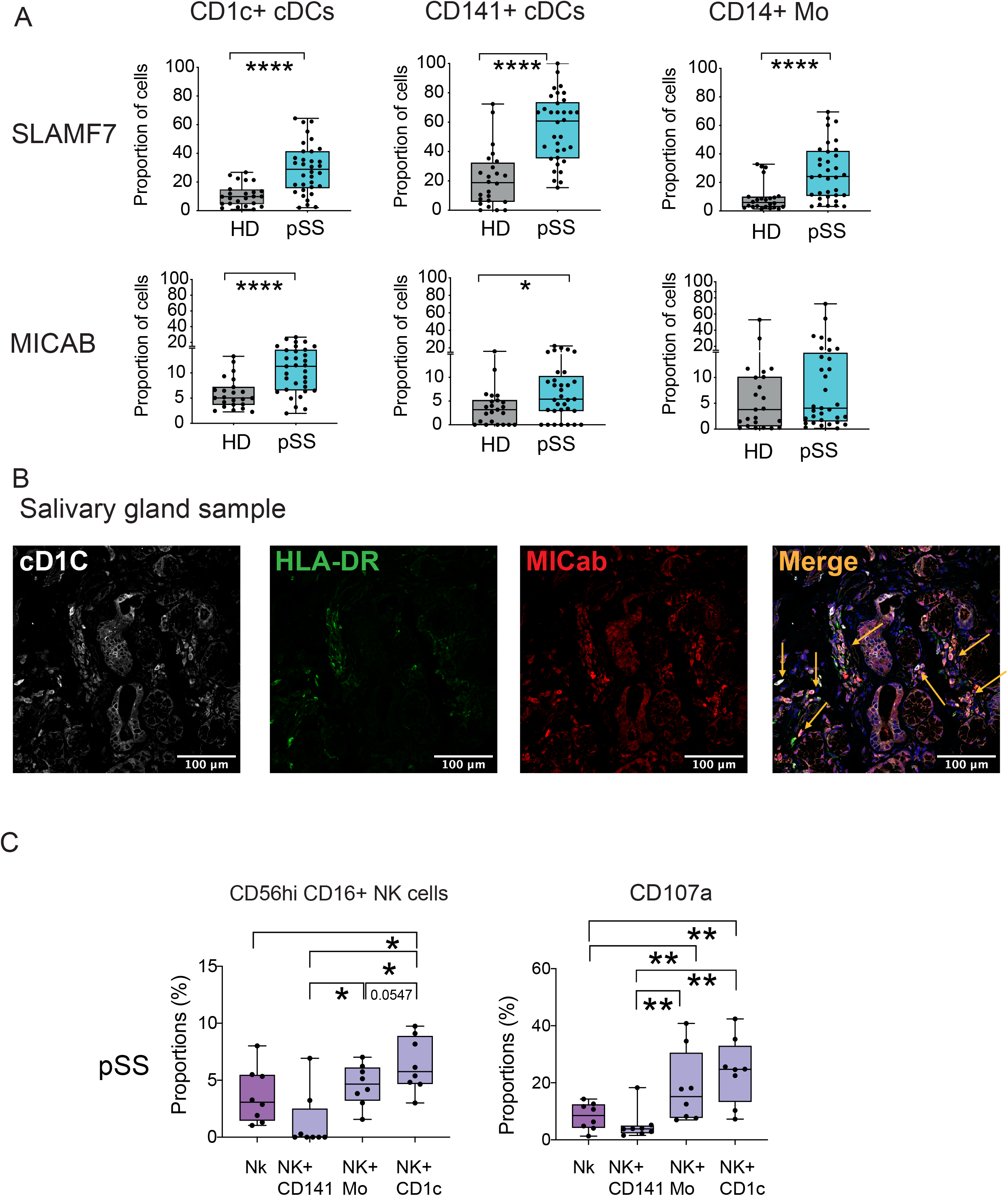
Functional NK stimulatory profile of CD1c+ cDCs, CD141 cDCs and Mo from pSS patients. (A): Proportions of SLAMF7+ and MICAB+ cells from circulating CD1c+ cDCs (left plots), CD141+ cDCs (middle plots) and CD14+ Mo (right plots) in n=24 HD (grey) and n=34 pSS patients (blue). Statistical significance was calculated using a two-tailed Mann Whitney test. *p<0.05; ****p<0.0001. (B): Representative confocal microscopy (40x magnification) showing immunofluorescence analysis of CD1c (white), HLADR (green) and MICAB (red) expression on a representative SG tissue section from a pSS patient. Cell nuclei were stained with DAPI (blue). Cells co-expressing CD1c, HLADR and MICAB are highlighted with an orange arrow. (C): Proportions of the CD56hi CD16+ NK cell subset proportions (left) after 16h culture of sorted autologous CD56+ NK cell in media alone or in the presence of sorted autologous circulating Mo, CD1c+ and CD141+ cDCs from n=8 pSS patients at ratio 1:2 (myeloid cell:NK) and expression of CD107a (right) on CD56+ NK cells in these functional assays is shown. Statistical significance was calculated using two-tailed Wilcoxon matched pairs signed rank test (*p<0.05; **p<0.01).

### RIG-I/DDX60 regulated expression of NK receptor ligands in CD1c+ cDCs

We asked whether in addition to upregulation of RIG-I and DDX60, differential expression of other innate RNA and DNA sensors could be affected in CD1c+ cDC from pSS patients compared to HD. Expression of TLR3 and TLR8, MDA5, cGAS or STING transcripts was either not affected or weakly significant in CD1c+ cDCs, (Figure 6A, Supplemental Figure 7A). In addition, we observed that CD1c+ DC and Mo from both HD and pSS induced significantly higher proportions of SLAMF7 after incubation with poly(I:C) (Figure 6B; Supplemental Figure 7B). Notably, MICAB was very significantly and more efficiently induced in CD1c+ cDCs from pSS patients than HD in response to poly(I:C) and compared with increase observed in Mo (Figure 6B, Supplemental Figure 7B). Also, no significant changes in PCNA expression were observed after poly(I:C) stimulation in these subsets (Supplemental Figure 7C). Thus, innate detection of dsRNA by CD1c+ cDC regulates the expression of ligands for activating NK receptors.

**Figure 6.**
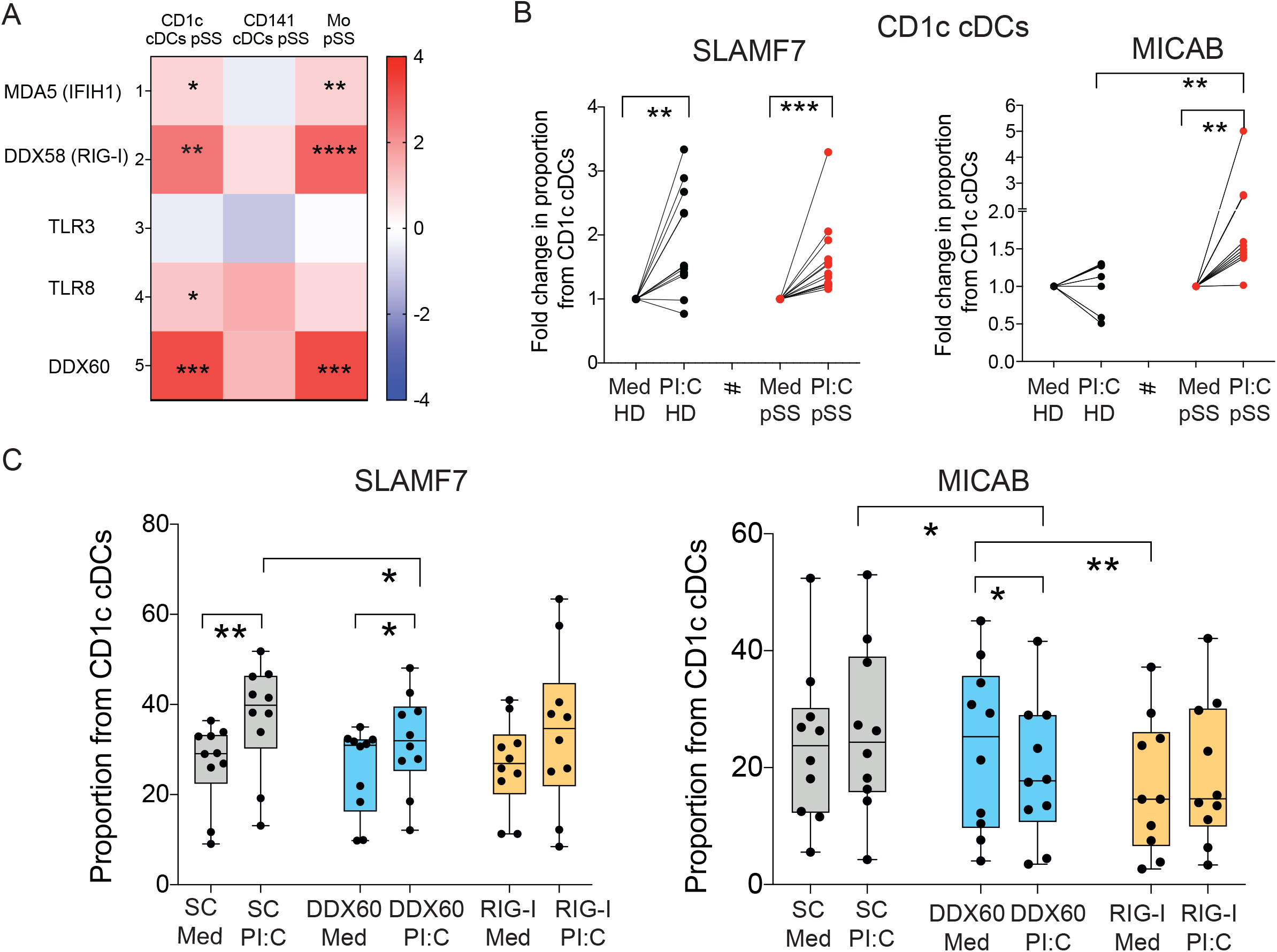
RIG-I and DDX60 are required for expression of ligands for activating NK cell receptors on CD1c+ cDCs. (A): Heatmap reflecting log2FC in transcription of innate RNA sensors in CD1c+, CD141+ cDCs and in Mo of n=4 pSS individuals compared to corresponding n=4 HD. Levels of significant comparing to HD are represent in red overexpressed and in blue repressed DEG. (B): Fold change in proportions of SLAMF7+ (left plot) and MICAB+ (right plot) cells after 16h poly(I:C) (PI:C) stimulation from gated CD1c+ cDCs in PBMC of HD (black, n=12 for SLAMF7, n=7 for MICAB) and pSS patients (red, n=14 for SLAMF7, n=9 for MICAB). Data was normalized to values in the presence of Media (Med). Statistical significance was calculated using two-tailed Wilcoxon matched pairs signed rank (*p<0.05; **p<0.01; ***p<0.001). (B): (C): Surface expression of SLAMF7 (left) and MICAB (right) on isolated primary CD1c+ cDC nucleofected with either DDX60 or RIG-I specific siRNAs or control scramble siRNA after 16h in the presence of media or poly(I:C) (PI:C). Statistical significance was calculated using two-tailed Wilcoxon matched pairs signed rank tests (*p<0.05; **p<0.01).

To determine whether this process could be dependent on expression of RIG-I or DDX60, we performed siRNA-mediated knock-down of either of these sensors or introduced control scramble siRNAs on CD1c+ cDC isolated from the PB of HD (Supplemental Figure 8A). As shown in Figure 6C, upregulation of expression of SLAMF7 was abrogated in CD1c+ cDC after DDX60 and more prominently RIG-I knock down. Moreover, expression of MICAB was significantly downregulated after stimulation with poly(I:C) or baseline in CD1c+ cDCs after both DDX60 and RIG-I knock down (Figure 6C). As expected, lower expression levels of ligands for activating NK receptors on RIG-I and DDX60 defective CD1c+ cDCs led to a partial defect in the ability to increase in the proportions of CD56dim CD16+ mature NK compared to untreated NK cells, in contrast to cells nucleofected with scramble siRNA (Supplemental Figure 8B). Together, our data indicate that expression of SLAMF7 and MICAB on CD1c+ cDC is dependent on the presence of RIG-I and DDX60 which might influence their ability to activate NK cells.

## DISCUSSION

In our study, provides new knowledge supporting increased activation of NK cells in pSS patients and their potential contribution to damage of gland tissue, ^2, 14, 42^. We characterized new biomarkers consisting in an enrichment in CD16+ CD56hi NK cells in pSS, associated with a cytotoxic phenotype and increased expression of the activating receptor NKG2D. Type I and II IFN might play opposite effects regulating NKG2D expression, ^43, 44^ but we have not directly addressed whether membrane NKG2D levels could be due to genetic or membrane trafficking alterations on pSS patients and further studies are needed to clarify these points. Alternatively, we provide novel evidence of the participation of CD1c+ cDCs as a potential cellular mechanism driving the activation of NK cells. In fact, our data indicate that CD1c+ cDC from pSS are characterized by specific IFN signatures that are associated with higher levels of the NKG2D ligand MICAB and enhanced abilities to support NK cell maturation *in vitro*. Importantly, we confirmed the infiltration of murine CD64+ CD11b+ cDCs and transitional NKG2D+ NK cells in a mouse model of pSS. Our data and previous studies in our or similar *in vivo* models, ^33, 45^ support the importance of the IFN response and the crosstalk between cDCs and NK cells during the onset of pSS, but additional studies are needed to address whether depletion of these populations impact the development of the disease. An open question that should be addressed is whether CD64 upregulation on CD1c+ cDC might be connected with the generation of Mo derived DC, which constitutively express this receptor.

We have shown that expression of DDX60, an helicase capable of potentiating RIG-I activity, is increased in CD1c+ cDC from pSS, ^41^. Supporting our findings, previous studies demonstrated that RIG-I and MDA5 can be detected *in vivo* in SGs from pSS patients ^10, 29^. In addition, we have described that IFIT1, a target gene of the RIG-I pathway, is more significantly increased in CD1c+ cDC from pSS, ^39, 46^. Importantly, IFIT1 might specifically regulate the surface expression of ligands for NK receptors on DC ^40^. We have demonstrated that induction of MICAB on CD1c+ cDC seems to be largely dependent on the expression of RIG-I and DDX60 on these cells, providing a mechanism of aberrant activation. Consistently, expression of NKG2D ligands by TLR and RNA virus stimulation has been previously reported, ^47^. Importantly, high MICAB expression has been observed in SGs from pSS, ^48^ and polymorphism of this molecule have been linked to higher susceptibility to develop the disease, ^49^. In addition, some of our data suggest that CD141+ and CD1c+ cDC might provide non-redundant signals to NK cells since we have shown that CD141+ cDC might be more effective inducing IFNg expression on NK cells or efficiently activate CD8+ T cells, as previously reported, ^26, 27, 50 5, 6^. Therefore, further studies are required to better understand complex interactions between different DC subsets contributing to pSS pathology. Despite some limitations, our study provides new cellular and mechanistic data for immunopathology of pSS, and provides CD1c+ cDC as potential new cell targets for future treatments.

## AUTHOR CONTRIBUTIONS

E.M.G., S.C., I.G.A, F.S.M., I.S.C, M.C.M and A.T.M developed the research idea and study concept, designed the study and wrote the manuscript. E.M.G., S.C. and I.G.A. supervised the study. I.S.C., M.C.M and A.T.M designed and conducted most experiments. I.S.C, D.C.F. and M.R.H. carried out in vivo experiments. C.D.A. provided technical support. I.G.A, S.C., E.R.V. and F.S.M. provided PB from study patient cohorts. S.C. A.T.M. and I.G.A. provided information of clinical parameters during the study. E.V.L provided bioinformatic support for computational RNA-Seq data analyses and supervised statistical analysis. A.A.F, R.M.V, A.D. performed and collaborated in the RNA-seq experiments. E.M.G, I.S.C and M.C.M. performed pathway analysis of transcriptional data. C.D.A, M.C.M. I.T and I.S.C. performed phenotypical analysis and sorting of samples. A.R. provided access and support for sample cell sorting. H.D.F. and F.S.M. provided qPCR reagents and cell samples from healthy donors and participated in data discussion.

## ACKNOWLEDGMENTS AND FUNDING

E.M.G. was supported by Comunidad de Madrid Talento Program (2017-T1/BMD-5396), Ramón y Cajal Program (RYC2018-024374-I), the MINECO RETOS program (RTI2018-097485-A-I00), the NIH R21 program (R21AI140930) and infectious diseases CIBER from ISCIII. C.D.A. was supported by Comunidad de Madrid Talento Program (2017-T1/BMD-5396). M.C.M was supported by the NIH (R21AI140930) and Gilead (GLD19/00168). HR17-00016 grant from “La Caixa” Banking Foundation to F.S.M and LCF/BQ/DR19/11740010 to D.C.F. A.T.M. is currently granted by a PhD fellowship from the Autonomous Region of Madrid (PEJD-2019-PRE/BMD-16851). I.G.A was supported by PEARL thanks to grants RD16/0011/0012 and PI18/0371 from the Ministerio de Economía y Competitividad (Instituto de Salud Carlos III) and co-funded by Fondo Europeo de Desarrollo Regional (FEDER). ER-V and ISC were funded by a Rio-Hortega grant CM19/00149 and CM21/00157, respectively supported by the Ministerio de Economía y Competitividad (Instituto de Salud Carlos III) and co-funded by The European Regional Development Fund (ERDF) “A way to make Europe”. S.C. has been supported by funds named AC17/00027 and FIS PI21/01474 from Instituto de Salud Carlos III, and co-funded by The European Regional Development Fund (ERDF).

