## Supplemental Data 1 for "Antiviral State of CD1c+ cDC contributes to Increased Maturation and Activation of Cytotoxic Natural Killer cells in Sjögren’s Syndrome"

**ONLINE SUPPLEMENTAL MATERIAL**

#### Supplemental Methods

*Longitudinal analysis of myeloid and NK subsets in vivo in a murine model of pSS*

Two groups of n=10 female C57BL/6 wild-type mice aged 8-10 weeks were intraperitoneally (i.p) injected either with 50 μg of poly I:C (Invivogen, San Diego, CA) or PBS in the same quantity every two days for fourteen days. Weight of animals was monitored on injection days until the endpoint. Right SGs were extracted and treated for 20 min at 37 °C with 250 μg/mL Liberase TL and 100 μg/mL DNAase I (Roche, Basel, Switzerland) in HBSS medium to facilitate tissue enzymatic digestion and cell disaggregation. Single-cell suspensions were then obtained by grating the digested organs through a 70μm cell strainer (Falcon). Part of these cells were directly used for phenotypical characterization of myeloid and NK cell subsets by flow cytometry, while others were cultured in the presence of phorbol 12-myristate 13-acetate (50 ng/mL) and ionomycin (1 μg/mL; Sigma-Aldrich, San Luis, MO) and incubated at 1h at 37ºC in 5% CO2 and then another 4h in the presence of 5 μg/mL Brefeldin A (Sigma-Aldrich), Monesin (Sigma-Aldrich) and an anti-CD107a antibody (BioLegend). In all cases, cells were firstly incubated with anti-FcRII/III and LIVE/DEAD Fixable Yellow Dead Cell Stain (Invitrogen, Waltham, MA) and then stained with specific antibodies (Supplemental Table 3). The Fixation/Permeabilization Solution Kit (BD Biosciences, San Jose, CA) was used for intracellular staining. Trucount Absolute Counting Tubes (BD Biosciences, San Jose, CA) were added to obtain and homogenize absolute cell counts. Samples were acquired in a LSRFortessa and FACSCanto Flow Cytometer (BD Biosciences, San Jose, CA) and analyzed using the FlowJo software (Tree Star). Results were corroborated in two independent experiments.

Analysis of proportions of hematopoietic CD45.2+ cells or absolute numbers in the SG of Ly6C+ Monocytes, Ly6C- MCH-II+ CD11cHi CD11b+ XCR1- (homologue to human CD1c+ cDC), Ly6C- MCH-II+ CD11cHi CD11b- XCR1+ (homologue to human CD141+ cDC) cDC subsets and CD3- NK1.1+ NK subsets defined by CD11b versus CD27 expression within the infiltrate were longitudinally analyzed at 8 and 14 days after treatment initiation by sacrifying n=5 mice per group at each timepoint. Levels of the activation/maturation marker CD64 were analyzed on each gated myeloid population. In addition, levels of NKG2D were determined on NK cell subsets. Left SGs were preserved in paraffin for subsequent histological eosin/hematoxylin staining.

#### SUPPLEMENTAL FIGURE LEGENDS

**Supplemental Figure 1. Phenotypic analysis of natural killer cell populations from pSS patients**. (A): Summarized proportions of circulating CD56- CD16+ (left panel), CD56dim CD16+ (middle plot) and CD56hi CD16- NK subsets (right panel) in 25 healthy donors (HD; grey) and 34 primary Sjögren´s syndrome (pSS; blue) patients. (B-C): Proportions of cells expressing CD107a (B), and frequencies of total IFNγ+, total TNFα+, IFNγ+ CD107a+ and TNFα+ CD107a+ cells (C) within the indicated circulating NK cell subsets from 16 HD (grey) and 25 pSS patients (blue). Statistical significance was calculated using a two-tailed Mann Whitney test. *p<0.05; **p<0.01. (D): Eosin/hemotoxilin tissue sections from SG tissue of a representative patient to different magnifications 20x (left) and 40x (right), showing low (black stars) and highly (red stars) infiltrated areas with altered morphology. (E): Image J quantification of proportions of CD56+ NK cells co-expressing Granzyme B on the mentioned high and low infiltrated areas.

**Supplemental Figure 2. Characterization of activating and inhibitory receptors and functional analysis of NK cells from pSS patients.** (A): Proportions of NKG2D (left plot), and SLAMF7 (right plot) activating receptors and the inhibitory NK receptor NKG2A (left middle plot) in all NK cell subsets from primary Sjögren´s syndrome (pSS; blue) and healthy donors (HD; grey). (B): Flow cytometry gating strategy representing frequencies of dead K562-GFP after culture in medium alone or in the presence of isolated NK cells. K562 cells were identified as Violet Trace to exclude NK cells. Dead K562 were quantified as cells losing GFP expression and acquiring of staining of the viability dye marker. Statistical significance was calculated using a two-tailed Mann Whitney test. *p<0.05; **p<0.01; ***p<0.001; ****p<0.0001.

**Supplemental Figure 3. Identification and frequencies of circulating myeloid subsets from pSS patients.** (A): Flow cytometry gating strategy identifying CD14+ Monocytes and HLA-DR+ CD11c+ conventional (cDC) subdivided into CD1c+ and CD141+ cDCs and CD11c- CD123+ plasmacytoid (pDC) dendritic cells from Lineage (CD3, CD19, CD20, CD56) negative CD14- cells. CD14+ Mo was further defined as CD14lo CD16hi non classical (NC), CD14+ CD16+ transitional (T) and CD14+ CD16- classical (C) Mo on the basis of the expression of CD16. (B-D): Box and whisker plots representing frequencies of Lin- HLA-DR+ CD11c- CD123+ pDCs (B), T Mo and C Mo subsets in live mononuclear cells from PB from HD (grey, n=27) and pSS patients (blue, n=34) and Mean Fluorescence Intensity (MFI) of expression of CD64 on these subsets (D) obtained by flow cytometry. Statistical significance was calculated using a two-tailed Mann Whitney test. *p<0.05; **p<0.01; ****p<0.0001.

**Supplemental Figure 4. *In vivo* analysis of cDC, NK cells, and Mo in a mouse model of Sjögren´s syndrome.** (A): Schematic representation of the experimental design for the *in vivo* experiments where myeloid and NK cell subsets were studied in the submandibular salivary gland (SMSG) after the injection of mice with poly I:C or PBS. (B): Frequencies (left) and absolute numbers (right) of hematopoietic CD45.2+ cells represented detected on homogenized SMSG from the mentioned mice groups. (C): Representative flow cytometry gating strategy defining cDC and NK subsets in the SMSG and the expression of the indicated markers on these cells. (D,F): Frequencies of XCR1+ CD11chi cDC, Ly6+ Mo and CD11cint cells (D) or the CD11blo CD27+ NK cell subset (F) in the SMSG of mice injected with PBS (blue) or poly I:C (red) at 8 and 14 days after treatment initiation. (E): Proportions of CD64+ cells included in XCR1+ cDC and Ly6C+ Mo in the SMSG from the two mice groups at the indicated timepoints are also shown. (G): Proportions of circulating CD11b+ CD27+ NK cell subset and expression of NKG2D in this subset in both groups of mice at day 8 and 14 post treatment. (H): Proportions of the CD107a+ IFNγ+ cells on the indicated gated NK cell subsets present in the SMSG and PB of the indicated experimental groups at 8 days after treatment initiation. These data are representative from one out of two independent experiments. Statistical significance was calculated with a 2way ANOVA test. *p<0.05; **p<0.01; ***p<0.001; ****p<0.0001.

**Supplemental Figure 5. Pathway analysis of differential transcriptional patterns of Mo and cDC subsets from the blood of pSS patients.** (A): Principal Component Analysis (PCA) representing transcriptional profiles in CD1c cDC, CD141+ cDC and Mo from the blood of n=4 primary Sjögren´s syndrome (pSS; green) and n=4 healthy donors (HD; red). (B): Heatmap reflecting Log2FC in expression of 47 transcripts associated with IFN signaling or IFN Stimulated Genes (ISG) on Mo, CD1c+ and CD141+ cDCs from the blood of 4 pSS patients compared to 4 HD. Highlighted Red show upregulated expression and blue, show downregulated pathways. Yellow dots size is proportional to statistical significance of DEG expression (p<0.05 FDR corrected values). (C): Venn´s diagram of overlapping significant

DEG included in selected IFN, PKR and IRF canonical pathways in PB Mo from pSS individuals. (D): Gene network including of DEG involved in IFNαβ and RIG-I (orange), IFNγ (pink) or both Type I and II IFNs (purple) signaling more significantly expressed in PB Mo from pSS patients compared to HD. (E): qPCR validation of the indicated transcript from sorted circulating Mo from n=7 pSS patients (blue) versus n=5 HD (grey). Statistical significance was calculated using a two tailed U Mann Whitney test. *p<0.05; **p<0.01.

**Supplemental Figure 6. Functional ability of CD1c+ cDCs, CD141 cDCs and Mo to stimulate NK cells** (A): Proportions of ULBP1+ (upper plots) and PCNA+ (bottom plots) cells from circulating CD1c+ cDCs (left plots), CD141+ cDCs (middle plots) and CD14+ Mo (right plots) in 24 healthy donors (HD; grey) and 34 primary Sjögren´s syndrome (pSS; blue) individuals. Statistical significance was calculated using a two-tailed Mann Whitney test. *p<0.05. (B): Proportions of the CD56dim CD16+ (left) and CD56dim CD16- (right) NK cell subsets from pSS patients, after 16h culture of sorted autologous CD56+ NK cell in media alone or in the presence of sorted autologous circulating Mo, CD1c+ and CD141+ cDCs from n=8 pSS patients at ratio 1:2 (myeloid cell:NK). (D): Proportions of IFNγ+ NK cells from pSS patients in these functional assays is shown. Statistical significance was calculated using two-tailed Wilcoxon matched pairs signed rank test (*p<0.05; **p<0.01).

**Supplementary Figure 7. Implication of innate sensing of RNA in the expression of ligands for NK cell receptors in circulating Mo and cDC from Sjögren patients**. (A): Heatmap reflecting fold change in transcriptional expression of the indicated intracellular RNA sensors in CD1c+ cDC, CD141+ cDC and Mo from pSS compared to HD. Significant changes are highlighted. Red and blue indicate upregulated and downregulated, expression respectively.

(B): Fold change in proportions of cells expressing SLAMF7 (left) and MICAB (right) in Mo (B) and PCNA (C) ligands in CD1c+ cDC and Mo from the blood of HD (black; n=12 SLAMF7 and for PCNA; n=7 for MICAB) or pSS individuals (red; n=14 SLAMF7 and for PCNA; n=9 for MICAB) after 16h of culture in the presence of poly I:C (PI:C) normalized to values present in cells cultured in media only (Med).

**Supplemental Figure 8. Impact of si-RNA-mediated knock down of DDX60 and RIG-I on CD1c+ cDCs.** (A): Fold change in transcriptional levels of DDX60 (left) and DDX58 (RIG-I, right) analyzed by RT-qPCR and normalized by endogenous β-Actin levels in isolated CD1c+ cDC nucleofected with specific siRNAs. RT-qPCR values were normalized to those in cells nucleofected with control scramble (SC) siRNAs. (B): Proportions of CD56dim CD16+ (left plot) NK cells after culture in media alone or in the presence of CD1c+ cDC nucleofected with the indicated control SC siRNA or DDX60 or RIG-I-specific siRNAs. Statistical significance was calculated using two-tailed Wilcoxon matched pairs signed rank tests (*p<0.05; **p<0.01; ***p<0.001).

Supplemental Figure 1

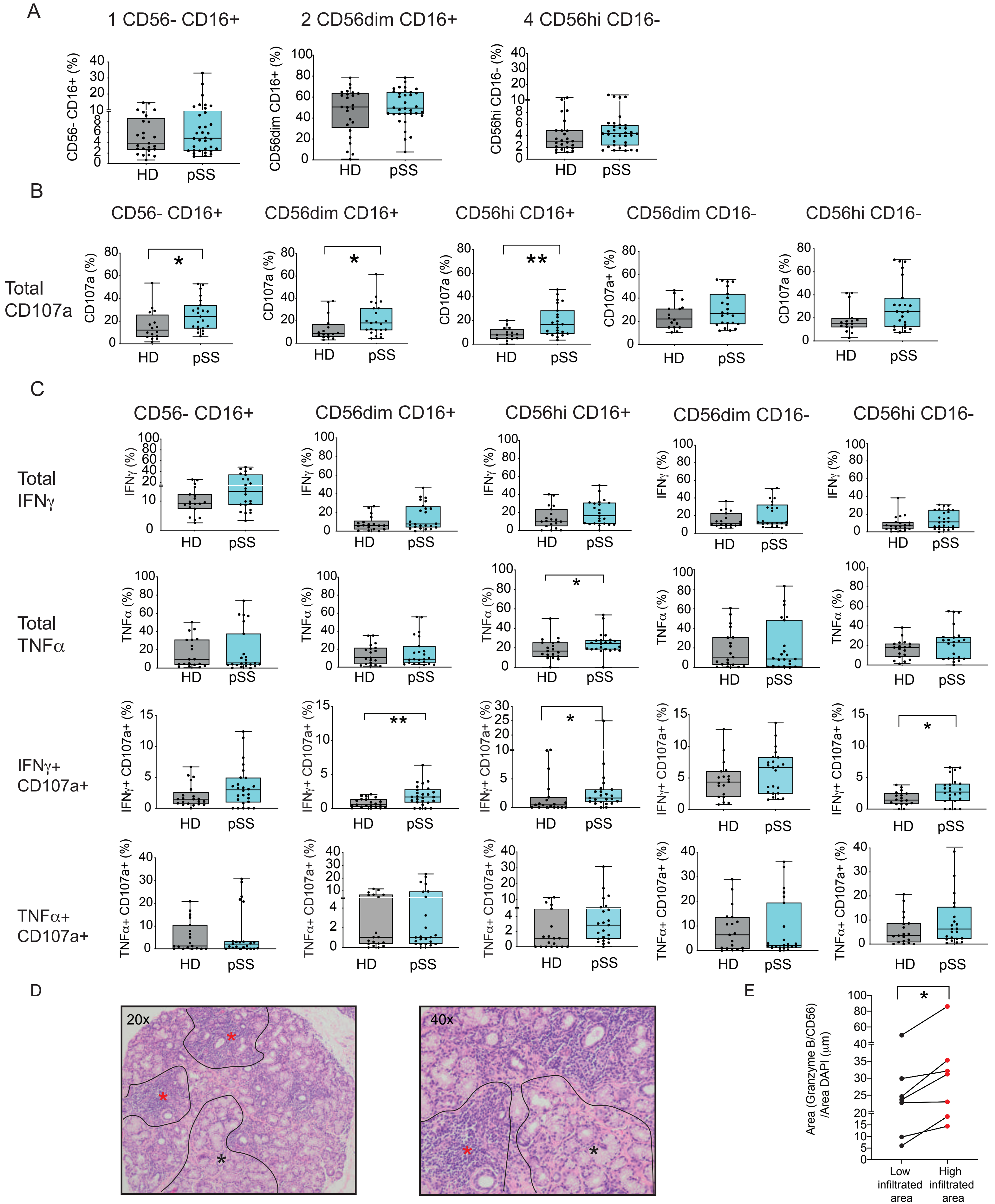

Supplemental Figure 2

A

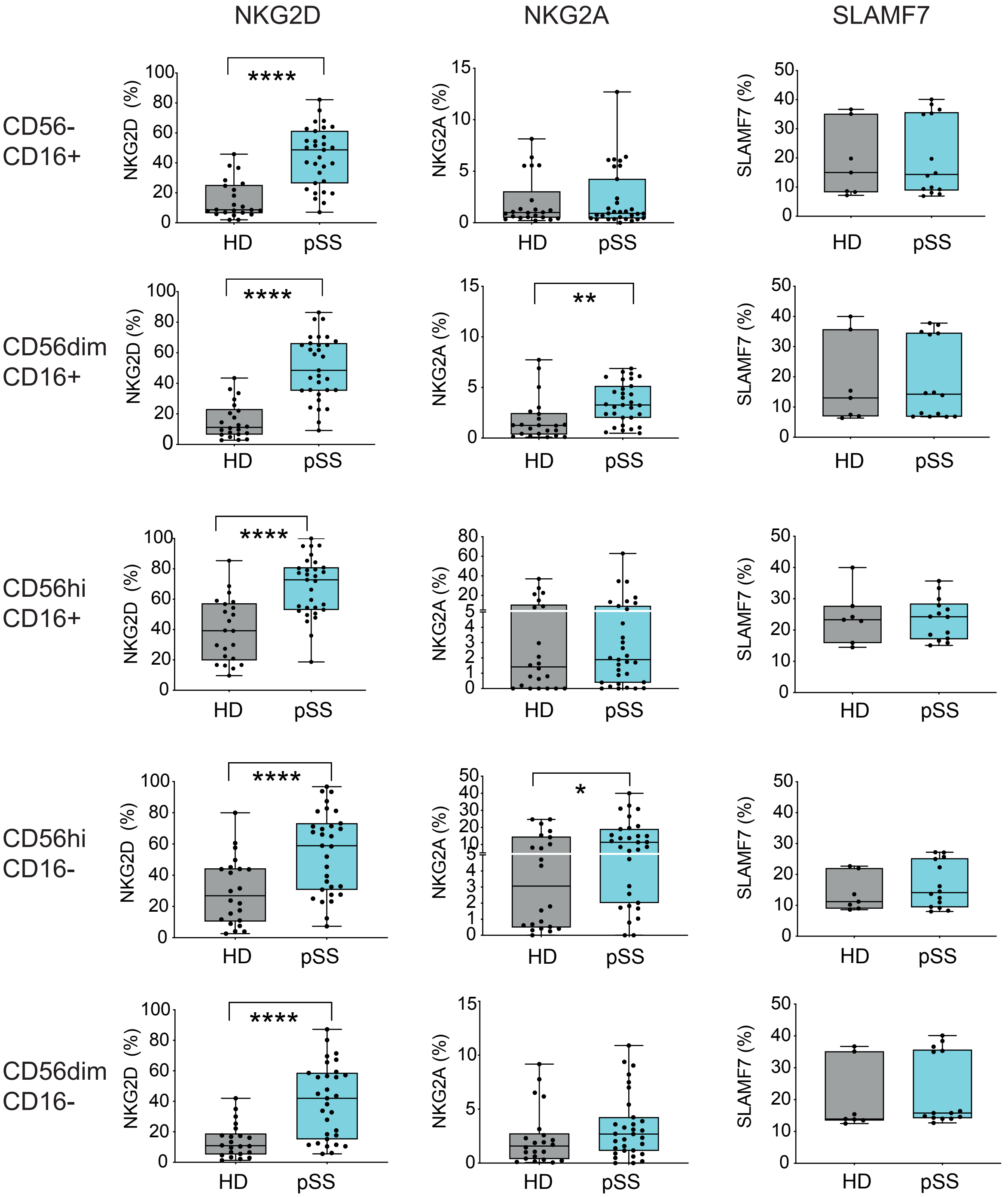


B

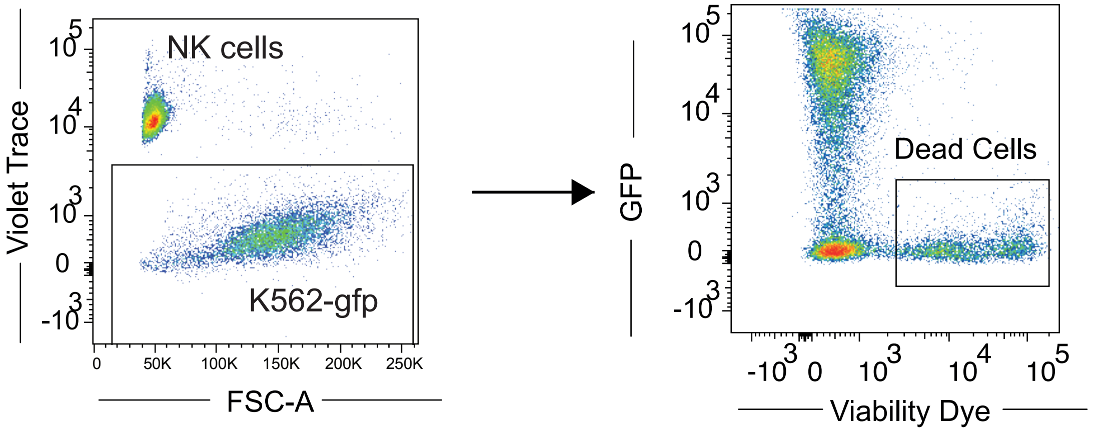

Suplemmental Figure 3

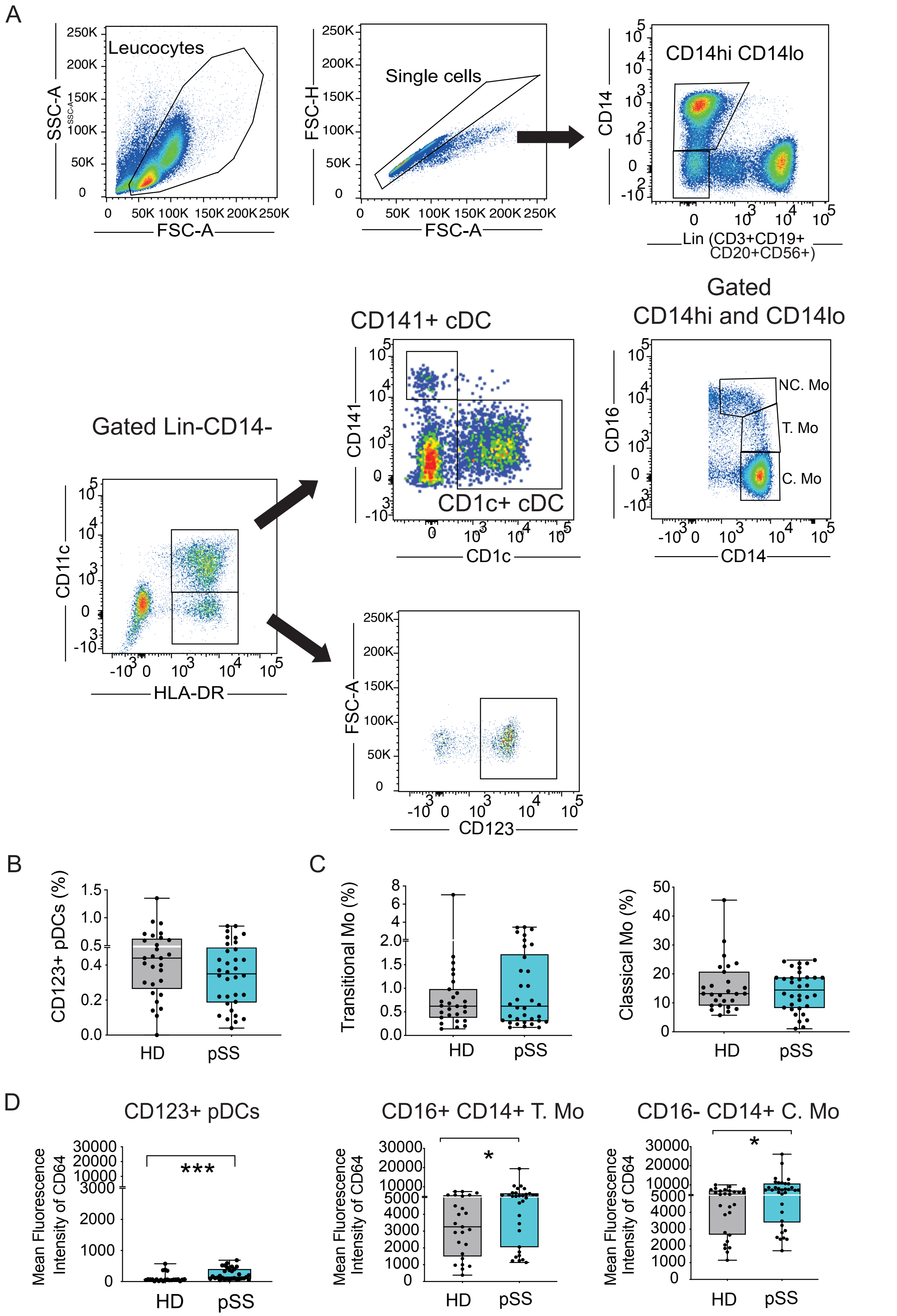

Suplemental Figure 4

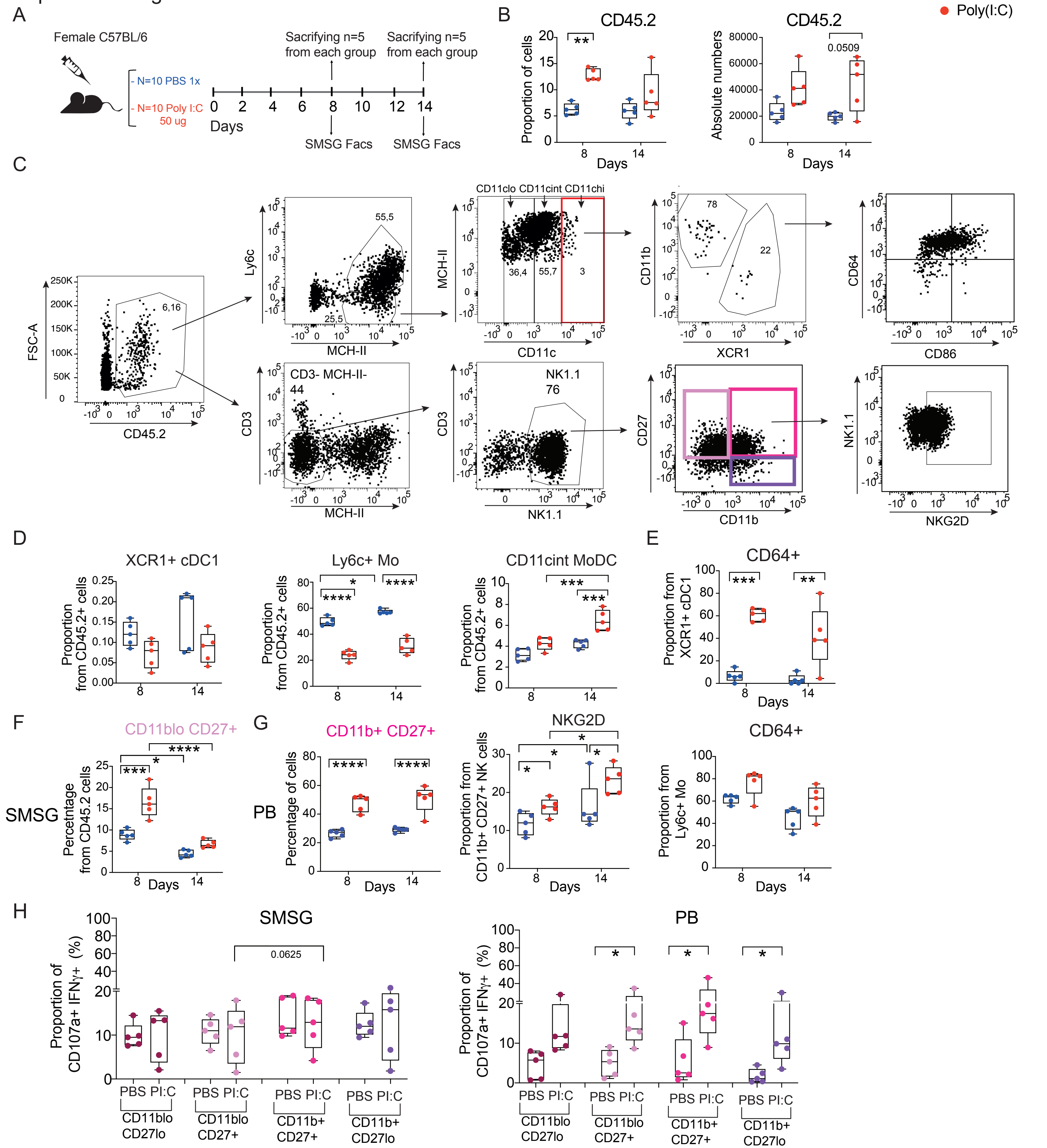

Suplemmental Figure 5

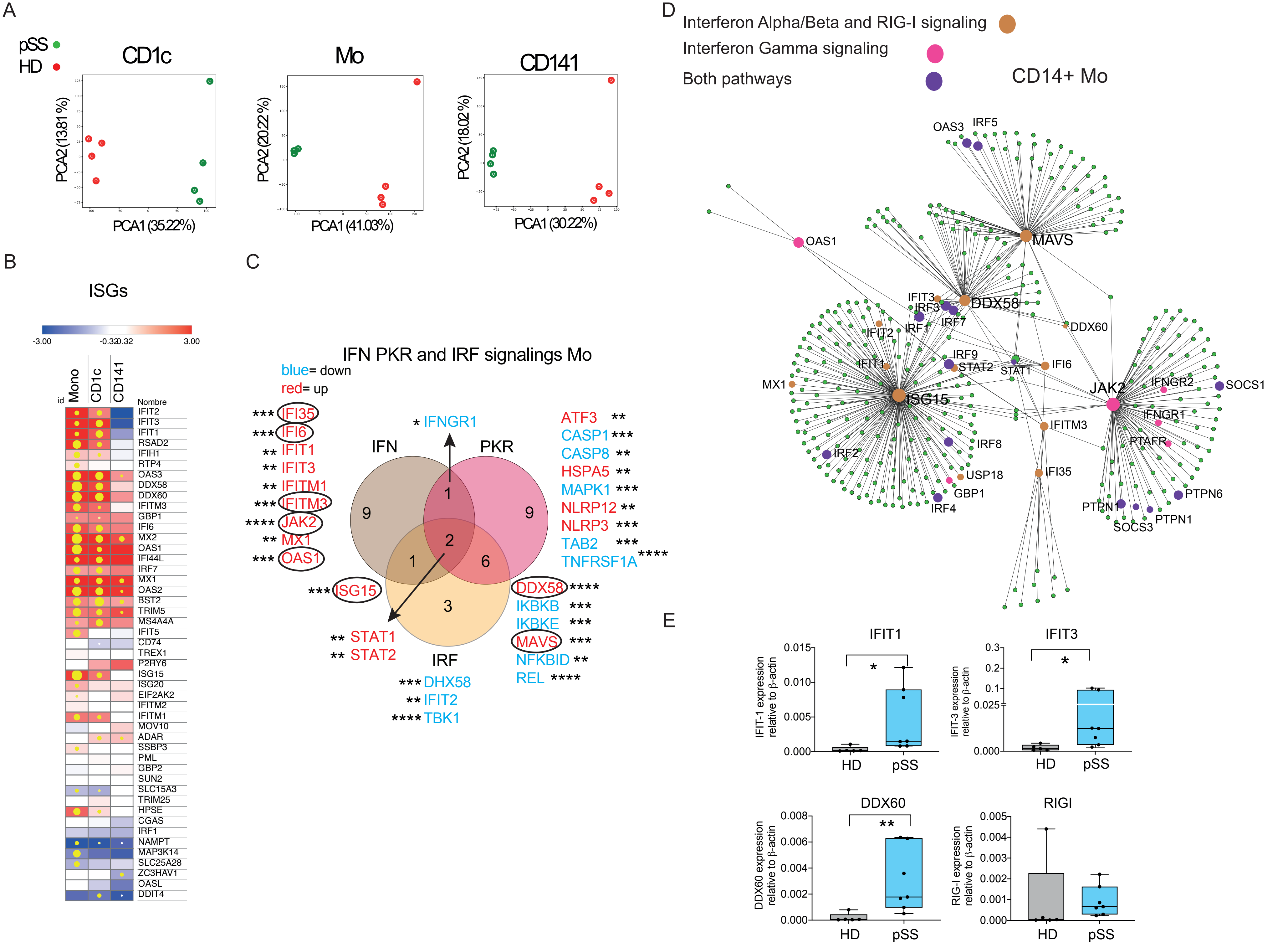

Supplemental Figure 6

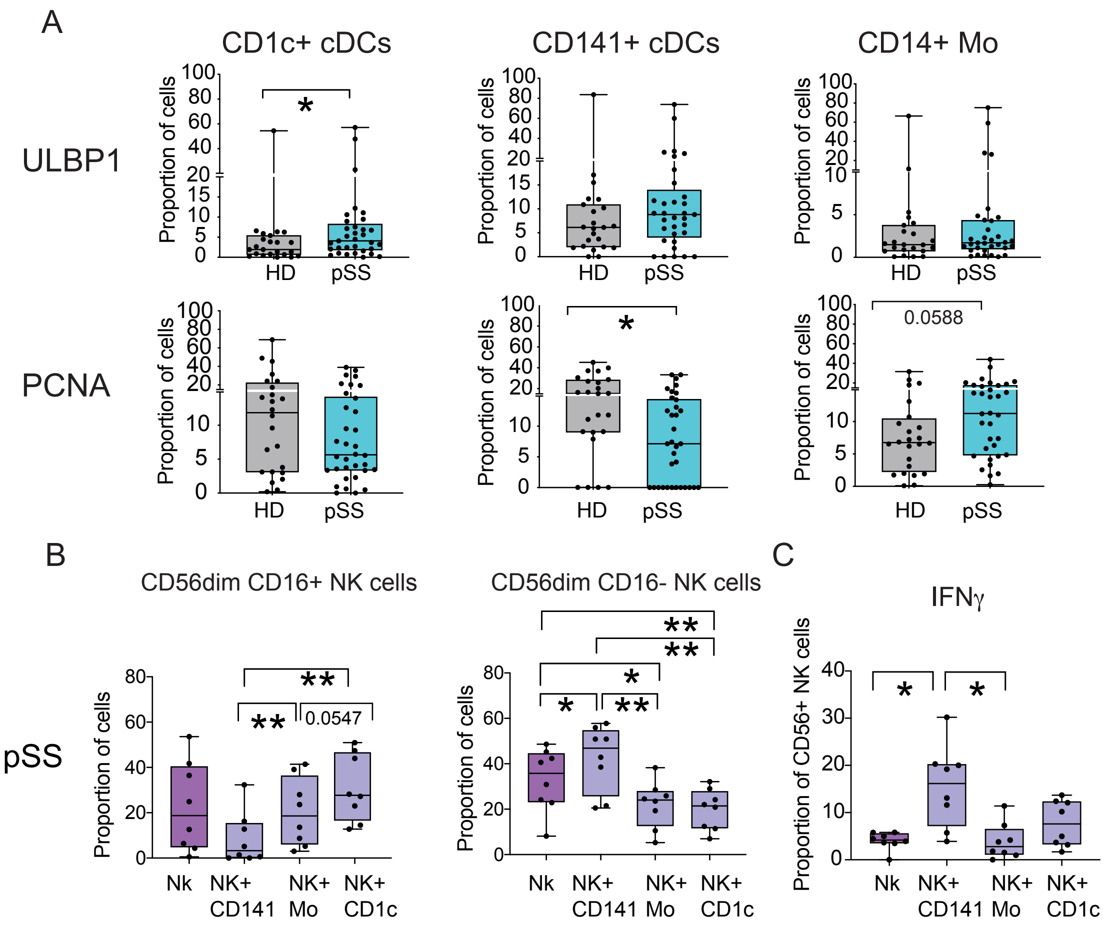

### Supplemental Figure 7

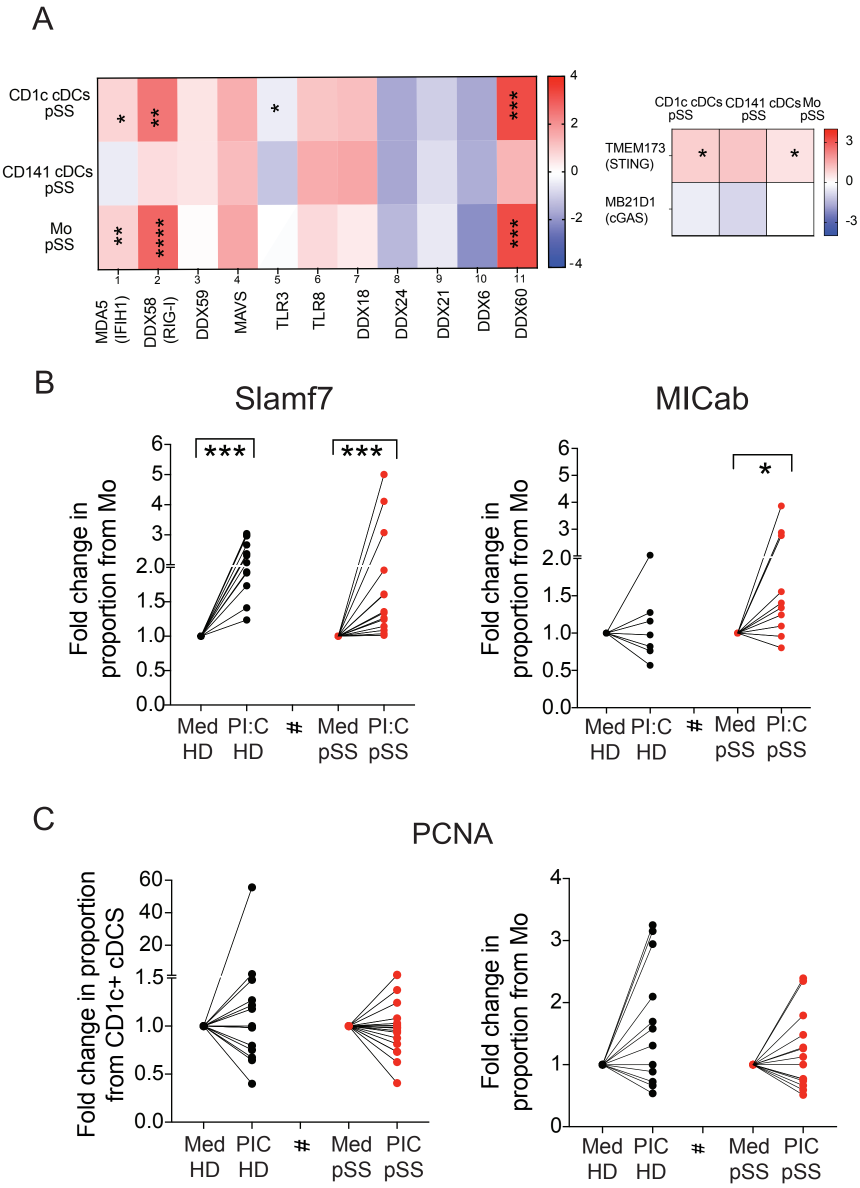

### Supplemental Figure 8

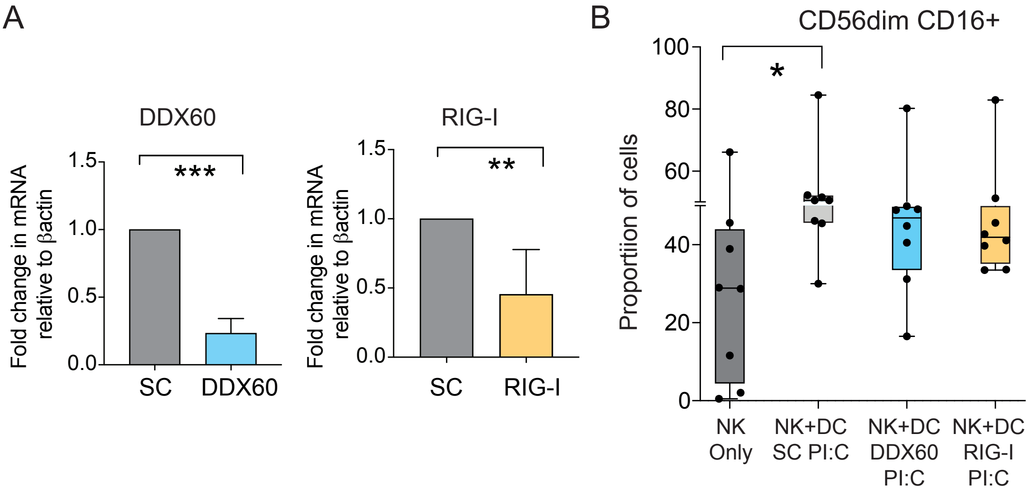

**Supplemental Table 1. Clinical characteristics of Sjögren´s syndrome patients**

| | pSS |
|---|---|
| **Number of patients, (n)** | **38** |
| Median Age *(years, Min-Max)* | 64 (23-88) |
| Sex (% Female) | 98% |
| **Severity, n (%)** | |
| Mild | 21 (56%) |
| Severe | 10 (26%) |
| Treated (GC, MTX or/and MA) | 7 (18%) |
| **Extra-glandularmanifestations, n (%)** | |
| Arthritis | 13 (34.2%) |
| Raynaud | 2 (5.3%) |
| Oral ulcers | 1 (2.6%) |
| Vasculitis | 2 (5.26%) |
| Polyneuropathy | 1 (2.6%) |
| Pulmonary affection | 1 (2.6%) |
| Kidney affection | 1 (2.6%) |
| Lymphoma | 2 (5.26%) |
| **Laboratory values** ***(Median, Min-Max)*** | |
| RF (UI/ml) | 24 (8-209) |
| ESR | 26 (3-79) |
| CRP (mg/dl) | 0.16 (0.03-1.7) |
| Lymphocyte count/mm$^3$ | 1545 (490-5190) |
| Gammaglobulin (g/L) | 1.43 (0.41-2.79) |
| IgG (mg/dL) | 1680 (591-3430) |
| C3 (mg/dL) | 105 (78.9-165) |
| C4 (mg/dL) | 20 (11-29) |
| **Treatment, n (%)** | |
| Hidroxychloroquine | 14 (36.8%) |
| Glucocorticoids (GC) | 4 (10.5%) |
| Methotrexate (MTX) | 4 (10.5%) |
| Monoclonal Antibodies (MA) | 3 (7.9%) |

Abbreviations: RF: rheumatoid factor; ESR: erythrocyte sedimentation rate; CRP: C-reactive protein; IgG: immunoglobulin type G; C3: fraction 3 of complement; C4: fraction 4 of complement; n: number.

**Supplemental table 2: DEGs on CD1c+ cDCs, CD141+ cDCs and Mo (Log2 FC cutoff upregulated >1.5; downregulated <-1.5)**

#### DEG CD1c+ cDCs pSS *versus* HD

| Transcipt ID | Gene name | Log2(FC) | FDR < 0.05 |
|---|---|---|---|
| ENSG00000137959 | IFI44L | 4.16535789 | 0.00167388 |
| ENSG00000111335 | OAS2 | 4.15678392 | 0.00068553 |
| ENSG00000182578 | CSF1R | 3.93436922 | 0.00186519 |
| ENSG00000168329 | CX3CR1 | 3.83812944 | 0.01267943 |
| ENSG00000177409 | SAMD9L | 3.76415944 | 0.00072853 |
| ENSG00000157601 | MX1 | 3.7111664 | 0.00190217 |
| ENSG00000089127 | OAS1 | 3.52032768 | 0.00433679 |
| ENSG00000111913 | RIPOR2 | 3.51241416 | 0.0040364 |
| ENSG00000121807 | CCR2 | 3.51168175 | 0.02587417 |
| ENSG00000132530 | XAF1 | 3.48286431 | 0.00089939 |
| ENSG00000101347 | SAMHD1 | 3.45232655 | 0.00847993 |
| ENSG00000249437 | NAIP | 3.42167956 | 0.04887038 |
| ENSG00000111331 | OAS3 | 3.41018922 | 0.00057232 |
| ENSG00000133706 | LARS | 3.38958469 | 0.0026949 |
| ENSG00000183486 | MX2 | 3.32443198 | 0.00080111 |
| ENSG00000140749 | IGSF6 | 3.31354682 | 0.0026949 |
| ENSG00000137628 | DDX60 | 3.2680394 | 0.00045076 |
| ENSG00000110876 | SELPLG | 3.24737994 | 0.00155933 |
| ENSG00000139970 | RTN1 | 3.22749388 | 0.00272842 |
| ENSG00000092964 | DPYSL2 | 3.14283953 | 0.00253643 |
| ENSG00000163683 | SMIM14 | 3.14167309 | 0.00668554 |
| ENSG00000170581 | STAT2 | 3.13212829 | 0.00080302 |
| ENSG00000005020 | SKAP2 | 3.1231954 | 0.00186707 |
| ENSG00000088827 | SIGLEC1 | 3.04532683 | 0.01503232 |
| ENSG00000138449 | SLC40A1 | 2.95156286 | 0.00108396 |
| ENSG00000088986 | DYNLL1 | 2.93772619 | 0.00233277 |
| ENSG00000158488 | CD1E | 2.89042691 | 0.00562529 |
| ENSG00000110077 | MS4A6A | 2.86675942 | 0.01645689 |
| ENSG00000082074 | FYB1 | 2.8576699 | 0.00513019 |
| ENSG00000133835 | HSD17B4 | 2.82579063 | 0.00365669 |
| ENSG00000240065 | PSMB9 | 2.81532891 | 0.00096838 |
| ENSG00000119917 | IFIT3 | 2.81308894 | 0.00098988 |
| ENSG00000117676 | RPS6KA1 | 2.79970312 | 0.00089939 |
| ENSG00000178927 | C17orf62 | 2.79591345 | 0.00088661 |
| ENSG00000038427 | VCAN | 2.76743251 | 0.00536505 |
| ENSG00000167220 | HDHD2 | 2.74889964 | 0.00261709 |

| | | | |
|---|---|---|---|
| ENSG00000006715 | VPS41 | 2.73934221 | 0.00068553 |
| ENSG00000107551 | RASSF4 | 2.73092693 | 0.00329946 |
| ENSG00000093072 | ADA2 | 2.72385816 | 0.00251682 |
| ENSG00000165168 | CYBB | 2.70586786 | 0.00264376 |
| ENSG00000181381 | DDX60L | 2.69717824 | 0.00052149 |
| ENSG00000148110 | MFSD14B | 2.63197719 | 0.00520463 |
| ENSG00000196975 | ANXA4 | 2.61333737 | 0.00077204 |
| ENSG00000197713 | RPE | 2.60713676 | 0.01153698 |
| ENSG00000179639 | FCER1A | 2.59644663 | 0.00882281 |
| ENSG00000175567 | UCP2 | 2.59370025 | 0.00563712 |
| ENSG00000197043 | ANXA6 | 2.58953409 | 0.00467602 |
| ENSG00000135218 | CD36 | 2.58500504 | 0.00349422 |
| ENSG00000134326 | CMPK2 | 2.56634923 | 0.00203231 |
| ENSG00000133574 | GIMAP4 | 2.5651703 | 0.02150282 |
| ENSG00000110031 | LPXN | 2.55736628 | 0.00563329 |
| ENSG00000188641 | DPYD | 2.53918064 | 0.00935057 |
| ENSG00000169385 | RNASE2 | 2.53792206 | 0.00498953 |
| ENSG00000196209 | SIRPB2 | 2.52111187 | 0.01585679 |
| ENSG00000107201 | DDX58 | 2.51544972 | 0.00080302 |
| ENSG00000105281 | SLC1A5 | 2.49745673 | 0.00099236 |
| ENSG00000133943 | DGLUCY | 2.49512886 | 0.00155927 |
| ENSG00000035403 | VCL | 2.47427384 | 0.0012446 |
| ENSG00000170915 | PAQR8 | 2.4699681 | 0.00200912 |
| ENSG00000197142 | ACSL5 | 2.46991698 | 0.00362829 |
| ENSG00000161929 | SCIMP | 2.45193688 | 0.00709936 |
| ENSG00000105483 | CARD8 | 2.43912494 | 0.00089939 |
| ENSG00000154930 | ACSS1 | 2.43622234 | 0.00230358 |
| ENSG00000138119 | MYOF | 2.4320701 | 0.0040802 |
| ENSG00000165795 | NDRG2 | 2.40671028 | 0.00744894 |
| ENSG00000177575 | CD163 | 2.39869809 | 0.00650305 |
| ENSG00000146192 | FGD2 | 2.39240703 | 0.00246913 |
| ENSG00000075303 | SLC25A40 | 2.39230879 | 0.00230358 |
| ENSG00000050344 | NFE2L3 | 2.39130111 | 0.00292999 |
| ENSG00000163220 | S100A9 | 2.38949301 | 0.02885702 |
| ENSG00000165672 | PRDX3 | 2.38019003 | 0.00142465 |
| ENSG00000121281 | ADCY7 | 2.37163873 | 0.00167388 |
| ENSG00000136631 | VPS45 | 2.37092946 | 0.00286187 |
| ENSG00000188554 | NBR1 | 2.36778573 | 0.0012446 |
| ENSG00000134986 | NREP | 2.35723992 | 0.00237378 |
| ENSG00000106785 | TRIM14 | 2.3368806 | 0.00576925 |
| ENSG00000084733 | RAB10 | 2.32913481 | 0.00108396 |
| ENSG00000184432 | COPB2 | 2.32693742 | 0.00467602 |

| | | | |
|---|---|---|---|
| ENSG00000169413 | RNASE6 | 2.32692653 | 0.0010458 |
| ENSG00000151702 | FLI1 | 2.31890751 | 0.00081022 |
| ENSG00000104974 | LILRA1 | 2.29125844 | 0.00474195 |
| ENSG00000127951 | FGL2 | 2.2824011 | 0.00089939 |
| ENSG00000166801 | FAM111A | 2.26969444 | 0.00208192 |
| ENSG00000111269 | CREBL2 | 2.26874304 | 0.00458456 |
| ENSG00000164414 | SLC35A1 | 2.26564961 | 0.0022793 |
| ENSG00000170458 | CD14 | 2.25623821 | 0.01184035 |
| ENSG00000185745 | IFIT1 | 2.2531576 | 0.00089939 |
| ENSG00000068079 | IFI35 | 2.25305844 | 0.00098988 |
| ENSG00000258659 | TRIM34 | 2.24478854 | 0.00339437 |
| ENSG00000164125 | FAM198B | 2.24095776 | 0.01271911 |
| ENSG00000065413 | ANKRD44 | 2.23995368 | 0.00068553 |
| ENSG00000138459 | SLC35A5 | 2.23385815 | 0.00360081 |
| ENSG00000204131 | NHSL2 | 2.23097905 | 0.00406198 |
| ENSG00000125703 | ATG4C | 2.23095256 | 0.0047099 |
| ENSG00000107929 | LARP4B | 2.22788083 | 0.00358122 |
| ENSG00000178685 | PARP10 | 2.22287319 | 0.00015003 |
| ENSG00000155097 | ATP6V1C1 | 2.21598765 | 0.00433873 |
| ENSG00000186088 | GSAP | 2.21576679 | 0.00261709 |
| ENSG00000082996 | RNF13 | 2.21087655 | 0.00073864 |
| ENSG00000072501 | SMC1A | 2.20979185 | 0.00376598 |
| ENSG00000096968 | JAK2 | 2.20812913 | 0.00130443 |
| ENSG00000198771 | RCSD1 | 2.20709862 | 0.00457885 |
| ENSG00000145781 | COMMD10 | 2.20515315 | 0.00155034 |
| ENSG00000136485 | DCAF7 | 2.19735281 | 0.01586099 |
| ENSG00000004455 | AK2 | 2.19351353 | 0.0008952 |
| ENSG00000115415 | STAT1 | 2.17610829 | 0.00211531 |
| ENSG00000102699 | PARP4 | 2.17444696 | 0.00340571 |
| ENSG00000025708 | TYMP | 2.17024743 | 0.00563582 |
| ENSG00000103381 | CPPED1 | 2.16814525 | 0.00150263 |
| ENSG00000090861 | AARS | 2.16649736 | 0.00043805 |
| ENSG00000093144 | ECHDC1 | 2.16126628 | 0.00419863 |
| ENSG00000143624 | INTS3 | 2.15695705 | 0.01343737 |
| ENSG00000113845 | TIMMDC1 | 2.15354406 | 0.00720168 |
| ENSG00000168310 | IRF2 | 2.15086059 | 0.0041688 |
| ENSG00000103313 | MEFV | 2.14612293 | 0.00418017 |
| ENSG00000102893 | PHKB | 2.14216041 | 0.00131723 |
| ENSG00000138413 | IDH1 | 2.14150089 | 0.00290205 |
| ENSG00000160593 | JAML | 2.14039823 | 0.00460845 |
| ENSG00000145416 | MARCH1 | 2.13486128 | 0.00931242 |
| ENSG00000133106 | EPSTI1 | 2.13206205 | 0.00068553 |

| | | | |
|---|---|---|---|
| ENSG00000159228 | CBR1 | 2.13014578 | 0.00045076 |
| ENSG00000116455 | WDR77 | 2.12930733 | 0.00358122 |
| ENSG00000137965 | IFI44 | 2.12777848 | 0.00876064 |
| ENSG00000166326 | TRIM44 | 2.1270268 | 0.00261709 |
| ENSG00000163606 | CD200R1 | 2.12284896 | 0.00152415 |
| ENSG00000112367 | FIG4 | 2.12218668 | 0.00167388 |
| ENSG00000103051 | COG4 | 2.1204803 | 0.00180663 |
| ENSG00000138078 | PREPL | 2.12006781 | 0.01270208 |
| ENSG00000173821 | RNF213 | 2.11768767 | 0.00553015 |
| ENSG00000138246 | DNAJC13 | 2.11570284 | 0.00068553 |
| ENSG00000159131 | GART | 2.11358372 | 0.00326645 |
| ENSG00000081087 | OSTM1 | 2.11289772 | 0.01186229 |
| ENSG00000101336 | HCK | 2.09767392 | 0.00418283 |
| ENSG00000074706 | IPCEF1 | 2.09384519 | 0.02342212 |
| ENSG00000175857 | GAPT | 2.08765079 | 0.00150263 |
| ENSG00000095585 | BLNK | 2.08376627 | 0.00803214 |
| ENSG00000144218 | AFF3 | 2.08136827 | 0.01047324 |
| ENSG00000136279 | DBNL | 2.07401784 | 0.00439901 |
| ENSG00000158517 | NCF1 | 2.06949637 | 0.02847331 |
| ENSG00000166888 | STAT6 | 2.06811372 | 0.00224048 |
| ENSG00000102189 | EEA1 | 2.06093793 | 0.00236146 |
| ENSG00000280153 | AC133065.6 | 2.06005278 | 0.01070935 |
| ENSG00000163563 | MNDA | 2.05970748 | 0.0043897 |
| ENSG00000116824 | CD2 | 2.0576007 | 0.00073864 |
| ENSG00000117054 | ACADM | 2.05707859 | 0.00152894 |
| ENSG00000162736 | NCSTN | 2.05415475 | 0.00194739 |
| ENSG00000150681 | RGS18 | 2.05320806 | 0.00130443 |
| ENSG00000131844 | MCCC2 | 2.0530904 | 0.01886776 |
| ENSG00000107099 | DOCK8 | 2.0516027 | 0.00570959 |
| ENSG00000198814 | GK | 2.04963172 | 0.00350313 |
| ENSG00000148634 | HERC4 | 2.04789706 | 0.00411384 |
| ENSG00000134851 | TMEM165 | 2.04511559 | 0.00084979 |
| ENSG00000136040 | PLXNC1 | 2.04407099 | 0.00885173 |
| ENSG00000139641 | ESYT1 | 2.02820528 | 0.00268199 |
| ENSG00000048052 | HDAC9 | 2.02033464 | 0.01399335 |
| ENSG00000124357 | NAGK | 2.01980368 | 0.0012446 |
| ENSG00000158714 | SLAMF8 | 2.01655113 | 0.02493296 |
| ENSG00000090863 | GLG1 | 2.01630472 | 0.00703198 |
| ENSG00000134321 | RSAD2 | 2.01046868 | 0.00489118 |
| ENSG00000130021 | PUDP | 2.00722498 | 0.00419863 |
| ENSG00000031081 | ARHGAP31 | 2.00094561 | 0.00235967 |
| ENSG00000164062 | APEH | 1.99681356 | 0.00478686 |

| | | | |
|---|---|---|---|
| ENSG00000158467 | AHCYL2 | 1.98490934 | 0.01332182 |
| ENSG00000184979 | USP18 | 1.98276515 | 0.00785233 |
| ENSG00000116990 | MYCL | 1.98235502 | 0.00516637 |
| ENSG00000180353 | HCLS1 | 1.98216761 | 0.00090802 |
| ENSG00000121210 | TMEM131L | 1.98071161 | 0.00224427 |
| ENSG00000142089 | IFITM3 | 1.97921372 | 0.01850953 |
| ENSG00000100364 | KIAA0930 | 1.97310857 | 0.00090772 |
| ENSG00000070785 | EIF2B3 | 1.97182446 | 0.00068553 |
| ENSG00000163565 | IFI16 | 1.97135082 | 0.00322266 |
| ENSG00000134955 | SLC37A2 | 1.97090771 | 0.00376598 |
| ENSG00000079950 | STX7 | 1.96308556 | 0.00166984 |
| ENSG00000058668 | ATP2B4 | 1.96027503 | 0.00934972 |
| ENSG00000103544 | C16orf62 | 1.96011824 | 0.00248227 |
| ENSG00000141510 | TP53 | 1.9574839 | 0.00284279 |
| ENSG00000132182 | NUP210 | 1.94812233 | 0.01100861 |
| ENSG00000188419 | CHM | 1.94649213 | 0.00235025 |
| ENSG00000134256 | CD101 | 1.94646613 | 0.0034249 |
| ENSG00000138646 | HERC5 | 1.94343046 | 0.00898743 |
| ENSG00000169220 | RGS14 | 1.94080353 | 0.00276203 |
| ENSG00000145348 | TBCK | 1.93932334 | 0.00398077 |
| ENSG00000155957 | TMBIM4 | 1.93849172 | 0.00057232 |
| ENSG00000134255 | CEPT1 | 1.93725411 | 0.00656895 |
| ENSG00000112118 | MCM3 | 1.93115618 | 0.00553015 |
| ENSG00000134910 | STT3A | 1.92914174 | 0.01415762 |
| ENSG00000157637 | SLC38A10 | 1.92596603 | 0.00200912 |
| ENSG00000100030 | MAPK1 | 1.92079694 | 0.00089939 |
| ENSG00000137752 | CASP1 | 1.91625457 | 0.00255612 |
| ENSG00000156110 | ADK | 1.91186848 | 0.03702661 |
| ENSG00000010671 | BTK | 1.9062616 | 0.00350274 |
| ENSG00000134318 | ROCK2 | 1.90399304 | 0.00430676 |
| ENSG00000160310 | PRMT2 | 1.9005587 | 0.00351578 |
| ENSG00000130309 | COLGALT1 | 1.90004736 | 0.00460345 |
| ENSG00000165071 | TMEM71 | 1.89529701 | 0.00356837 |
| ENSG00000243943 | ZNF512 | 1.88990246 | 0.00303209 |
| ENSG00000132256 | TRIM5 | 1.88565672 | 0.00229084 |
| ENSG00000273749 | CYFIP1 | 1.88474249 | 0.0066992 |
| ENSG00000133313 | CNDP2 | 1.88141659 | 0.0032195 |
| ENSG00000178537 | SLC25A20 | 1.88126405 | 0.00747632 |
| ENSG00000138642 | HERC6 | 1.88069728 | 0.00981248 |
| ENSG00000104998 | IL27RA | 1.88021435 | 0.00089939 |
| ENSG00000184178 | SCFD2 | 1.87811894 | 0.00194739 |
| ENSG00000150867 | PIP4K2A | 1.87751101 | 0.00205261 |

| | | | |
|---|---|---|---|
| ENSG00000135899 | SP110 | 1.87509223 | 0.00286812 |
| ENSG00000170854 | RIOX2 | 1.87144549 | 0.00472286 |
| ENSG00000214114 | MYCBP | 1.87095474 | 0.00322031 |
| ENSG00000156587 | UBE2L6 | 1.86989923 | 0.00542396 |
| ENSG00000059377 | TBXAS1 | 1.86972621 | 0.00256622 |
| ENSG00000187688 | TRPV2 | 1.8686118 | 0.01599641 |
| ENSG00000111540 | RAB5B | 1.85822044 | 0.00226659 |
| ENSG00000106780 | MEGF9 | 1.85807432 | 0.00309933 |
| ENSG00000110079 | MS4A4A | 1.85718018 | 0.00107684 |
| ENSG00000169403 | PTAFR | 1.85688839 | 0.0091414 |
| ENSG00000104972 | LILRB1 | 1.85246369 | 0.00656277 |
| ENSG00000188404 | SELL | 1.84886004 | 0.02601302 |
| ENSG00000204261 | PSMB8-AS1 | 1.84595288 | 0.00947011 |
| ENSG00000103423 | DNAJA3 | 1.84202531 | 0.00058696 |
| ENSG00000008869 | HEATR5B | 1.8416768 | 0.00089939 |
| ENSG00000198951 | NAGA | 1.84079564 | 0.00525137 |
| ENSG00000144468 | RHBDD1 | 1.83990652 | 0.00652895 |
| ENSG00000185722 | ANKFY1 | 1.83883439 | 0.01278329 |
| ENSG00000136518 | ACTL6A | 1.83498314 | 0.00330233 |
| ENSG00000103479 | RBL2 | 1.83345605 | 0.00107684 |
| ENSG00000163644 | PPM1K | 1.83301931 | 0.00810903 |
| ENSG00000040933 | INPP4A | 1.8290558 | 0.00351201 |
| ENSG00000109911 | ELP4 | 1.82900732 | 0.00431962 |
| ENSG00000143799 | PARP1 | 1.82636685 | 0.00888272 |
| ENSG00000126709 | IFI6 | 1.82601023 | 0.00130443 |
| ENSG00000143252 | SDHC | 1.82148734 | 0.00109198 |
| ENSG00000178175 | ZNF366 | 1.81805716 | 0.00877547 |
| ENSG00000161955 | TNFSF13 | 1.8140582 | 0.00155034 |
| ENSG00000213445 | SIPA1 | 1.81355362 | 0.00155933 |
| ENSG00000187608 | ISG15 | 1.81101135 | 0.00349422 |
| ENSG00000136100 | VPS36 | 1.8056376 | 0.00393222 |
| ENSG00000014216 | CAPN1 | 1.80513921 | 0.00068553 |
| ENSG00000102316 | MAGED2 | 1.80374069 | 0.00599034 |
| ENSG00000182179 | UBA7 | 1.79549594 | 0.00581898 |
| ENSG00000129675 | ARHGEF6 | 1.79496177 | 0.00572497 |
| ENSG00000143110 | C1orf162 | 1.79134014 | 0.00113898 |
| ENSG00000111667 | USP5 | 1.78841999 | 0.00362829 |
| ENSG00000111481 | COPZ1 | 1.78766003 | 0.00155933 |
| ENSG00000014919 | COX15 | 1.78480081 | 0.01278329 |
| ENSG00000196511 | TPK1 | 1.78455658 | 0.00084372 |
| ENSG00000138496 | PARP9 | 1.78403341 | 0.0028424 |
| ENSG00000168538 | TRAPPC11 | 1.78376308 | 0.00095746 |

| | | | |
|---|---|---|---|
| ENSG00000102710 | SUPT20H | 1.78220368 | 0.0016072 |
| ENSG00000102524 | TNFSF13B | 1.78177857 | 0.0015584 |
| ENSG00000136404 | TM6SF1 | 1.7797517 | 0.01106233 |
| ENSG00000143554 | SLC27A3 | 1.77628553 | 0.00383452 |
| ENSG00000165675 | ENOX2 | 1.77518869 | 0.00531959 |
| ENSG00000119471 | HSDL2 | 1.77443536 | 0.01839835 |
| ENSG00000166002 | SMCO4 | 1.771189 | 0.00652389 |
| ENSG00000160712 | IL6R | 1.76704675 | 0.00509474 |
| ENSG00000243749 | TMEM35B | 1.76307995 | 0.002218 |
| ENSG00000213983 | AP1G2 | 1.7561938 | 0.02118998 |
| ENSG00000111731 | C2CD5 | 1.74901382 | 0.00089939 |
| ENSG00000081189 | MEF2C | 1.74589214 | 0.00328847 |
| ENSG00000108798 | ABI3 | 1.74419429 | 0.00606651 |
| ENSG00000140395 | WDR61 | 1.74340544 | 0.00568 |
| ENSG00000162704 | ARPC5 | 1.74224952 | 0.00236146 |
| ENSG00000104133 | SPG11 | 1.7409701 | 0.02029751 |
| ENSG00000088682 | COQ9 | 1.74064775 | 0.00089939 |
| ENSG00000135317 | SNX14 | 1.74050946 | 0.00068553 |
| ENSG00000149311 | ATM | 1.73933874 | 0.00246913 |
| ENSG00000125124 | BBS2 | 1.73821256 | 0.00386312 |
| ENSG00000197798 | FAM118B | 1.73605372 | 0.01373726 |
| ENSG00000105639 | JAK3 | 1.73481175 | 0.03835979 |
| ENSG00000105726 | ATP13A1 | 1.73232 | 0.00215982 |
| ENSG00000250138 | AC139495.3 | 1.73142998 | 0.01204627 |
| ENSG00000104365 | IKBKB | 1.73039626 | 0.00170624 |
| ENSG00000165476 | REEP3 | 1.72913973 | 0.00190829 |
| ENSG00000140575 | IQGAP1 | 1.72869261 | 0.00068553 |
| ENSG00000189339 | SLC35E2B | 1.72516176 | 0.0086378 |
| ENSG00000112079 | STK38 | 1.72055039 | 0.00243303 |
| ENSG00000149196 | HIKESHI | 1.71739712 | 0.00080111 |
| ENSG00000160883 | HK3 | 1.71050513 | 0.00336571 |
| ENSG00000197157 | SND1 | 1.71045368 | 0.00235967 |
| ENSG00000134452 | FBXO18 | 1.70474196 | 0.00390878 |
| ENSG00000133103 | COG6 | 1.70350221 | 0.03295002 |
| ENSG00000172269 | DPAGT1 | 1.7034649 | 0.00923715 |
| ENSG00000071967 | CYBRD1 | 1.70229963 | 0.01899945 |
| ENSG00000169116 | PARM1 | 1.70212386 | 0.00581841 |
| ENSG00000132383 | RPA1 | 1.70063799 | 0.00278533 |
| ENSG00000121067 | SPOP | 1.69949896 | 0.00398133 |
| ENSG00000187554 | TLR5 | 1.6992318 | 0.02127473 |
| ENSG00000163154 | TNFAIP8L2 | 1.69414654 | 0.00601809 |
| ENSG00000154822 | PLCL2 | 1.69378163 | 0.00852904 |

| | | | |
|---|---|---|---|
| ENSG00000219545 | UMAD1 | 1.69103006 | 0.00229084 |
| ENSG00000166797 | FAM96A | 1.690023 | 0.01107601 |
| ENSG00000026751 | SLAMF7 | 1.6850367 | 0.00419863 |
| ENSG00000155926 | SLA | 1.68407908 | 0.00419863 |
| ENSG00000030066 | NUP160 | 1.68341977 | 0.0034249 |
| ENSG00000174944 | P2RY14 | 1.68228319 | 0.00359283 |
| ENSG00000164308 | ERAP2 | 1.68187261 | 0.00204546 |
| ENSG00000198876 | DCAF12 | 1.6788135 | 0.00418283 |
| ENSG00000099810 | MTAP | 1.67769763 | 0.00688879 |
| ENSG00000157483 | MYO1E | 1.67761878 | 0.00615876 |
| ENSG00000104518 | GSDMD | 1.67618977 | 0.00221039 |
| ENSG00000119922 | IFIT2 | 1.675019 | 0.00402472 |
| ENSG00000033050 | ABCF2 | 1.67497452 | 0.00210312 |
| ENSG00000163840 | DTX3L | 1.67472708 | 0.00338864 |
| ENSG00000170006 | TMEM154 | 1.67392517 | 0.0078963 |
| ENSG00000116005 | PCYOX1 | 1.669621 | 0.00077204 |
| ENSG00000121691 | CAT | 1.66938691 | 0.00264994 |
| ENSG00000152683 | SLC30A6 | 1.66696901 | 0.02358751 |
| ENSG00000185973 | TMLHE | 1.66541486 | 0.00719883 |
| ENSG00000138768 | USO1 | 1.66532486 | 0.00160581 |
| ENSG00000167797 | CDK2AP2 | 1.66381151 | 0.00464174 |
| ENSG00000117228 | GBP1 | 1.66192443 | 0.03832356 |
| ENSG00000125863 | MKKS | 1.661671 | 0.00340571 |
| ENSG00000143493 | INTS7 | 1.66145404 | 0.00358122 |
| ENSG00000256043 | CTSO | 1.65905713 | 0.00201407 |
| ENSG00000115234 | SNX17 | 1.65477931 | 0.00155933 |
| ENSG00000139687 | RB1 | 1.65432967 | 0.00599034 |
| ENSG00000134996 | OSTF1 | 1.65369005 | 0.00815818 |
| ENSG00000124942 | AHNAK | 1.65160242 | 0.01310182 |
| ENSG00000132646 | PCNA | 1.65025522 | 0.00075186 |
| ENSG00000119686 | FLVCR2 | 1.6496537 | 0.00409363 |
| ENSG00000139278 | GLIPR1 | 1.64663546 | 0.00668554 |
| ENSG00000143390 | RFX5 | 1.644153 | 0.0028804 |
| ENSG00000176783 | RUFY1 | 1.64278813 | 0.00019288 |
| ENSG00000110934 | BIN2 | 1.64169805 | 0.0076335 |
| ENSG00000162946 | DISC1 | 1.63975313 | 0.02702117 |
| ENSG00000118855 | MFSD1 | 1.63886239 | 0.00340003 |
| ENSG00000115204 | MPV17 | 1.63630852 | 0.00589844 |
| ENSG00000185885 | IFITM1 | 1.636154 | 0.0100445 |
| ENSG00000130150 | MOSPD2 | 1.63384537 | 0.02943076 |
| ENSG00000103047 | TANGO6 | 1.63139604 | 0.01973655 |
| ENSG00000116701 | NCF2 | 1.62979246 | 0.02288169 |

| | | | |
|---|---|---|---|
| ENSG00000100714 | MTHFD1 | 1.62878459 | 0.00744894 |
| ENSG00000110665 | C11orf21 | 1.62866074 | 0.00233277 |
| ENSG00000006756 | ARSD | 1.62737992 | 0.00261478 |
| ENSG00000167851 | CD300A | 1.62451488 | 0.00424199 |
| ENSG00000186687 | LYRM7 | 1.62448007 | 0.01398358 |
| ENSG00000196510 | ANAPC7 | 1.62292839 | 0.01199328 |
| ENSG00000122025 | FLT3 | 1.62276623 | 0.00503046 |
| ENSG00000170876 | TMEM43 | 1.62113951 | 0.00351394 |
| ENSG00000104325 | DECR1 | 1.61561589 | 0.00648504 |
| ENSG00000183978 | COA3 | 1.61537192 | 0.00546142 |
| ENSG00000127946 | HIP1 | 1.61235796 | 0.01311257 |
| ENSG00000131828 | PDHA1 | 1.61121525 | 0.00224048 |
| ENSG00000138375 | SMARCAL1 | 1.60521453 | 0.01682839 |
| ENSG00000064115 | TM7SF3 | 1.60501436 | 0.02874989 |
| ENSG00000120860 | WASHC3 | 1.59995858 | 0.00951642 |
| ENSG00000181192 | DHTKD1 | 1.59870003 | 0.00337093 |
| ENSG00000101337 | TM9SF4 | 1.59776353 | 0.00419863 |
| ENSG00000124508 | BTN2A2 | 1.59672095 | 0.00803266 |
| ENSG00000163932 | PRKCD | 1.59396075 | 0.00295864 |
| ENSG00000197548 | ATG7 | 1.59160887 | 0.00155933 |
| ENSG00000092531 | SNAP23 | 1.59108194 | 0.00330233 |
| ENSG00000128915 | ICE2 | 1.59062394 | 0.00580111 |
| ENSG00000136874 | STX17 | 1.59054099 | 0.00272216 |
| ENSG00000163389 | POGLUT1 | 1.58939529 | 0.00563712 |
| ENSG00000120451 | SNX19 | 1.58640112 | 0.00126831 |
| ENSG00000155659 | VSIG4 | 1.58489352 | 0.0408237 |
| ENSG00000137478 | FCHSD2 | 1.58247814 | 0.00660914 |
| ENSG00000160551 | TAOK1 | 1.58239354 | 0.0150404 |
| ENSG00000129691 | ASH2L | 1.58217542 | 0.00478686 |
| ENSG00000155660 | PDIA4 | 1.57763287 | 0.00398077 |
| ENSG00000100201 | DDX17 | 1.57737112 | 0.03353705 |
| ENSG00000113522 | RAD50 | 1.57335076 | 0.00398077 |
| ENSG00000091106 | NLRC4 | 1.57239752 | 0.01243906 |
| ENSG00000124356 | STAMBP | 1.57147346 | 0.02912533 |
| ENSG00000145246 | ATP10D | 1.57110413 | 0.01169583 |
| ENSG00000134014 | ELP3 | 1.57089038 | 0.01831573 |
| ENSG00000142687 | KIAA0319L | 1.57079909 | 0.00272842 |
| ENSG00000250687 | AC146944.2 | 1.57069338 | 0.01618458 |
| ENSG00000143106 | PSMA5 | 1.57054856 | 0.00303209 |
| ENSG00000106049 | HIBADH | 1.56946545 | 0.00155933 |
| ENSG00000008130 | NADK | 1.56913963 | 0.00155933 |
| ENSG00000173193 | PARP14 | 1.56722447 | 0.04430578 |

| | | | |
|---|---|---|---|
| ENSG00000166145 | SPINT1 | 1.56706073 | 0.00261709 |
| ENSG00000168010 | ATG16L2 | 1.56683923 | 0.01188302 |
| ENSG00000163946 | FAM208A | 1.56522132 | 0.00837266 |
| ENSG00000139163 | ETNK1 | 1.56452612 | 0.00391676 |
| ENSG00000123106 | CCDC91 | 1.56162639 | 0.00097295 |
| ENSG00000135842 | FAM129A | 1.56093832 | 0.00961836 |
| ENSG00000184992 | BRI3BP | 1.56076587 | 0.00653628 |
| ENSG00000130985 | UBA1 | 1.55856912 | 0.01170173 |
| ENSG00000007923 | DNAJC11 | 1.55848947 | 0.00232308 |
| ENSG00000229754 | CXCR2P1 | 1.55812835 | 0.0028754 |
| ENSG00000132274 | TRIM22 | 1.55293658 | 0.00530367 |
| ENSG00000148180 | GSN | 1.55125534 | 0.03250746 |
| ENSG00000124532 | MRS2 | 1.54854827 | 0.00267271 |
| ENSG00000119906 | SLF2 | 1.54812256 | 0.00192901 |
| ENSG00000138594 | TMOD3 | 1.5475971 | 0.00349422 |
| ENSG00000254505 | CHMP4A | 1.54643452 | 0.00089939 |
| ENSG00000147251 | DOCK11 | 1.54569043 | 0.00489336 |
| ENSG00000197943 | PLCG2 | 1.54491224 | 0.01603467 |
| ENSG00000155229 | MMS19 | 1.54354691 | 0.00355307 |
| ENSG00000108946 | PRKAR1A | 1.54287173 | 0.00987036 |
| ENSG00000136628 | EPRS | 1.54248327 | 0.00578374 |
| ENSG00000101096 | NFATC2 | 1.5422227 | 0.00697219 |
| ENSG00000100889 | PCK2 | 1.53930727 | 0.0174206 |
| ENSG00000158481 | CD1C | 1.53806136 | 0.02980208 |
| ENSG00000018699 | TTC27 | 1.53740158 | 0.00401831 |
| ENSG00000171307 | ZDHHC16 | 1.53552972 | 0.00214395 |
| ENSG00000185324 | CDK10 | 1.53547738 | 0.00343119 |
| ENSG00000149269 | PAK1 | 1.53498981 | 0.04782639 |
| ENSG00000151116 | UEVLD | 1.53468566 | 0.00115627 |
| ENSG00000138363 | ATIC | 1.53434242 | 0.00548977 |
| ENSG00000108679 | LGALS3BP | 1.5335672 | 0.00709936 |
| ENSG00000137845 | ADAM10 | 1.53333701 | 0.00648504 |
| ENSG00000104812 | GYS1 | 1.53254233 | 0.00647532 |
| ENSG00000167272 | POP5 | 1.53225337 | 0.00476653 |
| ENSG00000109854 | HTATIP2 | 1.53116263 | 0.01097056 |
| ENSG00000113552 | GNPDA1 | 1.52780061 | 0.00116354 |
| ENSG00000189091 | SF3B3 | 1.52570221 | 0.0027654 |
| ENSG00000123213 | NLN | 1.52198051 | 0.00810903 |
| ENSG00000138660 | AP1AR | 1.5204922 | 0.01532212 |
| ENSG00000088888 | MAVS | 1.52026475 | 0.02330006 |
| ENSG00000160326 | SLC2A6 | 1.51991534 | 0.01121565 |
| ENSG00000137509 | PRCP | 1.51757796 | 0.02010187 |

| | | | |
|---|---|---|---|
| ENSG00000168710 | AHCYL1 | 1.51740183 | 0.00205904 |
| ENSG00000166986 | MARS | 1.51668507 | 0.00786406 |
| ENSG00000156017 | CARNMT1 | 1.51223681 | 0.00605285 |
| ENSG00000134369 | NAV1 | 1.51142591 | 0.01566357 |
| ENSG00000204713 | TRIM27 | 1.51001976 | 0.0028334 |
| ENSG00000151414 | NEK7 | 1.50940757 | 0.00142767 |
| ENSG00000188352 | FOCAD | 1.50803856 | 0.0140459 |
| ENSG00000085224 | ATRX | 1.50709992 | 0.00481717 |
| ENSG00000100442 | FKBP3 | 1.50707858 | 0.00393222 |
| ENSG00000010292 | NCAPD2 | 1.50618187 | 0.01643738 |
| ENSG00000118961 | LDAH | 1.50329037 | 0.00088661 |
| ENSG00000092929 | UNC13D | 1.50312908 | 0.00362829 |
| ENSG00000108439 | PNPO | 1.50146753 | 0.00837266 |
| ENSG00000009790 | TRAF3IP3 | 1.50121255 | 0.00174835 |
| ENSG00000136169 | SETDB2 | 1.50094086 | 0.00497876 |
| ENSG00000110931 | CAMKK2 | 1.50039673 | 0.0048661 |
| ENSG00000123689 | G0S2 | -7.6084168 | 0.02613793 |
| ENSG00000161921 | CXCL16 | -4.5620958 | 0.01042913 |
| ENSG00000112149 | CD83 | -4.5009312 | 0.03606944 |
| ENSG00000095794 | CREM | -4.4779198 | 0.01900203 |
| ENSG00000166920 | C15orf48 | -4.0542904 | 0.01764021 |
| ENSG00000113448 | PDE4D | -3.9972163 | 0.02161677 |
| ENSG00000118515 | SGK1 | -3.9960238 | 0.00395078 |
| ENSG00000088826 | SMOX | -3.9659598 | 0.03956602 |
| ENSG00000120063 | GNA13 | -3.8727133 | 0.02583179 |
| ENSG00000162496 | DHRS3 | -3.8716416 | 0.01786884 |
| ENSG00000090104 | RGS1 | -3.8422815 | 0.02570437 |
| ENSG00000163376 | KBTBD8 | -3.8282484 | 0.00666811 |
| ENSG00000104312 | RIPK2 | -3.80965 | 0.00516045 |
| ENSG00000186594 | MIR22HG | -3.717995 | 0.00597583 |
| ENSG00000100644 | HIF1A | -3.5700622 | 0.00489336 |
| ENSG00000170525 | PFKFB3 | -3.4828636 | 0.0356038 |
| ENSG00000140379 | BCL2A1 | -3.4594922 | 0.01833318 |
| ENSG00000122644 | ARL4A | -3.4288512 | 0.00758077 |
| ENSG00000130340 | SNX9 | -3.4221805 | 0.01121546 |
| ENSG00000008083 | JARID2 | -3.3861327 | 0.01931415 |
| ENSG00000143507 | DUSP10 | -3.351403 | 0.00758077 |
| ENSG00000217801 | AL390719.1 | -3.3393149 | 0.04447417 |
| ENSG00000145860 | RNF145 | -3.3305447 | 0.00043805 |
| ENSG00000169155 | ZBTB43 | -3.3242159 | 0.00073864 |
| ENSG00000184205 | TSPYL2 | -3.3190953 | 0.01986667 |
| ENSG00000131669 | NINJ1 | -3.3057839 | 0.00354594 |

| | | | |
|---|---|---|---|
| ENSG00000160223 | ICOSLG | -3.244978 | 0.00747632 |
| ENSG00000231721 | LINC-PINT | -3.237142 | 0.02215288 |
| ENSG00000173166 | RAPH1 | -3.2366725 | 0.02706525 |
| ENSG00000255073 | ZFP91-CNTF | -3.2069268 | 0.00316541 |
| ENSG00000155307 | SAMSN1 | -3.1958956 | 0.01212833 |
| ENSG00000169508 | GPR183 | -3.1649492 | 0.00084979 |
| ENSG00000167996 | FTH1 | -3.1220734 | 0.00418283 |
| ENSG00000153234 | NR4A2 | -3.1215849 | 0.03084488 |
| ENSG00000113070 | HBEGF | -3.1121736 | 0.01863785 |
| ENSG00000173334 | TRIB1 | -3.1114145 | 0.0220503 |
| ENSG00000154640 | BTG3 | -3.1046384 | 0.00133507 |
| ENSG00000166128 | RAB8B | -3.1014849 | 0.03010848 |
| ENSG00000132906 | CASP9 | -3.0953133 | 0.03939224 |
| ENSG00000180530 | NRIP1 | -3.0636904 | 0.00339177 |
| ENSG00000125538 | IL1B | -3.0506771 | 0.00688974 |
| ENSG00000135604 | STX11 | -3.039988 | 0.0030748 |
| ENSG00000120705 | ETF1 | -3.036428 | 0.00817205 |
| ENSG00000011422 | PLAUR | -3.0294006 | 0.00817205 |
| ENSG00000272888 | LINC01578 | -3.0146861 | 0.01208127 |
| ENSG00000165030 | NFIL3 | -3.0104971 | 0.00601809 |
| ENSG00000168036 | CTNNB1 | -3.0086254 | 0.01162234 |
| ENSG00000146457 | WTAP | -2.9875884 | 0.00311935 |
| ENSG00000277117 | FP565260.3 | -2.9712348 | 0.0112941 |
| ENSG00000143622 | RIT1 | -2.9564428 | 0.02044292 |
| ENSG00000102760 | RGCC | -2.9555816 | 0.00503046 |
| ENSG00000175130 | MARCKSL1 | -2.9550318 | 0.02165853 |
| ENSG00000171488 | LRRC8C | -2.9124449 | 0.00987036 |
| ENSG00000140564 | FURIN | -2.9117479 | 0.02813783 |
| ENSG00000132952 | USPL1 | -2.8997561 | 0.0030748 |
| ENSG00000169895 | SYAP1 | -2.8975029 | 0.03098433 |
| ENSG00000132819 | RBM38 | -2.8929631 | 0.02885702 |
| ENSG00000057657 | PRDM1 | -2.8814963 | 0.02297287 |
| ENSG00000106004 | HOXA5 | -2.8736118 | 0.01402569 |
| ENSG00000137936 | BCAR3 | -2.8427275 | 0.04935297 |
| ENSG00000117036 | ETV3 | -2.8424885 | 0.04129631 |
| ENSG00000140743 | CDR2 | -2.8211935 | 0.01278329 |
| ENSG00000175040 | CHST2 | -2.8202678 | 0.04101553 |
| ENSG00000197170 | PSMD12 | -2.8030745 | 0.00068553 |
| ENSG00000211445 | GPX3 | -2.7908858 | 0.0279606 |
| ENSG00000075426 | FOSL2 | -2.784013 | 0.02667059 |
| ENSG00000138166 | DUSP5 | -2.7541356 | 0.02403311 |
| ENSG00000162772 | ATF3 | -2.7312135 | 0.00375073 |

| | | | |
|---|---|---|---|
| ENSG00000185022 | MAFF | -2.7224192 | 0.01710678 |
| ENSG00000114784 | EIF1B | -2.7198084 | 0.00156522 |
| ENSG00000186660 | ZFP91 | -2.7136779 | 0.00360138 |
| ENSG00000110852 | CLEC2B | -2.7117292 | 0.03716562 |
| ENSG00000125657 | TNFSF9 | -2.6940337 | 0.00170624 |
| ENSG00000162711 | NLRP3 | -2.6934593 | 0.02305255 |
| ENSG00000271614 | ATP2B1-AS1 | -2.6879716 | 0.00430935 |
| ENSG00000011566 | MAP4K3 | -2.6859818 | 0.01043629 |
| ENSG00000277632 | CCL3 | -2.6812142 | 0.00193765 |
| ENSG00000141682 | PMAIP1 | -2.6736041 | 0.00340571 |
| ENSG00000025156 | HSF2 | -2.6655426 | 0.00107181 |
| ENSG00000015475 | BID | -2.632563 | 0.03295002 |
| ENSG00000261740 | BOLA2-SMG1P6 | -2.629706 | 0.00122855 |
| ENSG00000136738 | STAM | -2.6137392 | 0.01121546 |
| ENSG00000239305 | RNF103 | -2.6050891 | 0.00152893 |
| ENSG00000117000 | RLF | -2.604355 | 0.00200912 |
| ENSG00000102908 | NFAT5 | -2.6031591 | 0.0113783 |
| ENSG00000100600 | LGMN | -2.5968497 | 0.04577179 |
| ENSG00000173812 | EIF1 | -2.5962337 | 0.00204067 |
| ENSG00000110721 | CHKA | -2.581298 | 0.01251936 |
| ENSG00000180628 | PCGF5 | -2.5780265 | 0.00326645 |
| ENSG00000105835 | NAMPT | -2.5580719 | 0.0357451 |
| ENSG00000131408 | NR1H2 | -2.5564414 | 0.0084248 |
| ENSG00000228830 | AL160408.2 | -2.5536041 | 0.02501364 |
| ENSG00000122068 | FYTTD1 | -2.5491067 | 0.00096838 |
| ENSG00000123091 | RNF11 | -2.5393281 | 0.00356837 |
| ENSG00000147119 | CHST7 | -2.5367945 | 0.00858563 |
| ENSG00000221869 | CEBPD | -2.533836 | 0.00120407 |
| ENSG00000141506 | PIK3R5 | -2.5294937 | 0.04023237 |
| ENSG00000139832 | RAB20 | -2.5208739 | 0.01470183 |
| ENSG00000020633 | RUNX3 | -2.5173964 | 0.00609972 |
| ENSG00000120616 | EPC1 | -2.511977 | 0.00599216 |
| ENSG00000156671 | SAMD8 | -2.5026093 | 0.00182038 |
| ENSG00000118689 | FOXO3 | -2.4834505 | 0.00654052 |
| ENSG00000112511 | PHF1 | -2.4821148 | 0.03838578 |
| ENSG00000168264 | IRF2BP2 | -2.4705596 | 0.0088892 |
| ENSG00000136950 | ARPC5L | -2.4661936 | 0.00546681 |
| ENSG00000123358 | NR4A1 | -2.4608666 | 0.00169166 |
| ENSG00000144802 | NFKBIZ | -2.460033 | 0.01089345 |
| ENSG00000155252 | PI4K2A | -2.4575166 | 0.00425322 |
| ENSG00000126524 | SBDS | -2.4403481 | 0.00186519 |

| | | | |
|---|---|---|---|
| ENSG00000138069 | RAB1A | -2.4313541 | 0.00759346 |
| ENSG00000196843 | ARID5A | -2.4206069 | 0.00418283 |
| ENSG00000086062 | B4GALT1 | -2.4193901 | 0.01382481 |
| ENSG00000175606 | TMEM70 | -2.4157729 | 0.00451031 |
| ENSG00000105856 | HBP1 | -2.4083623 | 0.00068687 |
| ENSG00000055483 | USP36 | -2.4070147 | 0.02957349 |
| ENSG00000113369 | ARRDC3 | -2.4042168 | 0.02261771 |
| ENSG00000100614 | PPM1A | -2.3961858 | 0.00130443 |
| ENSG00000107937 | GTPBP4 | -2.3954039 | 0.01170173 |
| ENSG00000145675 | PIK3R1 | -2.3823332 | 0.02075998 |
| ENSG00000171867 | PRNP | -2.3817915 | 0.00187431 |
| ENSG00000155304 | HSPA13 | -2.3758936 | 0.00068553 |
| ENSG00000013441 | CLK1 | -2.3717733 | 0.00509881 |
| ENSG00000123908 | AGO2 | -2.3688319 | 0.04186509 |
| ENSG00000059804 | SLC2A3 | -2.3674552 | 0.03311309 |
| ENSG00000177426 | TGIF1 | -2.3650867 | 0.00541373 |
| ENSG00000196428 | TSC22D2 | -2.3621518 | 0.00084979 |
| ENSG00000152484 | USP12 | -2.3562138 | 0.00628144 |
| ENSG00000128272 | ATF4 | -2.3559686 | 0.00089939 |
| ENSG00000087074 | PPP1R15A | -2.3477231 | 0.01230525 |
| ENSG00000213923 | CSNK1E | -2.3443671 | 0.00599034 |
| ENSG00000166165 | CKB | -2.3365594 | 0.00191753 |
| ENSG00000122862 | SRGN | -2.3273528 | 0.00167388 |
| ENSG00000124762 | CDKN1A | -2.3268303 | 0.00479775 |
| ENSG00000137409 | MTCH1 | -2.3223044 | 0.01442877 |
| ENSG00000101558 | VAPA | -2.3187794 | 0.00701573 |
| ENSG00000171522 | PTGER4 | -2.3166235 | 0.00843969 |
| ENSG00000025772 | TOMM34 | -2.315441 | 0.00854947 |
| ENSG00000120690 | ELF1 | -2.3099373 | 0.00131723 |
| ENSG00000186162 | CIDECP | -2.3095705 | 0.00460845 |
| ENSG00000115165 | CYTIP | -2.3070301 | 0.0084484 |
| ENSG00000173575 | CHD2 | -2.300325 | 0.00152893 |
| ENSG00000151553 | FAM160B1 | -2.2992749 | 0.00272842 |
| ENSG00000259884 | AC025259.3 | -2.2984853 | 0.00153329 |
| ENSG00000176946 | THAP4 | -2.2948812 | 0.02164364 |
| ENSG00000100906 | NFKBIA | -2.2869631 | 0.00120407 |
| ENSG00000132326 | PER2 | -2.284667 | 0.00190217 |
| ENSG00000083799 | CYLD | -2.2825129 | 0.00342468 |
| ENSG00000105968 | H2AFV | -2.2815598 | 0.00837434 |
| ENSG00000027697 | IFNGR1 | -2.2789363 | 0.00068553 |
| ENSG00000086666 | ZFAND6 | -2.2765428 | 0.0028622 |
| ENSG00000108179 | PPIF | -2.2745121 | 0.01004343 |

| | | | |
|---|---|---|---|
| ENSG00000143153 | ATP1B1 | -2.2691712 | 0.00478686 |
| ENSG00000114098 | ARMC8 | -2.2609608 | 0.00199149 |
| ENSG00000141580 | WDR45B | -2.2609042 | 0.00977231 |
| ENSG00000166669 | ATF7IP2 | -2.2602164 | 0.02069455 |
| ENSG00000082153 | BZW1 | -2.2562513 | 0.01364735 |
| ENSG00000256235 | SMIM3 | -2.2558118 | 0.00772947 |
| ENSG00000116044 | NFE2L2 | -2.255481 | 0.00359283 |
| ENSG00000134107 | BHLHE40 | -2.2548603 | 0.00929148 |
| ENSG00000154710 | RABGEF1 | -2.2496546 | 0.03965112 |
| ENSG00000030110 | BAK1 | -2.2473536 | 0.04265766 |
| ENSG00000185728 | YTHDF3 | -2.2411364 | 0.00209529 |
| ENSG00000069956 | MAPK6 | -2.2409442 | 0.00130054 |
| ENSG00000137331 | IER3 | -2.2285083 | 0.01475667 |
| ENSG00000270681 | AC095055.1 | -2.2274383 | 0.00337861 |
| ENSG00000121966 | CXCR4 | -2.224568 | 0.00709936 |
| ENSG00000165233 | CARD19 | -2.2243595 | 0.02956604 |
| ENSG00000176407 | KCMF1 | -2.220784 | 0.0128914 |
| ENSG00000023734 | STRAP | -2.2162293 | 0.01354867 |
| ENSG00000239857 | GET4 | -2.2154294 | 0.00077204 |
| ENSG00000176845 | METRNL | -2.2127164 | 0.0012446 |
| ENSG00000272886 | DCP1A | -2.2052986 | 6.3344E-05 |
| ENSG00000136603 | SKIL | -2.203748 | 0.03903684 |
| ENSG00000112245 | PTP4A1 | -2.2015299 | 0.0214355 |
| ENSG00000102265 | TIMP1 | -2.1948074 | 0.0034608 |
| ENSG00000079332 | SAR1A | -2.190543 | 0.03812963 |
| ENSG00000005483 | KMT2E | -2.1903478 | 0.02518335 |
| ENSG00000091317 | CMTM6 | -2.1902788 | 0.00520463 |
| ENSG00000101544 | ADNP2 | -2.1847581 | 0.01951335 |
| ENSG00000006451 | RALA | -2.1830848 | 0.01575296 |
| ENSG00000159128 | IFNGR2 | -2.1825726 | 0.00632567 |
| ENSG00000153066 | TXNDC11 | -2.1784838 | 0.00582255 |
| ENSG00000182831 | C16orf72 | -2.1781934 | 0.00940502 |
| ENSG00000005379 | TSPOAP1 | -2.1779897 | 0.04328864 |
| ENSG00000068697 | LAPTM4A | -2.1749313 | 0.01217754 |
| ENSG00000171604 | CXXC5 | -2.1725344 | 0.00509474 |
| ENSG00000119801 | YPEL5 | -2.1686411 | 0.01396869 |
| ENSG00000140941 | MAP1LC3B | -2.1642957 | 0.00272842 |
| ENSG00000117569 | PTBP2 | -2.1642283 | 0.01869289 |
| ENSG00000135018 | UBQLN1 | -2.1582418 | 0.00655063 |
| ENSG00000145241 | CENPC | -2.1575822 | 0.02464372 |
| ENSG00000196396 | PTPN1 | -2.1570102 | 0.03002164 |
| ENSG00000162511 | LAPTM5 | -2.1549726 | 0.03818104 |

| | | | |
|---|---|---|---|
| ENSG00000078269 | SYNJ2 | -2.1548806 | 0.04403393 |
| ENSG00000163961 | RNF168 | -2.1499857 | 0.00578065 |
| ENSG00000134480 | CCNH | -2.1479935 | 0.02504962 |
| ENSG00000092820 | EZR | -2.1437161 | 0.02593285 |
| ENSG00000106610 | STAG3L4 | -2.143527 | 0.00599034 |
| ENSG00000065809 | FAM107B | -2.1424545 | 0.00970995 |
| ENSG00000151247 | EIF4E | -2.141887 | 0.00276203 |
| ENSG00000260708 | AL118516.1 | -2.1401102 | 0.00272216 |
| ENSG00000169251 | NMD3 | -2.1376019 | 0.00360081 |
| ENSG00000005812 | FBXL3 | -2.1372538 | 0.00669784 |
| ENSG00000183696 | UPP1 | -2.1346803 | 0.0030247 |
| ENSG00000101421 | CHMP4B | -2.13348 | 0.0084259 |
| ENSG00000116741 | RGS2 | -2.1331362 | 0.00388647 |
| ENSG00000163877 | SNIP1 | -2.1286004 | 0.00340571 |
| ENSG00000131263 | RLIM | -2.1271499 | 0.00885186 |
| ENSG00000182149 | IST1 | -2.1220026 | 0.00068553 |
| ENSG00000100401 | RANGAP1 | -2.117229 | 0.01141344 |
| ENSG00000115548 | KDM3A | -2.1165221 | 0.00728599 |
| ENSG00000101782 | RIOK3 | -2.1149049 | 0.00464235 |
| ENSG00000185477 | GPRIN3 | -2.113831 | 0.00084979 |
| ENSG00000107263 | RAPGEF1 | -2.1097344 | 0.02971227 |
| ENSG00000184007 | PTP4A2 | -2.1085876 | 0.00362616 |
| ENSG00000019995 | ZRANB1 | -2.1078433 | 0.00541373 |
| ENSG00000224531 | SMIM13 | -2.1078034 | 0.02714729 |
| ENSG00000205423 | CNEP1R1 | -2.1062582 | 0.02888384 |
| ENSG00000130522 | JUND | -2.1045991 | 0.00709518 |
| ENSG00000171988 | JMJD1C | -2.1019 | 0.00283377 |
| ENSG00000121274 | PAPD5 | -2.0987985 | 0.03894852 |
| ENSG00000106635 | BCL7B | -2.092901 | 0.00023522 |
| ENSG00000138081 | FBXO11 | -2.082102 | 0.00084979 |
| ENSG00000184182 | UBE2F | -2.0783818 | 0.00465281 |
| ENSG00000140332 | TLE3 | -2.0771629 | 0.02785036 |
| ENSG00000121671 | CRY2 | -2.0757999 | 0.015409 |
| ENSG00000197061 | HIST1H4C | -2.075254 | 0.02335005 |
| ENSG00000138750 | NUP54 | -2.0752146 | 0.00345464 |
| ENSG00000126775 | ATG14 | -2.0750812 | 0.00702311 |
| ENSG00000078804 | TP53INP2 | -2.0749929 | 0.00215171 |
| ENSG00000160570 | DEDD2 | -2.0734776 | 0.00300687 |
| ENSG00000070495 | JMJD6 | -2.0730925 | 0.00020762 |
| ENSG00000135404 | CD63 | -2.0723006 | 0.02502702 |
| ENSG00000137947 | GTF2B | -2.0715962 | 0.00166984 |
| ENSG00000136732 | GYPC | -2.0682301 | 0.00573304 |

| | | | |
|---|---|---|---|
| ENSG00000165195 | PIGA | -2.0645569 | 0.00842185 |
| ENSG00000132475 | H3F3B | -2.0602954 | 0.00669784 |
| ENSG00000111832 | RWDD1 | -2.0599284 | 0.01295877 |
| ENSG00000174574 | AKIRIN1 | -2.0497049 | 0.00218967 |
| ENSG00000164823 | OSGIN2 | -2.0493887 | 0.00240547 |
| ENSG00000117318 | ID3 | -2.0457496 | 0.03864295 |
| ENSG00000085433 | WDR47 | -2.0428813 | 0.01756356 |
| ENSG00000102034 | ELF4 | -2.0396186 | 0.02228775 |
| ENSG00000101216 | GMEB2 | -2.039088 | 0.00363002 |
| ENSG00000115520 | COQ10B | -2.0383301 | 0.00340571 |
| ENSG00000178381 | ZFAND2A | -2.0345729 | 0.00503046 |
| ENSG00000105193 | RPS16 | -2.0333475 | 0.02582355 |
| ENSG00000164211 | STARD4 | -2.031064 | 0.04103226 |
| ENSG00000008294 | SPAG9 | -2.0306064 | 0.01396572 |
| ENSG00000143384 | MCL1 | -2.0302729 | 0.02313935 |
| ENSG00000033327 | GAB2 | -2.0297601 | 0.01533113 |
| ENSG00000241978 | AKAP2 | -2.0291565 | 0.01179582 |
| ENSG00000185947 | ZNF267 | -2.0266958 | 0.00660914 |
| ENSG00000006607 | FARP2 | -2.0249938 | 0.00572652 |
| ENSG00000083937 | CHMP2B | -2.0232671 | 0.00155933 |
| ENSG00000183735 | TBK1 | -2.0229868 | 0.00068553 |
| ENSG00000132823 | OSER1 | -2.0215438 | 0.00581841 |
| ENSG00000174738 | NR1D2 | -2.019269 | 0.00174835 |
| ENSG00000162664 | ZNF326 | -2.0174041 | 0.00088661 |
| ENSG00000263020 | AL662899.2 | -2.0171997 | 0.00303209 |
| ENSG00000100225 | FBXO7 | -2.0112347 | 0.002218 |
| ENSG00000198833 | UBE2J1 | -2.0108671 | 0.02781843 |
| ENSG00000137309 | HMGA1 | -2.0106825 | 0.01380074 |
| ENSG00000161835 | GRASP | -2.0106253 | 0.0239467 |
| ENSG00000072110 | ACTN1 | -2.0082421 | 0.02476494 |
| ENSG00000167491 | GATAD2A | -2.0076679 | 0.0126736 |
| ENSG00000113742 | CPEB4 | -2.0063751 | 0.00629665 |
| ENSG00000051108 | HERPUD1 | -2.0029501 | 0.00168749 |
| ENSG00000118503 | TNFAIP3 | -2.0020555 | 0.02209667 |
| ENSG00000100393 | EP300 | -2.0018885 | 0.00200055 |
| ENSG00000010072 | SPRTN | -2.0009042 | 0.00155933 |
| ENSG00000075415 | SLC25A3 | -1.9977476 | 0.01300954 |
| ENSG00000141551 | CSNK1D | -1.9973424 | 0.03828641 |
| ENSG00000115816 | CEBPZ | -1.9962322 | 0.00261709 |
| ENSG00000104450 | SPAG1 | -1.9958727 | 0.00940502 |
| ENSG00000140044 | JDP2 | -1.9942568 | 0.00710159 |
| ENSG00000165650 | PDZD8 | -1.9925158 | 0.01583418 |

| | | | |
|---|---|---|---|
| ENSG00000165806 | CASP7 | -1.9923525 | 0.00433347 |
| ENSG00000121749 | TBC1D15 | -1.9913267 | 0.00073864 |
| ENSG00000135655 | USP15 | -1.990058 | 0.00157557 |
| ENSG00000090339 | ICAM1 | -1.985087 | 0.0011615 |
| ENSG00000124782 | RREB1 | -1.9848633 | 0.01993667 |
| ENSG00000187522 | HSPA14 | -1.984788 | 0.00148654 |
| ENSG00000104969 | SGTA | -1.9845314 | 0.00406198 |
| ENSG00000102580 | DNAJC3 | -1.9826848 | 0.00316541 |
| ENSG00000069849 | ATP1B3 | -1.9803725 | 0.03301678 |
| ENSG00000107372 | ZFAND5 | -1.9774872 | 0.00335516 |
| ENSG00000130202 | NECTIN2 | -1.9702677 | 0.04379249 |
| ENSG00000109332 | UBE2D3 | -1.9694495 | 0.00669784 |
| ENSG00000146232 | NFKBIE | -1.9685766 | 0.00368511 |
| ENSG00000067064 | IDI1 | -1.9666984 | 0.00513019 |
| ENSG00000183484 | GPR132 | -1.964038 | 0.03115245 |
| ENSG00000137876 | RSL24D1 | -1.9635082 | 0.00791952 |
| ENSG00000131051 | RBM39 | -1.9634459 | 0.006914 |
| ENSG00000117410 | ATP6V0B | -1.9594605 | 0.00200357 |
| ENSG00000173875 | ZNF791 | -1.9582482 | 0.00563712 |
| ENSG00000108932 | SLC16A6 | -1.9581589 | 0.00509474 |
| ENSG00000114120 | SLC25A36 | -1.9569907 | 0.00328847 |
| ENSG00000119523 | ALG2 | -1.9567845 | 0.00517978 |
| ENSG00000196704 | AMZ2 | -1.9557548 | 0.03094864 |
| ENSG00000180611 | MB21D2 | -1.9555644 | 0.00194739 |
| ENSG00000163874 | ZC3H12A | -1.953556 | 0.00345283 |
| ENSG00000113811 | SELENOK | -1.9509753 | 0.00697219 |
| ENSG00000182481 | KPNA2 | -1.9417789 | 0.00454367 |
| ENSG00000139826 | ABHD13 | -1.9402662 | 0.00068553 |
| ENSG00000104064 | GABPB1 | -1.9400449 | 0.00677138 |
| ENSG00000134758 | RNF138 | -1.938487 | 0.00566992 |
| ENSG00000116786 | PLEKHM2 | -1.9376457 | 0.03167434 |
| ENSG00000140367 | UBE2Q2 | -1.9372058 | 0.00296499 |
| ENSG00000283154 | IQCJ-SCHIP1 | -1.9349819 | 0.02389596 |
| ENSG00000156273 | BACH1 | -1.9305297 | 0.00988027 |
| ENSG00000117139 | KDM5B | -1.9272034 | 0.04200804 |
| ENSG00000150991 | UBC | -1.9263115 | 0.01118305 |
| ENSG00000172845 | SP3 | -1.9239939 | 0.00058696 |
| ENSG00000111011 | RSRC2 | -1.9205933 | 0.00068553 |
| ENSG00000124766 | SOX4 | -1.9189514 | 0.02692794 |
| ENSG00000145780 | FEM1C | -1.9187959 | 0.04665951 |
| ENSG00000157954 | WIPI2 | -1.916411 | 0.00467602 |
| ENSG00000121879 | PIK3CA | -1.9146928 | 0.03693198 |

| | | | |
|---|---|---|---|
| ENSG00000132912 | DCTN4 | -1.9131747 | 0.00489427 |
| ENSG00000143761 | ARF1 | -1.9111358 | 0.00308134 |
| ENSG00000171310 | CHST11 | -1.9077359 | 0.01618175 |
| ENSG00000189376 | C8orf76 | -1.9070412 | 0.00652677 |
| ENSG00000109787 | KLF3 | -1.9062143 | 0.00418283 |
| ENSG00000159346 | ADIPOR1 | -1.9041438 | 0.00214395 |
| ENSG00000070501 | POLB | -1.9013523 | 0.00599034 |
| ENSG00000267520 | AC010733.2 | -1.9013328 | 0.00224048 |
| ENSG00000163811 | WDR43 | -1.9006545 | 0.00284279 |
| ENSG00000057757 | PITHD1 | -1.8991198 | 0.00541373 |
| ENSG00000178127 | NDUFV2 | -1.8977143 | 0.00068553 |
| ENSG00000197872 | FAM49A | -1.8976251 | 0.00642418 |
| ENSG00000158122 | AAED1 | -1.8922164 | 0.00828958 |
| ENSG00000183624 | HMCES | -1.8915489 | 0.00155927 |
| ENSG00000279766 | AC067931.1 | -1.8906824 | 0.01405517 |
| ENSG00000125037 | EMC3 | -1.890385 | 0.00541373 |
| ENSG00000123562 | MORF4L2 | -1.8875291 | 0.00391676 |
| ENSG00000156535 | CD109 | -1.8867933 | 0.00907356 |
| ENSG00000173349 | SFT2D3 | -1.8852676 | 0.00272216 |
| ENSG00000137817 | PARP6 | -1.8848749 | 0.01481879 |
| ENSG00000122257 | RBBP6 | -1.8831845 | 0.00570959 |
| ENSG00000161011 | SQSTM1 | -1.8818954 | 0.00068553 |
| ENSG00000251022 | THAP9-AS1 | -1.879013 | 0.03173612 |
| ENSG00000197619 | ZNF615 | -1.8774678 | 0.03738591 |
| ENSG00000164169 | PRMT9 | -1.8770873 | 0.00113486 |
| ENSG00000162616 | DNAJB4 | -1.8748711 | 0.00197763 |
| ENSG00000166200 | COPS2 | -1.8744975 | 0.00217078 |
| ENSG00000160789 | LMNA | -1.8728405 | 0.01500442 |
| ENSG00000128271 | ADORA2A | -1.8721744 | 0.00581841 |
| ENSG00000162924 | REL | -1.8707183 | 0.00562921 |
| ENSG00000180228 | PRKRA | -1.8703994 | 0.00068553 |
| ENSG00000138670 | RASGEF1B | -1.8692749 | 0.01681053 |
| ENSG00000170852 | KBTBD2 | -1.8689308 | 0.00541373 |
| ENSG00000130559 | CAMSAP1 | -1.8685161 | 0.04305401 |
| ENSG00000100425 | BRD1 | -1.8681931 | 0.01639526 |
| ENSG00000056972 | TRAF3IP2 | -1.8659941 | 0.00355307 |
| ENSG00000196646 | ZNF136 | -1.8650425 | 0.00764292 |
| ENSG00000168175 | MAPK1IP1L | -1.8646852 | 0.00368155 |
| ENSG00000139725 | RHOF | -1.8612864 | 0.00089939 |
| ENSG00000116670 | MAD2L2 | -1.860474 | 0.00520463 |
| ENSG00000160799 | CCDC12 | -1.8604399 | 0.00179151 |
| ENSG00000110046 | ATG2A | -1.8574903 | 0.04672487 |

| | | | |
|---|---|---|---|
| ENSG00000156232 | WHAMM | -1.8571372 | 0.0069367 |
| ENSG00000044574 | HSPA5 | -1.8566284 | 0.03098812 |
| ENSG00000156875 | MFSD14A | -1.8550645 | 0.00100563 |
| ENSG00000127824 | TUBA4A | -1.854568 | 0.0356038 |
| ENSG00000144747 | TMF1 | -1.8533699 | 0.01907077 |
| ENSG00000113575 | PPP2CA | -1.845288 | 0.00155927 |
| ENSG00000166822 | TMEM170A | -1.8428681 | 0.00028968 |
| ENSG00000114796 | KLHL24 | -1.8426384 | 0.01944808 |
| ENSG00000122406 | RPL5 | -1.8421604 | 0.00084979 |
| ENSG00000091527 | CDV3 | -1.8393821 | 0.01912849 |
| ENSG00000124688 | MAD2L1BP | -1.8379636 | 0.01436972 |
| ENSG00000232956 | SNHG15 | -1.8378725 | 0.02908505 |
| ENSG00000156502 | SUPV3L1 | -1.8376414 | 0.02406189 |
| ENSG00000008056 | SYN1 | -1.8369916 | 0.03260989 |
| ENSG00000111328 | CDK2AP1 | -1.8365447 | 0.00470819 |
| ENSG00000181555 | SETD2 | -1.8349682 | 0.01078639 |
| ENSG00000121741 | ZMYM2 | -1.8347027 | 0.00475524 |
| ENSG00000070756 | PABPC1 | -1.83213 | 0.01041945 |
| ENSG00000280138 | AC027290.2 | -1.8290481 | 0.0091546 |
| ENSG00000105849 | TWISTNB | -1.8270408 | 0.01340294 |
| ENSG00000143702 | CEP170 | -1.8267153 | 0.0250423 |
| ENSG00000067082 | KLF6 | -1.8252776 | 0.0003396 |
| ENSG00000150787 | PTS | -1.8239395 | 0.02885702 |
| ENSG00000111615 | KRR1 | -1.8237644 | 0.00277722 |
| ENSG00000149658 | YTHDF1 | -1.8222774 | 0.00068553 |
| ENSG00000106546 | AHR | -1.8195494 | 0.00370312 |
| ENSG00000119977 | TCTN3 | -1.8157545 | 0.00888906 |
| ENSG00000145779 | TNFAIP8 | -1.8131266 | 0.0141295 |
| ENSG00000170540 | ARL6IP1 | -1.8129335 | 0.00340571 |
| ENSG00000143514 | TP53BP2 | -1.8100934 | 0.01342164 |
| ENSG00000153201 | RANBP2 | -1.8072062 | 0.02608793 |
| ENSG00000133134 | BEX2 | -1.806631 | 0.0196015 |
| ENSG00000166225 | FRS2 | -1.8065146 | 0.00647532 |
| ENSG00000133112 | TPT1 | -1.8063189 | 0.00132049 |
| ENSG00000106615 | RHEB | -1.8038768 | 0.00837266 |
| ENSG00000106346 | USP42 | -1.8019186 | 0.00190335 |
| ENSG00000116560 | SFPQ | -1.7994113 | 0.00068553 |
| ENSG00000124226 | RNF114 | -1.7993923 | 0.00073864 |
| ENSG00000109670 | FBXW7 | -1.7973337 | 0.00854947 |
| ENSG00000004897 | CDC27 | -1.7965648 | 0.00261709 |
| ENSG00000114013 | CD86 | -1.7955992 | 0.01681053 |
| ENSG00000185862 | EVI2B | -1.7937298 | 0.00456187 |

| | | | |
|---|---|---|---|
| ENSG00000164758 | MED30 | -1.7926099 | 0.00068553 |
| ENSG00000196233 | LCOR | -1.7918862 | 0.00394084 |
| ENSG00000076053 | RBM7 | -1.790178 | 0.00425431 |
| ENSG00000280254 | AC233723.2 | -1.7897342 | 0.02069455 |
| ENSG00000177733 | HNRNPA0 | -1.7892427 | 0.01337033 |
| ENSG00000169826 | CSGALNACT2 | -1.7868556 | 0.0020197 |
| ENSG00000196850 | PPTC7 | -1.7814752 | 0.00205451 |
| ENSG00000108960 | MMD | -1.7806926 | 0.00622658 |
| ENSG00000114126 | TFDP2 | -1.780669 | 0.00340571 |
| ENSG00000163605 | PPP4R2 | -1.7794263 | 0.00328847 |
| ENSG00000111711 | GOLT1B | -1.7781447 | 0.00057232 |
| ENSG00000160218 | TRAPPC10 | -1.7731389 | 0.01579272 |
| ENSG00000163682 | RPL9 | -1.770584 | 0.01545309 |
| ENSG00000116030 | SUMO1 | -1.7704237 | 0.00175493 |
| ENSG00000120727 | PAIP2 | -1.7686695 | 6.3344E-05 |
| ENSG00000170881 | RNF139 | -1.7631297 | 0.03231104 |
| ENSG00000103978 | TMEM87A | -1.7616235 | 0.01498914 |
| ENSG00000105851 | PIK3CG | -1.7614506 | 0.0083323 |
| ENSG00000163788 | SNRK | -1.7602372 | 0.00272216 |
| ENSG00000110848 | CD69 | -1.7598275 | 0.02816883 |
| ENSG00000124145 | SDC4 | -1.7572618 | 0.00478686 |
| ENSG00000135241 | PNPLA8 | -1.7567401 | 0.01448055 |
| ENSG00000150977 | RILPL2 | -1.7559385 | 0.03826914 |
| ENSG00000168066 | SF1 | -1.7517346 | 0.04243611 |
| ENSG00000055208 | TAB2 | -1.7516409 | 0.02673165 |
| ENSG00000116752 | BCAS2 | -1.7512089 | 0.00270338 |
| ENSG00000105993 | DNAJB6 | -1.7510581 | 0.0028536 |
| ENSG00000150907 | FOXO1 | -1.7504125 | 0.0426935 |
| ENSG00000172062 | SMN1 | -1.747204 | 0.01832629 |
| ENSG00000204178 | TMEM57 | -1.7452922 | 0.00355307 |
| ENSG00000269968 | AC006064.4 | -1.7450836 | 0.01134212 |
| ENSG00000213079 | SCAF8 | -1.7440465 | 0.00647382 |
| ENSG00000267165 | CHMP1B-AS1 | -1.7402032 | 0.0076333 |
| ENSG00000029993 | HMGB3 | -1.73794 | 0.01056441 |
| ENSG00000133606 | MKRN1 | -1.7369882 | 0.00251682 |
| ENSG00000139505 | MTMR6 | -1.7359162 | 0.00753737 |
| ENSG00000023287 | RB1CC1 | -1.735826 | 0.00898743 |
| ENSG00000115956 | PLEK | -1.735279 | 0.02325376 |
| ENSG00000136826 | KLF4 | -1.734594 | 0.01122844 |
| ENSG00000168883 | USP39 | -1.7340665 | 0.00244907 |
| ENSG00000089818 | NECAP1 | -1.7339017 | 0.00474195 |
| ENSG00000099985 | OSM | -1.7300245 | 0.01313513 |

| | | | |
|---|---|---|---|
| ENSG00000134352 | IL6ST | -1.7297119 | 0.00557216 |
| ENSG00000054967 | RELT | -1.7292776 | 0.00985902 |
| ENSG00000165782 | PIP4P1 | -1.7284522 | 0.02069455 |
| ENSG00000048405 | ZNF800 | -1.7253481 | 0.00068553 |
| ENSG00000153914 | SREK1 | -1.724517 | 0.00758153 |
| ENSG00000160741 | CRTC2 | -1.7237738 | 0.0356038 |
| ENSG00000153922 | CHD1 | -1.7229725 | 0.00075186 |
| ENSG00000147604 | RPL7 | -1.7216834 | 0.00328847 |
| ENSG00000089234 | BRAP | -1.721121 | 0.00415779 |
| ENSG00000204977 | TRIM13 | -1.7191353 | 0.00087585 |
| ENSG00000144597 | EAF1 | -1.7172654 | 0.00292472 |
| ENSG00000155508 | CNOT8 | -1.7169155 | 0.01661933 |
| ENSG00000109113 | RAB34 | -1.714764 | 0.00170624 |
| ENSG00000162783 | IER5 | -1.7133399 | 0.0032195 |
| ENSG00000121797 | CCRL2 | -1.7130671 | 0.00303209 |
| ENSG00000205937 | RNPS1 | -1.7113322 | 0.0052756 |
| ENSG00000132334 | PTPRE | -1.7103155 | 0.03823029 |
| ENSG00000169612 | FAM103A1 | -1.7100605 | 0.00327181 |
| ENSG00000114942 | EEF1B2 | -1.7078588 | 0.00398077 |
| ENSG00000089157 | RPLP0 | -1.7073876 | 0.00803214 |
| ENSG00000168092 | PAFAH1B2 | -1.7068097 | 0.00251682 |
| ENSG00000143774 | GUK1 | -1.7052018 | 0.0160308 |
| ENSG00000111647 | UHRF1BP1L | -1.7049002 | 0.00096838 |
| ENSG00000137714 | FDX1 | -1.7047534 | 0.00152415 |
| ENSG00000147403 | RPL10 | -1.7040316 | 0.0012446 |
| ENSG00000125651 | GTF2F1 | -1.7027506 | 0.00178606 |
| ENSG00000140299 | BNIP2 | -1.7020802 | 0.00112968 |
| ENSG00000107341 | UBE2R2 | -1.7018502 | 0.01041945 |
| ENSG00000006634 | DBF4 | -1.6999703 | 0.00251622 |
| ENSG00000124201 | ZNFX1 | -1.6994888 | 0.00516637 |
| ENSG00000065978 | YBX1 | -1.6982792 | 0.00985902 |
| ENSG00000067334 | DNTTIP2 | -1.6982219 | 0.01380074 |
| ENSG00000140455 | USP3 | -1.6974653 | 0.04122249 |
| ENSG00000167004 | PDIA3 | -1.697063 | 0.00356514 |
| ENSG00000266094 | RASSF5 | -1.6924631 | 0.0089013 |
| ENSG00000119048 | UBE2B | -1.6916263 | 0.0013773 |
| ENSG00000070831 | CDC42 | -1.6915173 | 0.00151997 |
| ENSG00000136802 | LRRC8A | -1.6901455 | 0.0114427 |
| ENSG00000102753 | KPNA3 | -1.6888021 | 0.00088963 |
| ENSG00000122482 | ZNF644 | -1.6876147 | 0.01319879 |
| ENSG00000065548 | ZC3H15 | -1.6864485 | 0.0069367 |
| ENSG00000182899 | RPL35A | -1.6862025 | 0.00217078 |

| | | | |
|---|---|---|---|
| ENSG00000104765 | BNIP3L | -1.684738 | 0.00647532 |
| ENSG00000183876 | ARSI | -1.6839177 | 0.01520729 |
| ENSG00000173020 | GRK2 | -1.683088 | 0.00473792 |
| ENSG00000136807 | CDK9 | -1.6816099 | 0.01764539 |
| ENSG00000179361 | ARID3B | -1.6809762 | 0.00476653 |
| ENSG00000114125 | RNF7 | -1.6808329 | 0.00599034 |
| ENSG00000162910 | MRPL55 | -1.6802915 | 0.04284787 |
| ENSG00000156650 | KAT6B | -1.6791182 | 0.00085958 |
| ENSG00000141753 | IGFBP4 | -1.6786195 | 0.03066234 |
| ENSG00000152056 | AP1S3 | -1.6784011 | 0.04880748 |
| ENSG00000110367 | DDX6 | -1.6782528 | 0.00345296 |
| ENSG00000108175 | ZMIZ1 | -1.676447 | 0.00843969 |
| ENSG00000197021 | CXorf40B | -1.6763325 | 0.00897301 |
| ENSG00000168209 | DDIT4 | -1.6762806 | 0.00955337 |
| ENSG00000149806 | FAU | -1.6757868 | 0.0029323 |
| ENSG00000163125 | RPRD2 | -1.6743337 | 0.01278329 |
| ENSG00000241839 | PLEKHO2 | -1.6742498 | 0.00152415 |
| ENSG00000205189 | ZBTB10 | -1.6738141 | 0.00605285 |
| ENSG00000163694 | RBM47 | -1.6728334 | 0.00362829 |
| ENSG00000111640 | GAPDH | -1.6727812 | 0.00278533 |
| ENSG00000170185 | USP38 | -1.67206 | 0.00898494 |
| ENSG00000122026 | RPL21 | -1.6702654 | 0.00373954 |
| ENSG00000165006 | UBAP1 | -1.669239 | 0.01424953 |
| ENSG00000197063 | MAFG | -1.6665465 | 0.00358122 |
| ENSG00000105821 | DNAJC2 | -1.6665112 | 0.01220964 |
| ENSG00000176624 | MEX3C | -1.6639347 | 0.00365743 |
| ENSG00000179119 | SPTY2D1 | -1.6635058 | 0.02607218 |
| ENSG00000157540 | DYRK1A | -1.661791 | 0.0124965 |
| ENSG00000089737 | DDX24 | -1.6611366 | 0.00482535 |
| ENSG00000084463 | WBP11 | -1.6609711 | 0.00066481 |
| ENSG00000165527 | ARF6 | -1.6604177 | 0.00847993 |
| ENSG00000082515 | MRPL22 | -1.657823 | 0.00543849 |
| ENSG00000130803 | ZNF317 | -1.6575024 | 0.00155933 |
| ENSG00000138433 | CIR1 | -1.6543325 | 0.00992425 |
| ENSG00000172766 | NAA16 | -1.6542532 | 0.01220964 |
| ENSG00000198612 | COPS8 | -1.652776 | 0.02661358 |
| ENSG00000172239 | PAIP1 | -1.649347 | 0.00383452 |
| ENSG00000163660 | CCNL1 | -1.6462447 | 0.02582355 |
| ENSG00000188994 | ZNF292 | -1.6460525 | 0.04161115 |
| ENSG00000129484 | PARP2 | -1.6455113 | 0.00479604 |
| ENSG00000148344 | PTGES | -1.6441474 | 0.00795236 |
| ENSG00000091164 | TXNL1 | -1.6430885 | 0.02273887 |

| | | | |
|---|---|---|---|
| ENSG00000185883 | ATP6V0C | -1.6425553 | 0.04974898 |
| ENSG00000089693 | MLF2 | -1.6419361 | 0.0015584 |
| ENSG00000278311 | GGNBP2 | -1.6397228 | 0.01062296 |
| ENSG00000188229 | TUBB4B | -1.6380862 | 0.01331674 |
| ENSG00000101247 | NDUFAF5 | -1.637469 | 0.007107 |
| ENSG00000185043 | CIB1 | -1.6365419 | 0.0034608 |
| ENSG00000187109 | NAP1L1 | -1.6353027 | 0.00664721 |
| ENSG00000112031 | MTRF1L | -1.6344423 | 0.00174835 |
| ENSG00000196235 | SUPT5H | -1.6337015 | 0.01375455 |
| ENSG00000145425 | RPS3A | -1.6335388 | 0.00437009 |
| ENSG00000106608 | URGCP | -1.6330278 | 0.00151997 |
| ENSG00000115339 | GALNT3 | -1.6320953 | 0.00803214 |
| ENSG00000177374 | HIC1 | -1.630597 | 0.02117842 |
| ENSG00000137818 | RPLP1 | -1.6292029 | 0.0028804 |
| ENSG00000113615 | SEC24A | -1.6286771 | 0.02259519 |
| ENSG00000022840 | RNF10 | -1.6246667 | 0.00643481 |
| ENSG00000153561 | RMND5A | -1.6235615 | 0.00261709 |
| ENSG00000119725 | ZNF410 | -1.6234925 | 0.01184035 |
| ENSG00000137154 | RPS6 | -1.6228355 | 0.00456305 |
| ENSG00000128524 | ATP6V1F | -1.62182 | 0.00085232 |
| ENSG00000177879 | AP3S1 | -1.6213965 | 0.0254258 |
| ENSG00000071462 | BUD23 | -1.6206945 | 0.0150404 |
| ENSG00000156482 | RPL30 | -1.6202439 | 0.00454367 |
| ENSG00000163041 | H3F3A | -1.6200579 | 0.00641944 |
| ENSG00000181467 | RAP2B | -1.6196627 | 0.0173468 |
| ENSG00000080546 | SESN1 | -1.6188649 | 0.02271425 |
| ENSG00000130734 | ATG4D | -1.6179839 | 0.00089939 |
| ENSG00000099968 | BCL2L13 | -1.6155117 | 0.01939163 |
| ENSG00000134453 | RBM17 | -1.6148427 | 0.00267271 |
| ENSG00000138032 | PPM1B | -1.6147717 | 0.02622642 |
| ENSG00000198431 | TXNRD1 | -1.6147491 | 0.00264376 |
| ENSG00000144713 | RPL32 | -1.614093 | 0.00472286 |
| ENSG00000225648 | SBDSP1 | -1.6135219 | 0.00088661 |
| ENSG00000272379 | AL008729.2 | -1.612116 | 0.00375779 |
| ENSG00000124209 | RAB22A | -1.612076 | 0.02643206 |
| ENSG00000100083 | GGA1 | -1.6112479 | 0.00467105 |
| ENSG00000008952 | SEC62 | -1.6097533 | 0.00174835 |
| ENSG00000100316 | RPL3 | -1.6091778 | 0.00215171 |
| ENSG00000255198 | SNHG9 | -1.6058942 | 0.00398077 |
| ENSG00000094975 | SUCO | -1.6049018 | 0.01955516 |
| ENSG00000184014 | DENND5A | -1.6044952 | 0.03763219 |
| ENSG00000143771 | CNIH4 | -1.6030478 | 0.02227966 |

| | | | |
|---|---|---|---|
| ENSG00000095574 | IKZF5 | -1.6022235 | 0.00553015 |
| ENSG00000113580 | NR3C1 | -1.601731 | 0.00098422 |
| ENSG00000009307 | CSDE1 | -1.6014377 | 0.00136992 |
| ENSG00000179456 | ZBTB18 | -1.6012679 | 0.01112659 |
| ENSG00000112033 | PPARD | -1.5986873 | 0.0494711 |
| ENSG00000181220 | ZNF746 | -1.5982507 | 0.00606651 |
| ENSG00000172071 | EIF2AK3 | -1.5962497 | 0.01874206 |
| ENSG00000124575 | HIST1H1D | -1.5959148 | 0.04985787 |
| ENSG00000109475 | RPL34 | -1.5953481 | 0.00581898 |
| ENSG00000148572 | NRBF2 | -1.5924814 | 0.01118305 |
| ENSG00000147526 | TACC1 | -1.5924675 | 0.02927448 |
| ENSG00000156508 | EEF1A1 | -1.5924164 | 0.00690199 |
| ENSG00000178623 | GPR35 | -1.5917165 | 0.03932732 |
| ENSG00000134153 | EMC7 | -1.5912327 | 0.00230358 |
| ENSG00000162928 | PEX13 | -1.5910762 | 0.00068553 |
| ENSG00000124198 | ARFGEF2 | -1.5903358 | 0.02438605 |
| ENSG00000142534 | RPS11 | -1.5880777 | 0.00479775 |
| ENSG00000174437 | ATP2A2 | -1.5879063 | 0.00895721 |
| ENSG00000184203 | PPP1R2 | -1.5867929 | 0.00112968 |
| ENSG00000175390 | EIF3F | -1.5859656 | 0.00261709 |
| ENSG00000176142 | TMEM39A | -1.585872 | 0.00174835 |
| ENSG00000197019 | SERTAD1 | -1.5846568 | 0.00131723 |
| ENSG00000135269 | TES | -1.5840686 | 0.00478686 |
| ENSG00000198369 | SPRED2 | -1.5840124 | 0.02118998 |
| ENSG00000139697 | SBNO1 | -1.583608 | 0.0052464 |
| ENSG00000083896 | YTHDC1 | -1.5835034 | 0.00816851 |
| ENSG00000188846 | RPL14 | -1.5809511 | 0.00131723 |
| ENSG00000148926 | ADM | -1.5806023 | 0.00115627 |
| ENSG00000129351 | ILF3 | -1.5801779 | 0.04150351 |
| ENSG00000123066 | MED13L | -1.5795238 | 0.01554896 |
| ENSG00000125633 | CCDC93 | -1.5790041 | 0.02193221 |
| ENSG00000119682 | AREL1 | -1.5788068 | 0.02024358 |
| ENSG00000173960 | UBXN2A | -1.5769502 | 0.0077505 |
| ENSG00000140988 | RPS2 | -1.5765242 | 0.00180611 |
| ENSG00000069399 | BCL3 | -1.5752995 | 0.0040364 |
| ENSG00000172216 | CEBPB | -1.5741391 | 0.03113618 |
| ENSG00000198355 | PIM3 | -1.5733536 | 0.00854947 |
| ENSG00000123728 | RAP2C | -1.5728618 | 0.00188356 |
| ENSG00000071082 | RPL31 | -1.5715256 | 0.00356514 |
| ENSG00000125835 | SNRPB | -1.5710062 | 0.00584684 |
| ENSG00000168298 | HIST1H1E | -1.5709735 | 0.03816405 |
| ENSG00000086598 | TMED2 | -1.5708252 | 0.00433347 |

| | | | |
|---|---|---|---|
| ENSG00000117155 | SSX2IP | -1.5701526 | 0.00746441 |
| ENSG00000156735 | BAG4 | -1.5699471 | 0.02812648 |
| ENSG00000169100 | SLC25A6 | -1.5696587 | 0.00084979 |
| ENSG00000177954 | RPS27 | -1.5682295 | 0.00536505 |
| ENSG00000167173 | C15orf39 | -1.5673275 | 0.00145819 |
| ENSG00000148154 | UGCG | -1.5669782 | 0.01115489 |
| ENSG00000115806 | GORASP2 | -1.5656612 | 0.00533443 |
| ENSG00000175348 | TMEM9B | -1.5653281 | 0.00068553 |
| ENSG00000104388 | RAB2A | -1.5634659 | 0.00454367 |
| ENSG00000112312 | GMNN | -1.5628654 | 0.04916348 |
| ENSG00000143256 | PFDN2 | -1.5606849 | 0.01221599 |
| ENSG00000146676 | PURB | -1.5606155 | 0.03868251 |
| ENSG00000135956 | TMEM127 | -1.5603145 | 0.01043618 |
| ENSG00000173846 | PLK3 | -1.559521 | 0.00138775 |
| ENSG00000115540 | MOB4 | -1.5589761 | 0.00261709 |
| ENSG00000101084 | C20orf24 | -1.5589552 | 0.0078963 |
| ENSG00000113013 | HSPA9 | -1.5586943 | 0.00945729 |
| ENSG00000171863 | RPS7 | -1.558037 | 0.00898494 |
| ENSG00000058729 | RIOK2 | -1.5579677 | 0.02601539 |
| ENSG00000173726 | TOMM20 | -1.5566251 | 0.00752548 |
| ENSG00000188647 | PTAR1 | -1.5561199 | 0.01520729 |
| ENSG00000169567 | HINT1 | -1.5556169 | 0.00073864 |
| ENSG00000142227 | EMP3 | -1.5538489 | 0.00110322 |
| ENSG00000135486 | HNRNPA1 | -1.5535808 | 0.00068553 |
| ENSG00000265681 | RPL17 | -1.5528705 | 0.00084979 |
| ENSG00000231500 | RPS18 | -1.5528442 | 0.00803214 |
| ENSG00000111897 | SERINC1 | -1.5522296 | 0.00419863 |
| ENSG00000161526 | SAP30BP | -1.5518241 | 0.00669784 |
| ENSG00000166747 | AP1G1 | -1.550922 | 0.0048531 |
| ENSG00000132155 | RAF1 | -1.5505873 | 0.00977231 |
| ENSG00000186468 | RPS23 | -1.5496541 | 0.00384737 |
| ENSG00000134108 | ARL8B | -1.5491257 | 0.01646059 |
| ENSG00000059728 | MXD1 | -1.5487421 | 0.03200845 |
| ENSG00000164587 | RPS14 | -1.5477543 | 0.01154483 |
| ENSG00000026508 | CD44 | -1.5456982 | 0.0344454 |
| ENSG00000243147 | MRPL33 | -1.545268 | 0.00340571 |
| ENSG00000132510 | KDM6B | -1.544837 | 0.00655725 |
| ENSG00000137393 | RNF144B | -1.5439776 | 0.04668784 |
| ENSG00000112137 | PHACTR1 | -1.5429751 | 0.00174835 |
| ENSG00000167526 | RPL13 | -1.5426482 | 0.0010458 |
| ENSG00000182827 | ACBD3 | -1.5424162 | 0.00211387 |
| ENSG00000152700 | SAR1B | -1.5422892 | 0.00107147 |

| | | | |
|---|---|---|---|
| ENSG00000087460 | GNAS | -1.5407708 | 0.00266233 |
| ENSG00000165637 | VDAC2 | -1.5403825 | 0.00760268 |
| ENSG00000137815 | RTF1 | -1.540021 | 0.01007801 |
| ENSG00000115128 | SF3B6 | -1.5393517 | 0.00553015 |
| ENSG00000101367 | MAPRE1 | -1.5383761 | 0.01752544 |
| ENSG00000197958 | RPL12 | -1.5382434 | 0.0024099 |
| ENSG00000104626 | ERI1 | -1.5379124 | 0.04510053 |
| ENSG00000104267 | CA2 | -1.5370192 | 0.01999961 |
| ENSG00000198925 | ATG9A | -1.5357 | 0.01311257 |
| ENSG00000147677 | EIF3H | -1.5353354 | 0.00085958 |
| ENSG00000241343 | RPL36A | -1.5333985 | 0.00294771 |
| ENSG00000145592 | RPL37 | -1.5323838 | 0.00478686 |
| ENSG00000117614 | SYF2 | -1.5302778 | 0.00333758 |
| ENSG00000036257 | CUL3 | -1.5298442 | 0.00194739 |
| ENSG00000047634 | SCML1 | -1.5298198 | 0.02908495 |
| ENSG00000141030 | COPS3 | -1.5291822 | 0.00261709 |
| ENSG00000138231 | DBR1 | -1.5291675 | 0.00094075 |
| ENSG00000054523 | KIF1B | -1.5285471 | 0.0467038 |
| ENSG00000112242 | E2F3 | -1.5285201 | 0.00540721 |
| ENSG00000071243 | ING3 | -1.5273524 | 0.00428193 |
| ENSG00000124214 | STAU1 | -1.5271249 | 0.00138775 |
| ENSG00000171222 | SCAND1 | -1.5258209 | 0.0029729 |
| ENSG00000072364 | AFF4 | -1.5256164 | 0.00115627 |
| ENSG00000215472 | RPL17-C18orf32 | -1.5251675 | 0.00778627 |
| ENSG00000169018 | FEM1B | -1.5250244 | 0.02583179 |
| ENSG00000100997 | ABHD12 | -1.52373 | 0.00200912 |
| ENSG00000165312 | OTUD1 | -1.5229335 | 0.01913591 |
| ENSG00000198918 | RPL39 | -1.5211883 | 0.0078963 |
| ENSG00000162702 | ZNF281 | -1.520689 | 0.01220964 |
| ENSG00000008405 | CRY1 | -1.5205039 | 0.01220964 |
| ENSG00000139433 | GLTP | -1.5192312 | 0.00458456 |
| ENSG00000254772 | EEF1G | -1.5186586 | 0.00312423 |
| ENSG00000123595 | RAB9A | -1.5186059 | 0.01342519 |
| ENSG00000128016 | ZFP36 | -1.5183352 | 0.00098988 |
| ENSG00000177600 | RPLP2 | -1.5180062 | 0.0027444 |
| ENSG00000094841 | UPRT | -1.5163391 | 0.01368036 |
| ENSG00000015479 | MATR3 | -1.5153627 | 0.00107181 |
| ENSG00000242125 | SNHG3 | -1.5130102 | 0.00155927 |
| ENSG00000141428 | C18orf21 | -1.5124046 | 0.01076323 |
| ENSG00000055211 | GINM1 | -1.5119738 | 0.01087427 |
| ENSG00000175061 | LRRC75A-AS1 | -1.5090321 | 0.00774986 |

| | | | |
|---|---|---|---|
| ENSG00000198160 | MIER1 | -1.5086014 | 0.00250324 |
| ENSG00000155545 | MIER3 | -1.5083991 | 0.02110531 |
| ENSG00000185359 | HGS | -1.5074004 | 0.014456 |
| ENSG00000198242 | RPL23A | -1.5067581 | 0.00183922 |
| ENSG00000198755 | RPL10A | -1.5053093 | 0.01354723 |
| ENSG00000142541 | RPL13A | -1.5037966 | 0.00599034 |
| ENSG00000137770 | CTDSPL2 | -1.5035044 | 0.00973379 |
| ENSG00000101146 | RAE1 | -1.5029774 | 0.00898743 |
| ENSG00000170779 | CDCA4 | -1.5020035 | 0.02440814 |
| ENSG00000111666 | CHPT1 | -1.50194 | 0.00232828 |
| ENSG00000099381 | SETD1A | -1.5018804 | 0.00454404 |

#### <u>DEG CD141+ cDCs pSS *versus* HD</u>

| Transcipt ID | Gene name | Log2(FC) | FDR < 0.05 |
|---|---|---|---|
| ENSG00000110876 | SELPLG | 3.82802031 | 0.01263788 |
| ENSG00000005020 | SKAP2 | 3.68966628 | 0.01433947 |
| ENSG00000132530 | XAF1 | 3.60076426 | 0.01959496 |
| ENSG00000101347 | SAMHD1 | 3.54846982 | 0.00530244 |
| ENSG00000197713 | RPE | 3.49566248 | 0.00231608 |
| ENSG00000196975 | ANXA4 | 3.44180679 | 0.00373194 |
| ENSG00000163683 | SMIM14 | 3.41399309 | 0.00329769 |
| ENSG00000240065 | PSMB9 | 3.34825852 | 0.00771252 |
| ENSG00000177409 | SAMD9L | 3.34415245 | 0.04700928 |
| ENSG00000151702 | FLI1 | 3.26794279 | 0.0071252 |
| ENSG00000088986 | DYNLL1 | 3.20207297 | 0.00718932 |
| ENSG00000197142 | ACSL5 | 3.18105019 | 0.01686697 |
| ENSG00000131203 | IDO1 | 3.17237199 | 0.04053815 |
| ENSG00000131844 | MCCC2 | 3.08943003 | 0.00809412 |
| ENSG00000117676 | RPS6KA1 | 3.08369812 | 0.00373378 |
| ENSG00000178927 | C17orf62 | 3.01908121 | 0.00231608 |
| ENSG00000178175 | ZNF366 | 3.00669325 | 0.01156951 |
| ENSG00000175567 | UCP2 | 2.97576587 | 0.00942001 |
| ENSG00000167797 | CDK2AP2 | 2.97169671 | 0.00695581 |
| ENSG00000092964 | DPYSL2 | 2.9662081 | 0.01733544 |
| ENSG00000133835 | HSD17B4 | 2.95560908 | 0.00373194 |
| ENSG00000004455 | AK2 | 2.88436107 | 0.00450671 |
| ENSG00000184432 | COPB2 | 2.85168951 | 0.00373194 |
| ENSG00000073849 | ST6GAL1 | 2.84253331 | 0.04936695 |
| ENSG00000108946 | PRKAR1A | 2.82726055 | 0.00785938 |
| ENSG00000157601 | MX1 | 2.81221268 | 0.00643515 |
| ENSG00000117054 | ACADM | 2.8098934 | 0.01038122 |
| ENSG00000133943 | DGLUCY | 2.80765017 | 0.00939014 |
| ENSG00000111335 | OAS2 | 2.79151142 | 0.02409658 |
| ENSG00000118855 | MFSD1 | 2.77966236 | 0.00231608 |
| ENSG00000111913 | RIPOR2 | 2.77926568 | 0.0164224 |
| ENSG00000188404 | SELL | 2.7711235 | 0.00683094 |
| ENSG00000112118 | MCM3 | 2.75448581 | 0.04775931 |
| ENSG00000127951 | FGL2 | 2.70464858 | 0.00471655 |
| ENSG00000165672 | PRDX3 | 2.68982116 | 0.018607 |
| ENSG00000136518 | ACTL6A | 2.6770965 | 0.03205559 |

| | | | |
|---|---|---|---|
| ENSG00000133106 | EPSTI1 | 2.6728163 | 0.00427995 |
| ENSG00000143390 | RFX5 | 2.64713348 | 0.00411253 |
| ENSG00000103381 | CPPED1 | 2.64591863 | 0.02817193 |
| ENSG00000116455 | WDR77 | 2.62619068 | 0.00939014 |
| ENSG00000113845 | TIMMDC1 | 2.61924211 | 0.00498634 |
| ENSG00000170581 | STAT2 | 2.61603253 | 0.01890536 |
| ENSG00000217555 | CKLF | 2.61565748 | 0.01017501 |
| ENSG00000138459 | SLC35A5 | 2.61037387 | 0.00574457 |
| ENSG00000093072 | ADA2 | 2.61031333 | 0.01118101 |
| ENSG00000105483 | CARD8 | 2.58053041 | 0.00366462 |
| ENSG00000146192 | FGD2 | 2.57459956 | 0.03185424 |
| ENSG00000214114 | MYCBP | 2.54384693 | 0.0123091 |
| ENSG00000111481 | COPZ1 | 2.53491668 | 0.00092215 |
| ENSG00000103018 | CYB5B | 2.53412838 | 0.03940052 |
| ENSG00000115415 | STAT1 | 2.51436658 | 0.00714668 |
| ENSG00000103423 | DNAJA3 | 2.48171181 | 0.00369871 |
| ENSG00000133739 | LRRCC1 | 2.47022279 | 0.00427995 |
| ENSG00000058668 | ATP2B4 | 2.46603127 | 0.01140717 |
| ENSG00000136631 | VPS45 | 2.4455971 | 0.00669336 |
| ENSG00000101336 | HCK | 2.43572212 | 0.00414186 |
| ENSG00000165934 | CPSF2 | 2.42895524 | 0.02319321 |
| ENSG00000166801 | FAM111A | 2.42695116 | 0.04149208 |
| ENSG00000170854 | RIOX2 | 2.42306435 | 0.01017501 |
| ENSG00000082074 | FYB1 | 2.40640034 | 0.01889942 |
| ENSG00000166002 | SMCO4 | 2.39157552 | 0.02742631 |
| ENSG00000174123 | TLR10 | 2.39131619 | 0.01831296 |
| ENSG00000197471 | SPN | 2.38349931 | 0.01377676 |
| ENSG00000132256 | TRIM5 | 2.37573124 | 0.02259125 |
| ENSG00000174500 | GCSAM | 2.36971573 | 0.02056281 |
| ENSG00000162736 | NCSTN | 2.36845665 | 0.01972577 |
| ENSG00000155097 | ATP6V1C1 | 2.36729947 | 0.04652814 |
| ENSG00000183486 | MX2 | 2.3666934 | 0.00366462 |
| ENSG00000181192 | DHTKD1 | 2.36357046 | 0.00643515 |
| ENSG00000154511 | FAM69A | 2.35841198 | 0.02481535 |
| ENSG00000113522 | RAD50 | 2.35497808 | 0.00375346 |
| ENSG00000109684 | CLNK | 2.34232203 | 0.01516728 |
| ENSG00000107551 | RASSF4 | 2.33187847 | 0.02021428 |
| ENSG00000161929 | SCIMP | 2.32602311 | 0.01801069 |
| ENSG00000133706 | LARS | 2.32397223 | 0.00643515 |
| ENSG00000145781 | COMMD10 | 2.32140285 | 0.02056247 |

| | | | |
|---|---|---|---|
| ENSG00000188554 | NBR1 | 2.31683543 | 0.02110815 |
| ENSG00000134910 | STT3A | 2.31416843 | 0.03073439 |
| ENSG00000197043 | ANXA6 | 2.30138687 | 0.01333465 |
| ENSG00000100442 | FKBP3 | 2.30137712 | 0.03950432 |
| ENSG00000198951 | NAGA | 2.30017514 | 0.01778545 |
| ENSG00000178537 | SLC25A20 | 2.29976075 | 0.00683094 |
| ENSG00000145907 | G3BP1 | 2.28657148 | 0.00366462 |
| ENSG00000159131 | GART | 2.27895148 | 0.02160911 |
| ENSG00000137513 | NARS2 | 2.27413426 | 0.0164287 |
| ENSG00000102699 | PARP4 | 2.26160551 | 0.02565202 |
| ENSG00000108798 | ABI3 | 2.26085742 | 0.0417526 |
| ENSG00000093144 | ECHDC1 | 2.25935241 | 0.01346053 |
| ENSG00000166797 | FAM96A | 2.25395611 | 0.01382958 |
| ENSG00000028528 | SNX1 | 2.2481123 | 0.00366462 |
| ENSG00000138073 | PREB | 2.23899359 | 0.00427995 |
| ENSG00000196511 | TPK1 | 2.23484766 | 0.00853143 |
| ENSG00000145982 | FARS2 | 2.23187495 | 0.01599822 |
| ENSG00000197943 | PLCG2 | 2.2254161 | 0.01382958 |
| ENSG00000169220 | RGS14 | 2.22529372 | 0.00471655 |
| ENSG00000143252 | SDHC | 2.22432491 | 0.00369871 |
| ENSG00000168310 | IRF2 | 2.21941679 | 0.01142001 |
| ENSG00000138185 | ENTPD1 | 2.21332138 | 0.01889942 |
| ENSG00000115204 | MPV17 | 2.21312672 | 0.00910123 |
| ENSG00000258659 | TRIM34 | 2.20303708 | 0.02833544 |
| ENSG00000136100 | VPS36 | 2.20130245 | 0.04311957 |
| ENSG00000068079 | IFI35 | 2.20055085 | 0.00471655 |
| ENSG00000028116 | VRK2 | 2.19526545 | 0.00366462 |
| ENSG00000138660 | AP1AR | 2.18304632 | 0.02083863 |
| ENSG00000136279 | DBNL | 2.1820547 | 0.00897674 |
| ENSG00000103051 | COG4 | 2.18108613 | 0.01480602 |
| ENSG00000175792 | RUVBL1 | 2.18096182 | 0.01382958 |
| ENSG00000170915 | PAQR8 | 2.17855823 | 0.03259356 |
| ENSG00000159228 | CBR1 | 2.17642933 | 0.01778545 |
| ENSG00000198771 | RCSD1 | 2.15975098 | 0.03365728 |
| ENSG00000065413 | ANKRD44 | 2.14995458 | 0.00414186 |
| ENSG00000137509 | PRCP | 2.14767494 | 0.02417588 |
| ENSG00000180353 | HCLS1 | 2.14375858 | 0.00795821 |
| ENSG00000140968 | IRF8 | 2.1415942 | 0.01040756 |
| ENSG00000110063 | DCPS | 2.13097937 | 0.01065996 |
| ENSG00000134152 | KATNBL1 | 2.12442073 | 0.002059 |

| | | | |
|---|---|---|---|
| ENSG00000026751 | SLAMF7 | 2.122695 | 0.00414186 |
| ENSG00000128513 | POT1 | 2.11684276 | 0.03724185 |
| ENSG00000153064 | BANK1 | 2.11592711 | 0.01948002 |
| ENSG00000166326 | TRIM44 | 2.10872196 | 0.03214462 |
| ENSG00000176087 | SLC35A4 | 2.10613373 | 0.00783832 |
| ENSG00000159063 | ALG8 | 2.09621344 | 0.00772963 |
| ENSG00000072501 | SMC1A | 2.0873365 | 0.00388963 |
| ENSG00000130021 | PUDP | 2.08334842 | 0.02254971 |
| ENSG00000133313 | CNDP2 | 2.08310086 | 0.00674003 |
| ENSG00000182511 | FES | 2.08124765 | 0.00585693 |
| ENSG00000090863 | GLG1 | 2.07175674 | 0.00686317 |
| ENSG00000006715 | VPS41 | 2.06786491 | 0.0058823 |
| ENSG00000123106 | CCDC91 | 2.06575858 | 0.01849331 |
| ENSG00000149084 | HSD17B12 | 2.06249852 | 0.00329769 |
| ENSG00000014138 | POLA2 | 2.05890883 | 0.03087971 |
| ENSG00000138768 | USO1 | 2.05724479 | 0.01698373 |
| ENSG00000155660 | PDIA4 | 2.05661756 | 0.01046554 |
| ENSG00000067704 | IARS2 | 2.05190677 | 0.00683094 |
| ENSG00000167220 | HDHD2 | 2.05170364 | 0.0074401 |
| ENSG00000170876 | TMEM43 | 2.05022357 | 0.01202718 |
| ENSG00000139641 | ESYT1 | 2.04007922 | 0.00706653 |
| ENSG00000159445 | THEM4 | 2.03370264 | 0.00238242 |
| ENSG00000162704 | ARPC5 | 2.031856 | 0.00621715 |
| ENSG00000243943 | ZNF512 | 2.02404075 | 0.01764812 |
| ENSG00000075239 | ACAT1 | 2.02336357 | 0.00678127 |
| ENSG00000164308 | ERAP2 | 2.00102837 | 0.01747714 |
| ENSG00000102524 | TNFSF13B | 1.99467751 | 0.02972242 |
| ENSG00000014216 | CAPN1 | 1.99448283 | 0.00798766 |
| ENSG00000087263 | OGFOD1 | 1.99421872 | 0.01403954 |
| ENSG00000255833 | TIFAB | 1.99346242 | 0.00212779 |
| ENSG00000138758 | SEPT11 | 1.99278602 | 0.0159471 |
| ENSG00000135317 | SNX14 | 1.99277301 | 0.02398012 |
| ENSG00000134996 | OSTF1 | 1.98817748 | 0.04949211 |
| ENSG00000156587 | UBE2L6 | 1.98563628 | 0.01659642 |
| ENSG00000166888 | STAT6 | 1.98342839 | 0.00366462 |
| ENSG00000128340 | RAC2 | 1.98281961 | 0.0291451 |
| ENSG00000103544 | C16orf62 | 1.97865123 | 0.04701596 |
| ENSG00000130429 | ARPC1B | 1.96710688 | 0.0060948 |
| ENSG00000088035 | ALG6 | 1.9664866 | 0.03133879 |
| ENSG00000143314 | MRPL24 | 1.9661189 | 0.02259125 |

| | | | |
|---|---|---|---|
| ENSG00000137845 | ADAM10 | 1.96078759 | 0.0239978 |
| ENSG00000128915 | ICE2 | 1.95925998 | 0.0053814 |
| ENSG00000177054 | ZDHHC13 | 1.95914759 | 0.018607 |
| ENSG00000178057 | NDUFAF3 | 1.95822904 | 0.00834834 |
| ENSG00000035687 | ADSS | 1.95778237 | 0.01565564 |
| ENSG00000084733 | RAB10 | 1.95772064 | 0.00366462 |
| ENSG00000115234 | SNX17 | 1.95654784 | 0.00450671 |
| ENSG00000147905 | ZCCHC7 | 1.94631911 | 0.03575874 |
| ENSG00000102710 | SUPT20H | 1.94521725 | 0.00471655 |
| ENSG00000272047 | GTF2H5 | 1.94299036 | 0.00748852 |
| ENSG00000164062 | APEH | 1.94196538 | 0.02589679 |
| ENSG00000198736 | MSRB1 | 1.93727756 | 0.01216395 |
| ENSG00000136874 | STX17 | 1.93569262 | 0.00373378 |
| ENSG00000112367 | FIG4 | 1.93328907 | 0.0499912 |
| ENSG00000119203 | CPSF3 | 1.93234535 | 0.03489201 |
| ENSG00000108651 | UTP6 | 1.93055903 | 0.02083891 |
| ENSG00000109814 | UGDH | 1.9275224 | 0.03064301 |
| ENSG00000120925 | RNF170 | 1.92584144 | 0.01371457 |
| ENSG00000159596 | TMEM69 | 1.91866781 | 0.01698373 |
| ENSG00000169116 | PARM1 | 1.91831262 | 0.01283324 |
| ENSG00000080189 | SLC35C2 | 1.91532369 | 0.02259125 |
| ENSG00000141510 | TP53 | 1.91435441 | 0.02477928 |
| ENSG00000138363 | ATIC | 1.9089075 | 0.01065996 |
| ENSG00000116133 | DHCR24 | 1.90747385 | 0.04834494 |
| ENSG00000183597 | TANGO2 | 1.90622629 | 0.03474393 |
| ENSG00000121691 | CAT | 1.90505411 | 0.00366462 |
| ENSG00000121281 | ADCY7 | 1.90312789 | 0.01321415 |
| ENSG00000101464 | PIGU | 1.89949586 | 0.00498718 |
| ENSG00000167272 | POP5 | 1.89909548 | 0.04297077 |
| ENSG00000162600 | OMA1 | 1.89457122 | 0.00563522 |
| ENSG00000185973 | TMLHE | 1.89071014 | 0.02052985 |
| ENSG00000256043 | CTSO | 1.89027423 | 0.03288142 |
| ENSG00000197157 | SND1 | 1.88915776 | 0.02428243 |
| ENSG00000032389 | EIPR1 | 1.87940006 | 0.01208677 |
| ENSG00000119321 | FKBP15 | 1.87712459 | 0.00538725 |
| ENSG00000181924 | COA4 | 1.87138696 | 0.04404324 |
| ENSG00000070269 | TMEM260 | 1.87022464 | 0.02813121 |
| ENSG00000155957 | TMBIM4 | 1.86478753 | 0.01382958 |
| ENSG00000138246 | DNAJC13 | 1.86384869 | 0.04149208 |
| ENSG00000090861 | AARS | 1.85873057 | 0.01863023 |

| | | | |
|---|---|---|---|
| ENSG00000131236 | CAP1 | 1.85861215 | 0.00366462 |
| ENSG00000004468 | CD38 | 1.85798611 | 0.00715816 |
| ENSG00000168385 | SEPT2 | 1.85482962 | 0.00505093 |
| ENSG00000110435 | PDHX | 1.85051266 | 0.0159471 |
| ENSG00000108984 | MAP2K6 | 1.84232405 | 0.0088983 |
| ENSG00000122694 | GLIPR2 | 1.83979317 | 0.00623347 |
| ENSG00000228223 | HCG11 | 1.8396572 | 0.04682276 |
| ENSG00000172775 | FAM192A | 1.83064498 | 0.03279199 |
| ENSG00000138796 | HADH | 1.82793971 | 0.00373378 |
| ENSG00000184992 | BRI3BP | 1.82782933 | 0.03259356 |
| ENSG00000174243 | DDX23 | 1.82695117 | 0.01920078 |
| ENSG00000135336 | ORC3 | 1.82458007 | 0.02598878 |
| ENSG00000196189 | SEMA4A | 1.82371466 | 0.04199847 |
| ENSG00000000938 | FGR | 1.82314394 | 0.00782979 |
| ENSG00000170006 | TMEM154 | 1.8145082 | 0.00695581 |
| ENSG00000085871 | MGST2 | 1.81235608 | 0.02052985 |
| ENSG00000110768 | GTF2H1 | 1.8076399 | 0.04916572 |
| ENSG00000143624 | INTS3 | 1.80407194 | 0.01491734 |
| ENSG00000100938 | GMPR2 | 1.80191394 | 0.00718932 |
| ENSG00000122033 | MTIF3 | 1.79769178 | 0.03842738 |
| ENSG00000139278 | GLIPR1 | 1.79725051 | 0.02046969 |
| ENSG00000128654 | MTX2 | 1.79453089 | 0.01831296 |
| ENSG00000213983 | AP1G2 | 1.79205768 | 0.00771252 |
| ENSG00000136270 | TBRG4 | 1.79087807 | 0.01740138 |
| ENSG00000136485 | DCAF7 | 1.79043521 | 0.03743491 |
| ENSG00000205765 | C5orf51 | 1.78837985 | 0.04794577 |
| ENSG00000181804 | SLC9A9 | 1.78735554 | 0.01740138 |
| ENSG00000165795 | NDRG2 | 1.78565459 | 0.03860348 |
| ENSG00000176102 | CSTF3 | 1.78210992 | 0.00498634 |
| ENSG00000108528 | SLC25A11 | 1.7809832 | 0.0225146 |
| ENSG00000138964 | PARVG | 1.77976448 | 0.00783112 |
| ENSG00000077232 | DNAJC10 | 1.77674394 | 0.03956014 |
| ENSG00000182247 | UBE2E2 | 1.77383984 | 0.02213213 |
| ENSG00000120860 | WASHC3 | 1.77325401 | 0.04700928 |
| ENSG00000152683 | SLC30A6 | 1.76954512 | 0.00642013 |
| ENSG00000168906 | MAT2A | 1.76732256 | 0.0184155 |
| ENSG00000083535 | PIBF1 | 1.76639128 | 0.01986989 |
| ENSG00000134215 | VAV3 | 1.76456928 | 0.04485765 |
| ENSG00000131374 | TBC1D5 | 1.7600876 | 0.0070359 |
| ENSG00000143106 | PSMA5 | 1.75667886 | 0.00518293 |

| | | | |
|---|---|---|---|
| ENSG00000171307 | ZDHHC16 | 1.75581291 | 0.0184155 |
| ENSG00000089063 | TMEM230 | 1.75554914 | 0.02435736 |
| ENSG00000188452 | CERKL | 1.75073598 | 0.01764523 |
| ENSG00000073969 | NSF | 1.74689502 | 0.04337462 |
| ENSG00000006744 | ELAC2 | 1.7451714 | 0.03815034 |
| ENSG00000138071 | ACTR2 | 1.74218255 | 0.00471655 |
| ENSG00000092531 | SNAP23 | 1.74089869 | 0.02373064 |
| ENSG00000078618 | NRDC | 1.73547728 | 0.04120458 |
| ENSG00000149089 | APIP | 1.72848666 | 0.00695581 |
| ENSG00000117632 | STMN1 | 1.72686 | 0.02567634 |
| ENSG00000177628 | GBA | 1.72678781 | 0.04466065 |
| ENSG00000273749 | CYFIP1 | 1.72137066 | 0.04149208 |
| ENSG00000138413 | IDH1 | 1.71448061 | 0.01517814 |
| ENSG00000131652 | THOC6 | 1.70927481 | 0.00718932 |
| ENSG00000104325 | DECR1 | 1.7063217 | 0.00086792 |
| ENSG00000113312 | TTC1 | 1.70394262 | 0.0102919 |
| ENSG00000136628 | EPRS | 1.70315632 | 0.037056 |
| ENSG00000088451 | TGDS | 1.70293082 | 0.04806862 |
| ENSG00000182551 | ADI1 | 1.69931137 | 0.00329769 |
| ENSG00000096968 | JAK2 | 1.69650899 | 0.04491806 |
| ENSG00000131828 | PDHA1 | 1.69645807 | 0.00471655 |
| ENSG00000140395 | WDR61 | 1.69609591 | 0.03676234 |
| ENSG00000122986 | HVCN1 | 1.68983788 | 0.04685849 |
| ENSG00000104812 | GYS1 | 1.6895187 | 0.00548871 |
| ENSG00000110108 | TMEM109 | 1.6848329 | 0.01936255 |
| ENSG00000031081 | ARHGAP31 | 1.68019503 | 0.00718932 |
| ENSG00000188321 | ZNF559 | 1.67943891 | 0.04733928 |
| ENSG00000145348 | TBCK | 1.6758879 | 0.04543213 |
| ENSG00000141959 | PFKL | 1.67301115 | 0.01734575 |
| ENSG00000088205 | DDX18 | 1.66292701 | 0.02034028 |
| ENSG00000141349 | G6PC3 | 1.65694636 | 0.00509902 |
| ENSG00000249915 | PDCD6 | 1.65438062 | 0.01607135 |
| ENSG00000094914 | AAAS | 1.65321137 | 0.02178995 |
| ENSG00000148634 | HERC4 | 1.65091532 | 0.04120458 |
| ENSG00000241553 | ARPC4 | 1.64852211 | 0.00471655 |
| ENSG00000101246 | ARFRP1 | 1.6474074 | 0.01498374 |
| ENSG00000014641 | MDH1 | 1.64713785 | 0.01184988 |
| ENSG00000204619 | PPP1R11 | 1.64151227 | 0.00751565 |
| ENSG00000117280 | RAB29 | 1.64123066 | 0.02513578 |
| ENSG00000109911 | ELP4 | 1.63994861 | 0.04936695 |

| | | | |
|---|---|---|---|
| ENSG00000127946 | HIP1 | 1.63988273 | 0.02095779 |
| ENSG00000114956 | DGUOK | 1.62764348 | 0.03247044 |
| ENSG00000146376 | ARHGAP18 | 1.62659247 | 0.00236932 |
| ENSG00000169019 | COMMD8 | 1.61957748 | 0.02882527 |
| ENSG00000156110 | ADK | 1.61949922 | 0.00366462 |
| ENSG00000162813 | BPNT1 | 1.61931129 | 0.01918626 |
| ENSG00000147471 | PLPBP | 1.61714025 | 0.01972577 |
| ENSG00000120053 | GOT1 | 1.61486095 | 0.00414186 |
| ENSG00000198814 | GK | 1.61386071 | 0.00771252 |
| ENSG00000066583 | ISOC1 | 1.61272735 | 0.01836102 |
| ENSG00000105676 | ARMC6 | 1.61269789 | 0.00709631 |
| ENSG00000163346 | PBXIP1 | 1.61145872 | 0.02274656 |
| ENSG00000156973 | PDE6D | 1.60933264 | 0.0108102 |
| ENSG00000125703 | ATG4C | 1.6090627 | 0.02248772 |
| ENSG00000124702 | KLHDC3 | 1.6080931 | 0.00373194 |
| ENSG00000108622 | ICAM2 | 1.60770762 | 0.00809412 |
| ENSG00000100201 | DDX17 | 1.60554428 | 0.04949211 |
| ENSG00000182179 | UBA7 | 1.60384845 | 0.0287894 |
| ENSG00000187688 | TRPV2 | 1.60332056 | 0.00798925 |
| ENSG00000149196 | HIKESHI | 1.59916945 | 0.01636258 |
| ENSG00000162946 | DISC1 | 1.59411755 | 0.00785938 |
| ENSG00000088827 | SIGLEC1 | 1.59338331 | 0.02399459 |
| ENSG00000162735 | PEX19 | 1.59181511 | 0.04044427 |
| ENSG00000129691 | ASH2L | 1.59058308 | 0.02886473 |
| ENSG00000168394 | TAP1 | 1.58770237 | 0.03148322 |
| ENSG00000132182 | NUP210 | 1.58555771 | 0.0473165 |
| ENSG00000175938 | ORAI3 | 1.58502623 | 0.00373194 |
| ENSG00000134452 | FBXO18 | 1.58402095 | 0.00906886 |
| ENSG00000196576 | PLXNB2 | 1.58273003 | 0.04480124 |
| ENSG00000099899 | TRMT2A | 1.58232477 | 0.02820495 |
| ENSG00000125877 | ITPA | 1.58025065 | 0.00373194 |
| ENSG00000170248 | PDCD6IP | 1.57926176 | 0.0080564 |
| ENSG00000102316 | MAGED2 | 1.57519043 | 0.00897674 |
| ENSG00000136271 | DDX56 | 1.57485191 | 0.00892823 |
| ENSG00000119616 | FCF1 | 1.57422682 | 0.01778545 |
| ENSG00000147592 | LACTB2 | 1.57350347 | 0.01491734 |
| ENSG00000109572 | CLCN3 | 1.57237796 | 0.03724185 |
| ENSG00000108389 | MTMR4 | 1.57116178 | 0.00885073 |
| ENSG00000138629 | UBL7 | 1.57055584 | 0.03327519 |
| ENSG00000100714 | MTHFD1 | 1.56665968 | 0.03205559 |

| | | | |
|---|---|---|---|
| ENSG00000135457 | TFCP2 | 1.56555175 | 0.002059 |
| ENSG00000129480 | DTD2 | 1.56436779 | 0.02044521 |
| ENSG00000160310 | PRMT2 | 1.56277539 | 0.04787676 |
| ENSG00000108479 | GALK1 | 1.56201279 | 0.00706653 |
| ENSG00000100395 | L3MBTL2 | 1.56123945 | 0.01666316 |
| ENSG00000160213 | CSTB | 1.56071938 | 0.01530022 |
| ENSG00000140262 | TCF12 | 1.55900951 | 0.00366462 |
| ENSG00000223501 | VPS52 | 1.55486214 | 0.0069917 |
| ENSG00000157637 | SLC38A10 | 1.55228723 | 0.00808263 |
| ENSG00000198252 | STYX | 1.55066317 | 0.00373194 |
| ENSG00000180370 | PAK2 | 1.54978257 | 0.00678127 |
| ENSG00000163344 | PMVK | 1.54958534 | 0.04949211 |
| ENSG00000124508 | BTN2A2 | 1.54923435 | 0.01040756 |
| ENSG00000204261 | PSMB8-AS1 | 1.54647747 | 0.01382796 |
| ENSG00000165609 | NUDT5 | 1.5464016 | 0.01920078 |
| ENSG00000077380 | DYNC1I2 | 1.54433065 | 0.00719913 |
| ENSG00000135624 | CCT7 | 1.54087321 | 0.01530022 |
| ENSG00000013364 | MVP | 1.53339149 | 0.01298825 |
| ENSG00000110871 | COQ5 | 1.53185169 | 0.01698373 |
| ENSG00000131473 | ACLY | 1.53045419 | 0.02615942 |
| ENSG00000129317 | PUS7L | 1.52568904 | 0.04483242 |
| ENSG00000134851 | TMEM165 | 1.52339767 | 0.00678127 |
| ENSG00000033050 | ABCF2 | 1.52272164 | 0.04794577 |
| ENSG00000241468 | ATP5J2 | 1.52189757 | 0.0095478 |
| ENSG00000112079 | STK38 | 1.5184563 | 0.04133962 |
| ENSG00000223960 | AC009948.1 | 1.51472539 | 0.00719913 |
| ENSG00000085491 | SLC25A24 | 1.51410362 | 0.037056 |
| ENSG00000165476 | REEP3 | 1.5113931 | 0.03067089 |
| ENSG00000150753 | CCT5 | 1.50788036 | 0.01638433 |
| ENSG00000102309 | PIN4 | 1.50580323 | 0.04940805 |
| ENSG00000171155 | C1GALT1C1 | 1.50396023 | 0.01677843 |
| ENSG00000160551 | TAOK1 | 1.50351125 | 0.04395127 |
| ENSG00000163154 | TNFAIP8L2 | 1.50292149 | 0.01607135 |
| ENSG00000127948 | POR | 1.50205485 | 0.0473165 |
| ENSG00000186594 | MIR22HG | -4.1138605 | 0.037056 |
| ENSG00000169895 | SYAP1 | -3.6935282 | 0.04888382 |
| ENSG00000175197 | DDIT3 | -3.6292117 | 0.01786554 |
| ENSG00000104312 | RIPK2 | -3.5049456 | 0.02687948 |
| ENSG00000131669 | NINJ1 | -3.4464943 | 0.0095529 |
| ENSG00000171988 | JMJD1C | -3.4364109 | 0.02222424 |

| | | | |
|---|---|---|---|
| ENSG00000120690 | ELF1 | -3.3945734 | 0.02594793 |
| ENSG00000169508 | GPR183 | -3.3343572 | 0.02220983 |
| ENSG00000154640 | BTG3 | -3.3100222 | 0.01341595 |
| ENSG00000277117 | FP565260.3 | -3.2554304 | 0.03348651 |
| ENSG00000160223 | ICOSLG | -3.2221071 | 0.01516728 |
| ENSG00000011422 | PLAUR | -3.1857979 | 0.02259125 |
| ENSG00000114784 | EIF1B | -3.1774242 | 0.01366317 |
| ENSG00000165233 | CARD19 | -3.0676944 | 0.00771252 |
| ENSG00000120705 | ETF1 | -3.0238314 | 0.02922772 |
| ENSG00000099860 | GADD45B | -2.9778697 | 0.02435736 |
| ENSG00000130340 | SNX9 | -2.9684615 | 0.03585533 |
| ENSG00000196233 | LCOR | -2.9437394 | 0.02052985 |
| ENSG00000197170 | PSMD12 | -2.8873085 | 0.0159471 |
| ENSG00000173812 | EIF1 | -2.8629371 | 0.01118101 |
| ENSG00000169905 | TOR1AIP2 | -2.845181 | 0.02109402 |
| ENSG00000272886 | DCP1A | -2.8334997 | 0.00414186 |
| ENSG00000025156 | HSF2 | -2.8308445 | 0.00136833 |
| ENSG00000143153 | ATP1B1 | -2.7778394 | 0.00509326 |
| ENSG00000184205 | TSPYL2 | -2.7635336 | 0.0276593 |
| ENSG00000068697 | LAPTM4A | -2.742152 | 0.01921875 |
| ENSG00000143507 | DUSP10 | -2.7355762 | 0.03535037 |
| ENSG00000146457 | WTAP | -2.735327 | 0.01746478 |
| ENSG00000110852 | CLEC2B | -2.7079704 | 0.00373194 |
| ENSG00000151012 | SLC7A11 | -2.705993 | 0.02775042 |
| ENSG00000150907 | FOXO1 | -2.6787004 | 0.01662887 |
| ENSG00000121966 | CXCR4 | -2.676161 | 0.0414865 |
| ENSG00000067082 | KLF6 | -2.671159 | 0.03034903 |
| ENSG00000134107 | BHLHE40 | -2.6703527 | 0.03259356 |
| ENSG00000116044 | NFE2L2 | -2.6667114 | 0.04483242 |
| ENSG00000113070 | HBEGF | -2.657094 | 0.01651978 |
| ENSG00000138069 | RAB1A | -2.6237485 | 0.02565202 |
| ENSG00000171522 | PTGER4 | -2.6039809 | 0.02407448 |
| ENSG00000104765 | BNIP3L | -2.6025565 | 0.01209023 |
| ENSG00000119326 | CTNNAL1 | -2.5966732 | 0.00812086 |
| ENSG00000114796 | KLHL24 | -2.5933178 | 0.00943543 |
| ENSG00000135655 | USP15 | -2.5879499 | 0.01686697 |
| ENSG00000277632 | CCL3 | -2.5855865 | 0.03558605 |
| ENSG00000118308 | LRMP | -2.5694642 | 0.0291451 |
| ENSG00000173575 | CHD2 | -2.5502927 | 0.03262661 |
| ENSG00000267520 | AC010733.2 | -2.5416381 | 0.01944561 |

| | | | |
|---|---|---|---|
| ENSG00000120129 | DUSP1 | -2.5176534 | 0.01849331 |
| ENSG00000136950 | ARPC5L | -2.4969028 | 0.00848138 |
| ENSG00000185477 | GPRIN3 | -2.4953763 | 0.00538725 |
| ENSG00000162924 | REL | -2.4769513 | 0.00366462 |
| ENSG00000166822 | TMEM170A | -2.474987 | 0.03035108 |
| ENSG00000124782 | RREB1 | -2.4708018 | 0.00498718 |
| ENSG00000120616 | EPC1 | -2.4696606 | 0.00719913 |
| ENSG00000117000 | RLF | -2.4630327 | 0.02304511 |
| ENSG00000111615 | KRR1 | -2.4532643 | 0.01803367 |
| ENSG00000107937 | GTPBP4 | -2.4521505 | 0.01700522 |
| ENSG00000163961 | RNF168 | -2.4425438 | 0.01530022 |
| ENSG00000185650 | ZFP36L1 | -2.436076 | 0.00373378 |
| ENSG00000089818 | NECAP1 | -2.4218155 | 0.01118014 |
| ENSG00000150977 | RILPL2 | -2.4151521 | 0.04793703 |
| ENSG00000132952 | USPL1 | -2.4128638 | 0.02471595 |
| ENSG00000013441 | CLK1 | -2.4099282 | 0.00574457 |
| ENSG00000272888 | LINC01578 | -2.3985687 | 0.0071252 |
| ENSG00000139977 | NAA30 | -2.3978653 | 0.00373194 |
| ENSG00000105821 | DNAJC2 | -2.3925275 | 0.01118101 |
| ENSG00000115165 | CYTIP | -2.3875843 | 0.03287197 |
| ENSG00000144674 | GOLGA4 | -2.3827561 | 0.00427995 |
| ENSG00000081320 | STK17B | -2.3761399 | 0.03823614 |
| ENSG00000011566 | MAP4K3 | -2.3741461 | 0.01565564 |
| ENSG00000128272 | ATF4 | -2.367223 | 0.0102919 |
| ENSG00000133134 | BEX2 | -2.3659944 | 0.00231608 |
| ENSG00000135404 | CD63 | -2.3636386 | 0.03593778 |
| ENSG00000100644 | HIF1A | -2.3585647 | 0.00643515 |
| ENSG00000204977 | TRIM13 | -2.3487972 | 0.02488781 |
| ENSG00000100393 | EP300 | -2.3299326 | 0.002059 |
| ENSG00000163874 | ZC3H12A | -2.3204056 | 0.01651978 |
| ENSG00000189376 | C8orf76 | -2.3191063 | 0.02882527 |
| ENSG00000102908 | NFAT5 | -2.31265 | 0.04949211 |
| ENSG00000059728 | MXD1 | -2.3005657 | 0.04005277 |
| ENSG00000122644 | ARL4A | -2.3000878 | 0.0498361 |
| ENSG00000115946 | PNO1 | -2.2847106 | 0.01651978 |
| ENSG00000270681 | AC095055.1 | -2.2813465 | 0.01046554 |
| ENSG00000145241 | CENPC | -2.2796577 | 0.03259356 |
| ENSG00000120733 | KDM3B | -2.2779398 | 0.02350696 |
| ENSG00000005483 | KMT2E | -2.2773755 | 0.00952539 |
| ENSG00000178381 | ZFAND2A | -2.2759391 | 0.01529195 |

| | | | |
|---|---|---|---|
| ENSG00000255198 | SNHG9 | -2.2736608 | 0.01641025 |
| ENSG00000163605 | PPP4R2 | -2.264769 | 0.03228479 |
| ENSG00000001561 | ENPP4 | -2.2537182 | 0.04703942 |
| ENSG00000117569 | PTBP2 | -2.2519153 | 0.03034903 |
| ENSG00000080822 | CLDND1 | -2.2506605 | 0.03814153 |
| ENSG00000188647 | PTAR1 | -2.2470642 | 0.02053825 |
| ENSG00000114098 | ARMC8 | -2.2421448 | 0.02589679 |
| ENSG00000112137 | PHACTR1 | -2.2401225 | 0.00670978 |
| ENSG00000115520 | COQ10B | -2.2373364 | 0.013493 |
| ENSG00000091164 | TXNL1 | -2.2326181 | 0.00709631 |
| ENSG00000075415 | SLC25A3 | -2.2167682 | 0.04792734 |
| ENSG00000153922 | CHD1 | -2.2050932 | 0.00373194 |
| ENSG00000102580 | DNAJC3 | -2.2043728 | 0.00427995 |
| ENSG00000047634 | SCML1 | -2.2033829 | 0.04637292 |
| ENSG00000271614 | ATP2B1-AS1 | -2.2008705 | 0.02056281 |
| ENSG00000117139 | KDM5B | -2.1966822 | 0.01686697 |
| ENSG00000100614 | PPM1A | -2.1878349 | 0.04597372 |
| ENSG00000136603 | SKIL | -2.18575 | 0.02360332 |
| ENSG00000155307 | SAMSN1 | -2.1854028 | 0.04158965 |
| ENSG00000105968 | H2AFV | -2.1838924 | 0.01212266 |
| ENSG00000121741 | ZMYM2 | -2.1836391 | 0.01040756 |
| ENSG00000161011 | SQSTM1 | -2.1831974 | 0.02504834 |
| ENSG00000032219 | ARID4A | -2.1799884 | 0.00498634 |
| ENSG00000138593 | SECISBP2L | -2.179669 | 0.02398012 |
| ENSG00000067064 | IDI1 | -2.1776063 | 0.0095478 |
| ENSG00000117410 | ATP6V0B | -2.1766065 | 0.03645327 |
| ENSG00000064012 | CASP8 | -2.164785 | 0.04923352 |
| ENSG00000137947 | GTF2B | -2.1631284 | 0.01651978 |
| ENSG00000126524 | SBDS | -2.1627109 | 0.0088983 |
| ENSG00000087460 | GNAS | -2.1617571 | 0.0383737 |
| ENSG00000106615 | RHEB | -2.1573998 | 0.01404513 |
| ENSG00000008294 | SPAG9 | -2.1568151 | 0.03950432 |
| ENSG00000163788 | SNRK | -2.1497816 | 0.01688977 |
| ENSG00000160570 | DEDD2 | -2.1427921 | 0.03185424 |
| ENSG00000111011 | RSRC2 | -2.1414172 | 0.00366462 |
| ENSG00000180747 | SMG1P3 | -2.1398988 | 0.04580405 |
| ENSG00000197021 | CXorf40B | -2.1371799 | 0.01981956 |
| ENSG00000048405 | ZNF800 | -2.1367196 | 0.01636258 |
| ENSG00000144747 | TMF1 | -2.1363366 | 0.00894515 |
| ENSG00000113580 | NR3C1 | -2.1357775 | 0.02274656 |

| | | | |
|---|---|---|---|
| ENSG00000083799 | CYLD | -2.128274 | 0.03645327 |
| ENSG00000213923 | CSNK1E | -2.1231989 | 0.0088983 |
| ENSG00000176407 | KCMF1 | -2.1192581 | 0.01565564 |
| ENSG00000161526 | SAP30BP | -2.118821 | 0.01633796 |
| ENSG00000118689 | FOXO3 | -2.1162289 | 0.00414186 |
| ENSG00000113811 | SELENOK | -2.1155648 | 0.00366462 |
| ENSG00000139697 | SBNO1 | -2.1025971 | 0.013493 |
| ENSG00000135093 | USP30 | -2.0988346 | 0.02085529 |
| ENSG00000006607 | FARP2 | -2.0975951 | 0.03960245 |
| ENSG00000132823 | OSER1 | -2.0975217 | 0.02360332 |
| ENSG00000165806 | CASP7 | -2.0951807 | 0.02132332 |
| ENSG00000136451 | VEZF1 | -2.0882355 | 0.00450671 |
| ENSG00000187837 | HIST1H1C | -2.0821429 | 0.02440446 |
| ENSG00000176845 | METRNL | -2.080076 | 0.00373194 |
| ENSG00000140743 | CDR2 | -2.0730612 | 0.02259125 |
| ENSG00000162664 | ZNF326 | -2.0716828 | 0.00471655 |
| ENSG00000005339 | CREBBP | -2.0672713 | 0.0360832 |
| ENSG00000124688 | MAD2L1BP | -2.0644079 | 0.00231608 |
| ENSG00000160799 | CCDC12 | -2.0627806 | 0.0184155 |
| ENSG00000020633 | RUNX3 | -2.0618496 | 0.03365163 |
| ENSG00000100528 | CNIH1 | -2.053928 | 0.01118014 |
| ENSG00000138050 | THUMPD2 | -2.0529135 | 0.02922772 |
| ENSG00000140995 | DEF8 | -2.0505927 | 0.03309042 |
| ENSG00000115738 | ID2 | -2.0494412 | 0.01003557 |
| ENSG00000105849 | TWISTNB | -2.0375851 | 0.00366462 |
| ENSG00000015568 | RGPD5 | -2.0328509 | 0.00462374 |
| ENSG00000186162 | CIDECP | -2.0315175 | 0.00471655 |
| ENSG00000124226 | RNF114 | -2.0279585 | 0.00366462 |
| ENSG00000152484 | USP12 | -2.024435 | 0.0278803 |
| ENSG00000113163 | COL4A3BP | -2.0221942 | 0.00366462 |
| ENSG00000122068 | FYTTD1 | -2.018704 | 0.00695581 |
| ENSG00000113615 | SEC24A | -2.0168449 | 0.03950432 |
| ENSG00000239305 | RNF103 | -2.0139912 | 0.01786554 |
| ENSG00000074266 | EED | -2.0126258 | 0.03279199 |
| ENSG00000086666 | ZFAND6 | -2.0125353 | 0.00471655 |
| ENSG00000114416 | FXR1 | -2.0109795 | 0.0164287 |
| ENSG00000183484 | GPR132 | -2.0088306 | 0.01901204 |
| ENSG00000070831 | CDC42 | -2.004092 | 0.00950579 |
| ENSG00000123091 | RNF11 | -1.9982661 | 0.03288142 |
| ENSG00000101146 | RAE1 | -1.997937 | 0.01800346 |

| | | | |
|---|---|---|---|
| ENSG00000220205 | VAMP2 | -1.9935544 | 0.00366094 |
| ENSG00000198780 | FAM169A | -1.9880729 | 0.00643515 |
| ENSG00000124177 | CHD6 | -1.9868299 | 0.03012117 |
| ENSG00000083937 | CHMP2B | -1.9863551 | 0.01284201 |
| ENSG00000145779 | TNFAIP8 | -1.9853279 | 0.02833847 |
| ENSG00000155744 | FAM126B | -1.9852854 | 0.04698791 |
| ENSG00000131263 | RLIM | -1.9850418 | 0.023638 |
| ENSG00000156650 | KAT6B | -1.9847407 | 0.00518293 |
| ENSG00000076554 | TPD52 | -1.983622 | 0.02230772 |
| ENSG00000172845 | SP3 | -1.9789946 | 0.01831296 |
| ENSG00000179833 | SERTAD2 | -1.9784744 | 0.00471655 |
| ENSG00000108510 | MED13 | -1.9775367 | 0.00565028 |
| ENSG00000213096 | ZNF254 | -1.9762201 | 0.01659642 |
| ENSG00000115310 | RTN4 | -1.974329 | 0.02095779 |
| ENSG00000083896 | YTHDC1 | -1.9737135 | 0.02845741 |
| ENSG00000152133 | GPATCH11 | -1.9736697 | 0.04700928 |
| ENSG00000104447 | TRPS1 | -1.9712435 | 0.0080564 |
| ENSG00000144802 | NFKBIZ | -1.9680102 | 0.03365163 |
| ENSG00000101247 | NDUFAF5 | -1.9651746 | 0.02471963 |
| ENSG00000180228 | PRKRA | -1.9650885 | 0.01530022 |
| ENSG00000278311 | GGNBP2 | -1.9647999 | 0.02052985 |
| ENSG00000183604 | SMG1P5 | -1.9634269 | 0.00910301 |
| ENSG00000101782 | RIOK3 | -1.9623211 | 0.04752992 |
| ENSG00000197857 | ZNF44 | -1.9585977 | 0.01118101 |
| ENSG00000008083 | JARID2 | -1.9565897 | 0.00393434 |
| ENSG00000144566 | RAB5A | -1.9552067 | 0.0157351 |
| ENSG00000111832 | RWDD1 | -1.9549769 | 0.02573744 |
| ENSG00000125651 | GTF2F1 | -1.9502405 | 0.00525643 |
| ENSG00000112242 | E2F3 | -1.9492172 | 0.02046898 |
| ENSG00000057657 | PRDM1 | -1.9490263 | 0.0095478 |
| ENSG00000101596 | SMCHD1 | -1.9456298 | 0.03221962 |
| ENSG00000135018 | UBQLN1 | -1.9455278 | 0.00450671 |
| ENSG00000197019 | SERTAD1 | -1.944542 | 0.00373194 |
| ENSG00000164615 | CAMLG | -1.9443022 | 0.01565564 |
| ENSG00000145675 | PIK3R1 | -1.943078 | 0.04297077 |
| ENSG00000137876 | RSL24D1 | -1.9416112 | 0.01834035 |
| ENSG00000123358 | NR4A1 | -1.9402544 | 0.0291451 |
| ENSG00000124201 | ZNFX1 | -1.9402019 | 0.04010247 |
| ENSG00000119801 | YPEL5 | -1.9398959 | 0.02259125 |
| ENSG00000162923 | WDR26 | -1.938859 | 0.01323649 |

| | | | |
|---|---|---|---|
| ENSG00000168264 | IRF2BP2 | -1.9382528 | 0.00414186 |
| ENSG00000159346 | ADIPOR1 | -1.9348955 | 0.02589679 |
| ENSG00000100425 | BRD1 | -1.9339988 | 0.00943543 |
| ENSG00000101654 | RNMT | -1.9267108 | 0.0376018 |
| ENSG00000103121 | CMC2 | -1.9191775 | 0.01882969 |
| ENSG00000156675 | RAB11FIP1 | -1.9183849 | 0.01401736 |
| ENSG00000006576 | PHTF2 | -1.917113 | 0.01969645 |
| ENSG00000147872 | PLIN2 | -1.91664 | 0.03653071 |
| ENSG00000100852 | ARHGAP5 | -1.9165827 | 0.01922859 |
| ENSG00000170385 | SLC30A1 | -1.9081168 | 0.0498361 |
| ENSG00000140153 | WDR20 | -1.9078328 | 0.00643515 |
| ENSG00000196850 | PPTC7 | -1.9043128 | 0.02750331 |
| ENSG00000096746 | HNRNPH3 | -1.8950537 | 0.002059 |
| ENSG00000272196 | HIST2H2AA4 | -1.8877796 | 0.01651978 |
| ENSG00000129351 | ILF3 | -1.8846709 | 0.01824229 |
| ENSG00000120727 | PAIP2 | -1.8772952 | 0.0467091 |
| ENSG00000095574 | IKZF5 | -1.8754389 | 0.04020907 |
| ENSG00000082515 | MRPL22 | -1.8707563 | 0.03228368 |
| ENSG00000150991 | UBC | -1.8689767 | 0.02820495 |
| ENSG00000187522 | HSPA14 | -1.867056 | 0.00414186 |
| ENSG00000183624 | HMCES | -1.863544 | 0.00231608 |
| ENSG00000027697 | IFNGR1 | -1.8604347 | 0.00518293 |
| ENSG00000134453 | RBM17 | -1.8584692 | 0.02610459 |
| ENSG00000173276 | ZBTB21 | -1.8564865 | 0.03259356 |
| ENSG00000101544 | ADNP2 | -1.8551142 | 0.0322021 |
| ENSG00000132475 | H3F3B | -1.8493775 | 0.0108102 |
| ENSG00000169826 | CSGALNACT2 | -1.8489192 | 0.02540537 |
| ENSG00000184182 | UBE2F | -1.8486154 | 0.03300659 |
| ENSG00000146425 | DYNLT1 | -1.8468949 | 0.04820922 |
| ENSG00000230551 | AC021078.1 | -1.8467533 | 0.00674003 |
| ENSG00000269926 | DDIT4-AS1 | -1.8458646 | 0.04840339 |
| ENSG00000094975 | SUCO | -1.8424783 | 0.03515012 |
| ENSG00000265681 | RPL17 | -1.8407111 | 0.01750196 |
| ENSG00000101558 | VAPA | -1.84037 | 0.04158427 |
| ENSG00000004897 | CDC27 | -1.8398906 | 0.03650675 |
| ENSG00000097007 | ABL1 | -1.8381003 | 0.0154968 |
| ENSG00000106829 | TLE4 | -1.8359712 | 0.02841211 |
| ENSG00000182149 | IST1 | -1.8342697 | 0.00718932 |
| ENSG00000228830 | AL160408.2 | -1.8299424 | 0.00366462 |
| ENSG00000101109 | STK4 | -1.8210315 | 0.00414186 |

| | | | |
|---|---|---|---|
| ENSG00000138750 | NUP54 | -1.8166616 | 0.00518293 |
| ENSG00000109466 | KLHL2 | -1.8165383 | 0.01488832 |
| ENSG00000112773 | FAM46A | -1.8155196 | 0.03645327 |
| ENSG00000075426 | FOSL2 | -1.8140914 | 0.0416784 |
| ENSG00000019995 | ZRANB1 | -1.812823 | 0.03764973 |
| ENSG00000176624 | MEX3C | -1.8101519 | 0.002059 |
| ENSG00000177885 | GRB2 | -1.8096175 | 0.00678127 |
| ENSG00000115540 | MOB4 | -1.8093547 | 0.00414186 |
| ENSG00000196396 | PTPN1 | -1.8083053 | 0.0075056 |
| ENSG00000163960 | UBXN7 | -1.8042536 | 0.03602727 |
| ENSG00000140455 | USP3 | -1.8015012 | 0.04024955 |
| ENSG00000196504 | PRPF40A | -1.8012192 | 0.01734575 |
| ENSG00000100483 | VCPKMT | -1.8006405 | 0.01298825 |
| ENSG00000008952 | SEC62 | -1.7986837 | 0.02615942 |
| ENSG00000183735 | TBK1 | -1.7949466 | 0.03259356 |
| ENSG00000124762 | CDKN1A | -1.790485 | 0.03653071 |
| ENSG00000068745 | IP6K2 | -1.7896868 | 0.0064376 |
| ENSG00000143702 | CEP170 | -1.7801982 | 0.01863023 |
| ENSG00000011007 | ELOA | -1.77355 | 0.01599822 |
| ENSG00000184007 | PTP4A2 | -1.7733847 | 0.01801069 |
| ENSG00000137815 | RTF1 | -1.7721063 | 0.02407448 |
| ENSG00000108175 | ZMIZ1 | -1.7699173 | 0.01301122 |
| ENSG00000134058 | CDK7 | -1.7674828 | 0.01404513 |
| ENSG00000033327 | GAB2 | -1.7656548 | 0.00427995 |
| ENSG00000156535 | CD109 | -1.7612595 | 0.01948002 |
| ENSG00000118181 | RPS25 | -1.7593344 | 0.04113773 |
| ENSG00000116030 | SUMO1 | -1.7579458 | 0.00518293 |
| ENSG00000104388 | RAB2A | -1.7575095 | 0.01753258 |
| ENSG00000260708 | AL118516.1 | -1.7563172 | 0.01641025 |
| ENSG00000122482 | ZNF644 | -1.752177 | 0.01423498 |
| ENSG00000185811 | IKZF1 | -1.7518942 | 0.00538725 |
| ENSG00000198265 | HELZ | -1.7505386 | 0.02861146 |
| ENSG00000198355 | PIM3 | -1.748421 | 0.01386124 |
| ENSG00000168092 | PAFAH1B2 | -1.7443667 | 0.01740339 |
| ENSG00000112096 | SOD2 | -1.7439983 | 0.00366462 |
| ENSG00000197780 | TAF13 | -1.7436453 | 0.03327495 |
| ENSG00000142227 | EMP3 | -1.7416279 | 0.02066978 |
| ENSG00000023287 | RB1CC1 | -1.7411566 | 0.03095398 |
| ENSG00000168214 | RBPJ | -1.7361356 | 0.02765943 |
| ENSG00000162910 | MRPL55 | -1.7361119 | 0.04949211 |

| | | | |
|---|---|---|---|
| ENSG00000168438 | CDC40 | -1.7360775 | 0.01602721 |
| ENSG00000106052 | TAX1BP1 | -1.7324488 | 0.00349365 |
| ENSG00000047249 | ATP6V1H | -1.7225702 | 0.00694626 |
| ENSG00000121749 | TBC1D15 | -1.7182926 | 0.01118014 |
| ENSG00000132478 | UNK | -1.7155076 | 0.04727882 |
| ENSG00000147548 | NSD3 | -1.7152432 | 0.01757603 |
| ENSG00000185947 | ZNF267 | -1.712521 | 0.04169608 |
| ENSG00000131876 | SNRPA1 | -1.7094338 | 0.01750196 |
| ENSG00000105993 | DNAJB6 | -1.7081557 | 0.00562571 |
| ENSG00000051108 | HERPUD1 | -1.7078525 | 0.02402911 |
| ENSG00000118620 | ZNF430 | -1.7032836 | 0.01359262 |
| ENSG00000182568 | SATB1 | -1.701464 | 0.01892978 |
| ENSG00000099622 | CIRBP | -1.7006455 | 0.01599822 |
| ENSG00000275740 | AC091959.3 | -1.7000485 | 0.02098066 |
| ENSG00000117614 | SYF2 | -1.6995587 | 0.01404513 |
| ENSG00000143761 | ARF1 | -1.6975774 | 0.03298113 |
| ENSG00000109920 | FNBP4 | -1.6964028 | 0.01659642 |
| ENSG00000153066 | TXNDC11 | -1.6916219 | 0.02356641 |
| ENSG00000089737 | DDX24 | -1.6900113 | 0.00552569 |
| ENSG00000164611 | PTTG1 | -1.6888559 | 0.01921875 |
| ENSG00000150787 | PTS | -1.6860297 | 0.00684755 |
| ENSG00000128016 | ZFP36 | -1.6833588 | 0.01528261 |
| ENSG00000255073 | ZFP91-CNTF | -1.682648 | 0.02815656 |
| ENSG00000198815 | FOXJ3 | -1.6814044 | 0.00471655 |
| ENSG00000187109 | NAP1L1 | -1.6785751 | 0.02552889 |
| ENSG00000116670 | MAD2L2 | -1.6785118 | 0.02775042 |
| ENSG00000116560 | SFPQ | -1.6754683 | 0.01636932 |
| ENSG00000173960 | UBXN2A | -1.6747454 | 0.00972062 |
| ENSG00000100083 | GGA1 | -1.6745857 | 0.01040756 |
| ENSG00000166848 | TERF2IP | -1.6732968 | 0.03645327 |
| ENSG00000188342 | GTF2F2 | -1.6714031 | 0.00427995 |
| ENSG00000132326 | PER2 | -1.6705062 | 0.01473428 |
| ENSG00000196470 | SIAH1 | -1.6700226 | 0.03185424 |
| ENSG00000089234 | BRAP | -1.6662239 | 0.02841211 |
| ENSG00000034677 | RNF19A | -1.6651371 | 0.04313599 |
| ENSG00000167193 | CRK | -1.6582165 | 0.03309042 |
| ENSG00000132819 | RBM38 | -1.6578812 | 0.02822721 |
| ENSG00000124214 | STAU1 | -1.6562073 | 0.03815966 |
| ENSG00000162928 | PEX13 | -1.6557464 | 0.01698373 |
| ENSG00000070495 | JMJD6 | -1.6520593 | 0.01492402 |

| | | | |
|---|---|---|---|
| ENSG00000163682 | RPL9 | -1.6475759 | 0.02776518 |
| ENSG00000234545 | FAM133B | -1.6421073 | 0.01530022 |
| ENSG00000136819 | C9orf78 | -1.6385135 | 0.0060948 |
| ENSG00000217128 | FNIP1 | -1.6363172 | 0.02409658 |
| ENSG00000106723 | SPIN1 | -1.6352289 | 0.04870387 |
| ENSG00000143774 | GUK1 | -1.6344216 | 0.04337462 |
| ENSG00000169641 | LUZP1 | -1.6338975 | 0.03185424 |
| ENSG00000269968 | AC006064.4 | -1.6316226 | 0.01882969 |
| ENSG00000122406 | RPL5 | -1.6300295 | 0.02259125 |
| ENSG00000133226 | SRRM1 | -1.6299634 | 0.01882969 |
| ENSG00000114125 | RNF7 | -1.6273242 | 0.03343504 |
| ENSG00000151332 | MBIP | -1.6225855 | 0.01681343 |
| ENSG00000155508 | CNOT8 | -1.6224458 | 0.03766087 |
| ENSG00000163374 | YY1AP1 | -1.6221834 | 0.02562787 |
| ENSG00000138032 | PPM1B | -1.613043 | 0.02578823 |
| ENSG00000139572 | GPR84 | -1.6119569 | 0.04491806 |
| ENSG00000091527 | CDV3 | -1.6111565 | 0.00715816 |
| ENSG00000175348 | TMEM9B | -1.6109897 | 0.01067079 |
| ENSG00000106245 | BUD31 | -1.6072295 | 0.00366462 |
| ENSG00000132334 | PTPRE | -1.6041146 | 0.01101924 |
| ENSG00000234745 | HLA-B | -1.6021945 | 0.02302363 |
| ENSG00000006634 | DBF4 | -1.6018116 | 0.0287894 |
| ENSG00000179295 | PTPN11 | -1.6006623 | 0.01017501 |
| ENSG00000156030 | ELMSAN1 | -1.5993026 | 0.00366462 |
| ENSG00000061936 | SFSWAP | -1.5980879 | 0.03455066 |
| ENSG00000132388 | UBE2G1 | -1.5863168 | 0.00427995 |
| ENSG00000100982 | PCIF1 | -1.586076 | 0.00458777 |
| ENSG00000164032 | H2AFZ | -1.5829662 | 0.02358364 |
| ENSG00000117523 | PRRC2C | -1.5823651 | 0.00421696 |
| ENSG00000168615 | ADAM9 | -1.5815231 | 0.04551538 |
| ENSG00000168137 | SETD5 | -1.5803551 | 0.04087958 |
| ENSG00000116752 | BCAS2 | -1.5794338 | 0.04794577 |
| ENSG00000033030 | ZCCHC8 | -1.5790862 | 0.04297077 |
| ENSG00000163624 | CDS1 | -1.5761092 | 0.04870387 |
| ENSG00000119048 | UBE2B | -1.5728153 | 0.03917306 |
| ENSG00000166747 | AP1G1 | -1.5719046 | 0.0383737 |
| ENSG00000255112 | CHMP1B | -1.5712458 | 0.04448535 |
| ENSG00000143079 | CTTNBP2NL | -1.5684593 | 0.01403954 |
| ENSG00000146232 | NFKBIE | -1.5668712 | 0.02052812 |
| ENSG00000151461 | UPF2 | -1.5632883 | 0.0037268 |

| | | | |
|---|---|---|---|
| ENSG00000180596 | HIST1H2BC | -1.5602294 | 0.023638 |
| ENSG00000054267 | ARID4B | -1.5600476 | 0.0080564 |
| ENSG00000179094 | PER1 | -1.5596991 | 0.03414375 |
| ENSG00000147604 | RPL7 | -1.5523558 | 0.03067089 |
| ENSG00000056097 | ZFR | -1.5443294 | 0.02861146 |
| ENSG00000090061 | CCNK | -1.5435958 | 0.0108102 |
| ENSG00000185043 | CIB1 | -1.5420194 | 0.00526983 |
| ENSG00000177879 | AP3S1 | -1.5417923 | 0.00517546 |
| ENSG00000170889 | RPS9 | -1.5371501 | 0.04672437 |
| ENSG00000145414 | NAF1 | -1.5358676 | 0.00939014 |
| ENSG00000134248 | LAMTOR5 | -1.535767 | 0.01905624 |
| ENSG00000267165 | CHMP1B-AS1 | -1.53324 | 0.03839989 |
| ENSG00000205937 | RNPS1 | -1.531859 | 0.01764812 |
| ENSG00000137955 | RABGGTB | -1.5311364 | 0.02187734 |
| ENSG00000147526 | TACC1 | -1.5295004 | 0.00817443 |
| ENSG00000271869 | AC026979.3 | -1.5276153 | 0.01698373 |
| ENSG00000144848 | ATG3 | -1.5257477 | 0.02328206 |
| ENSG00000198160 | MIER1 | -1.5249849 | 0.00783832 |
| ENSG00000166266 | CUL5 | -1.5246382 | 0.02315457 |
| ENSG00000138326 | RPS24 | -1.5235435 | 0.0495915 |
| ENSG00000101166 | PRELID3B | -1.5209207 | 0.01745368 |
| ENSG00000187514 | PTMA | -1.5196051 | 0.00812086 |
| ENSG00000102409 | BEX4 | -1.5147338 | 0.00366462 |
| ENSG00000204178 | TMEM57 | -1.5111317 | 0.04977237 |
| ENSG00000184014 | DENND5A | -1.5096529 | 0.0311748 |
| ENSG00000109220 | CHIC2 | -1.5092584 | 0.02274656 |
| ENSG00000143256 | PFDN2 | -1.5092422 | 0.02230772 |
| ENSG00000168036 | CTNNB1 | -1.5081516 | 0.0488667 |
| ENSG00000259884 | AC025259.3 | -1.5079142 | 0.0291451 |
| ENSG00000156735 | BAG4 | -1.507731 | 0.03297477 |
| ENSG00000119844 | AFTPH | -1.5075552 | 0.0383737 |
| ENSG00000137185 | ZSCAN9 | -1.5056565 | 0.04795835 |
| ENSG00000184922 | FMNL1 | -1.5042614 | 0.00427995 |
| ENSG00000203644 | AC083799.1 | -1.5031031 | 0.00604041 |
| ENSG00000129484 | PARP2 | -1.5019141 | 0.01498374 |
| ENSG00000172062 | SMN1 | -1.5017795 | 0.00809412 |
| ENSG00000114354 | TFG | -1.5006429 | 0.02308955 |
| ENSG00000090905 | TNRC6A | -1.5001209 | 0.01786554 |

#### DEG Mo pSS versus HD

| Transcipt ID | Gene name | Log2(FC) | FDR < 0.05 |
|---|---|---|---|
| ENSG00000088827 | SIGLEC1 | 4.81222715 | 0.00061969 |
| ENSG00000127951 | FGL2 | 4.63307294 | 3.69E-05 |
| ENSG00000177409 | SAMD9L | 4.48867916 | 7.2716E-05 |
| ENSG00000111331 | OAS3 | 4.42230211 | 0.00037088 |
| ENSG00000111335 | OAS2 | 4.41241143 | 0.00023971 |
| ENSG00000182578 | CSF1R | 4.12149732 | 0.00018821 |
| ENSG00000111913 | RIPOR2 | 4.05133531 | 5.6748E-05 |
| ENSG00000204131 | NHSL2 | 3.98423953 | 8.698E-05 |
| ENSG00000089127 | OAS1 | 3.88177349 | 0.00014848 |
| ENSG00000082074 | FYB1 | 3.86940796 | 0.00011131 |
| ENSG00000168329 | CX3CR1 | 3.66901016 | 0.00346942 |
| ENSG00000132530 | XAF1 | 3.57786021 | 0.00019855 |
| ENSG00000137959 | IFI44L | 3.56620595 | 0.00405956 |
| ENSG00000106780 | MEGF9 | 3.50812058 | 0.00020506 |
| ENSG00000110876 | SELPLG | 3.45149629 | 3.725E-05 |
| ENSG00000119917 | IFIT3 | 3.41478814 | 0.00578098 |
| ENSG00000157601 | MX1 | 3.40051191 | 0.00273973 |
| ENSG00000093072 | ADA2 | 3.30929409 | 0.00035785 |
| ENSG00000137628 | DDX60 | 3.26174787 | 0.00017728 |
| ENSG00000119922 | IFIT2 | 3.15400711 | 0.00698448 |
| ENSG00000096968 | JAK2 | 3.1171167 | 4.002E-05 |
| ENSG00000183486 | MX2 | 3.0908071 | 0.00012058 |
| ENSG00000203747 | FCGR3A | 3.05824159 | 0.00908083 |
| ENSG00000185745 | IFIT1 | 3.04472512 | 0.00769807 |
| ENSG00000092964 | DPYSL2 | 3.03157003 | 0.00036453 |
| ENSG00000179583 | CIITA | 2.97329048 | 3.69E-05 |
| ENSG00000178927 | C17orf62 | 2.95098495 | 0.00011008 |
| ENSG00000146192 | FGD2 | 2.94266455 | 3.69E-05 |
| ENSG00000171115 | GIMAP8 | 2.91127768 | 0.00035769 |
| ENSG00000121858 | TNFSF10 | 2.90815474 | 0.00085748 |
| ENSG00000101347 | SAMHD1 | 2.8875798 | 0.00020568 |
| ENSG00000107551 | RASSF4 | 2.86741339 | 0.00020286 |
| ENSG00000133706 | LARS | 2.85358246 | 0.00024204 |
| ENSG00000157637 | SLC38A10 | 2.83559968 | 4.0167E-05 |
| ENSG00000105483 | CARD8 | 2.82180872 | 0.00010905 |
| ENSG00000134321 | RSAD2 | 2.8216868 | 0.00044978 |
| ENSG00000166801 | FAM111A | 2.82114926 | 4.0167E-05 |

| | | | |
|---|---|---|---|
| ENSG00000133574 | GIMAP4 | 2.80831292 | 0.00205538 |
| ENSG00000005020 | SKAP2 | 2.80342257 | 0.00011131 |
| ENSG00000107201 | DDX58 | 2.79847046 | 7.6564E-05 |
| ENSG00000139970 | RTN1 | 2.79625262 | 0.00161234 |
| ENSG00000143624 | INTS3 | 2.76935299 | 0.00025431 |
| ENSG00000135218 | CD36 | 2.7687755 | 0.00441468 |
| ENSG00000121807 | CCR2 | 2.76812231 | 0.00459916 |
| ENSG00000134326 | CMPK2 | 2.76441331 | 7.007E-05 |
| ENSG00000124491 | F13A1 | 2.76137384 | 0.0436662 |
| ENSG00000038427 | VCAN | 2.76089778 | 0.00031344 |
| ENSG00000139278 | GLIPR1 | 2.75148728 | 3.69E-05 |
| ENSG00000188641 | DPYD | 2.7460871 | 0.00016589 |
| ENSG00000160593 | JAML | 2.73785141 | 1.5464E-05 |
| ENSG00000129003 | VPS13C | 2.72450583 | 0.00042793 |
| ENSG00000177575 | CD163 | 2.7119008 | 0.00130504 |
| ENSG00000101336 | HCK | 2.69647494 | 4.6204E-05 |
| ENSG00000196975 | ANXA4 | 2.68121227 | 0.00013137 |
| ENSG00000143546 | S100A8 | 2.67257265 | 0.00020529 |
| ENSG00000133835 | HSD17B4 | 2.66258299 | 5.2951E-05 |
| ENSG00000103381 | CPPED1 | 2.65717733 | 0.00031447 |
| ENSG00000181381 | DDX60L | 2.6567232 | 0.00050571 |
| ENSG00000105953 | OGDH | 2.64511854 | 0.00021448 |
| ENSG00000130449 | ZSWIM6 | 2.63544798 | 7.2413E-05 |
| ENSG00000187608 | ISG15 | 2.62966453 | 0.00010145 |
| ENSG00000165168 | CYBB | 2.62649243 | 0.00017515 |
| ENSG00000163683 | SMIM14 | 2.584365 | 0.00013629 |
| ENSG00000215458 | AATBC | 2.57203412 | 5.6247E-05 |
| ENSG00000196209 | SIRPB2 | 2.55920596 | 0.00116963 |
| ENSG00000073849 | ST6GAL1 | 2.550513 | 0.00048075 |
| ENSG00000249437 | NAIP | 2.54772158 | 0.03856131 |
| ENSG00000138119 | MYOF | 2.52298352 | 0.00116629 |
| ENSG00000133106 | EPSTI1 | 2.51730507 | 0.00024243 |
| ENSG00000163563 | MNDA | 2.51154611 | 0.00019527 |
| ENSG00000240065 | PSMB9 | 2.49985703 | 0.00043175 |
| ENSG00000134256 | CD101 | 2.49461745 | 0.00672014 |
| ENSG00000185722 | ANKFY1 | 2.4915029 | 0.00017719 |
| ENSG00000006756 | ARSD | 2.456905 | 0.00034289 |
| ENSG00000068079 | IFI35 | 2.45084083 | 0.00011131 |
| ENSG00000102524 | TNFSF13B | 2.4295903 | 0.00016701 |
| ENSG00000110665 | C11orf21 | 2.42600156 | 4.3201E-05 |
| ENSG00000142405 | NLRP12 | 2.42233631 | 0.0011604 |
| ENSG00000213203 | GIMAP1 | 2.40060946 | 0.00029261 |

| | | | |
|---|---|---|---|
| ENSG00000250138 | AC139495.3 | 2.39948877 | 0.00070469 |
| ENSG00000127954 | STEAP4 | 2.39860441 | 0.02252014 |
| ENSG00000138246 | DNAJC13 | 2.38403223 | 0.00029419 |
| ENSG00000178685 | PARP10 | 2.38363732 | 1.8312E-05 |
| ENSG00000128872 | TMOD2 | 2.38333014 | 0.00011131 |
| ENSG00000163565 | IFI16 | 2.37871911 | 5.9604E-05 |
| ENSG00000165092 | ALDH1A1 | 2.36838002 | 0.00200702 |
| ENSG00000149131 | SERPING1 | 2.36410268 | 0.0017831 |
| ENSG00000100364 | KIAA0930 | 2.3531638 | 7.9831E-05 |
| ENSG00000135636 | DYSF | 2.35302925 | 0.00056476 |
| ENSG00000126709 | IFI6 | 2.35060464 | 0.00045242 |
| ENSG00000111961 | SASH1 | 2.34814151 | 0.04833761 |
| ENSG00000050344 | NFE2L3 | 2.33003379 | 5.9604E-05 |
| ENSG00000142089 | IFITM3 | 2.32819656 | 0.00076829 |
| ENSG00000146592 | CREB5 | 2.32706132 | 0.00069182 |
| ENSG00000119638 | NEK9 | 2.32309385 | 8.2108E-05 |
| ENSG00000140575 | IQGAP1 | 2.30892835 | 0.00026643 |
| ENSG00000105967 | TFEC | 2.30676328 | 8.2108E-05 |
| ENSG00000081189 | MEF2C | 2.29629681 | 0.00037032 |
| ENSG00000142687 | KIAA0319L | 2.28793993 | 0.0002003 |
| ENSG00000137965 | IFI44 | 2.27380106 | 0.00211031 |
| ENSG00000138413 | IDH1 | 2.27237293 | 0.00056064 |
| ENSG00000168310 | IRF2 | 2.27032454 | 0.00101781 |
| ENSG00000229754 | CXCR2P1 | 2.26268149 | 0.0008627 |
| ENSG00000106785 | TRIM14 | 2.26268053 | 0.00011864 |
| ENSG00000135709 | KIAA0513 | 2.26070245 | 0.00011131 |
| ENSG00000140749 | IGSF6 | 2.25914668 | 0.00110115 |
| ENSG00000188554 | NBR1 | 2.24022924 | 0.00011131 |
| ENSG00000058668 | ATP2B4 | 2.23782734 | 0.00196697 |
| ENSG00000165071 | TMEM71 | 2.22947868 | 0.00013137 |
| ENSG00000175567 | UCP2 | 2.2209786 | 0.00179198 |
| ENSG00000031081 | ARHGAP31 | 2.21924073 | 0.00241362 |
| ENSG00000133943 | DGLUCY | 2.21857708 | 4.0167E-05 |
| ENSG00000187554 | TLR5 | 2.21767842 | 0.00029084 |
| ENSG00000188404 | SELL | 2.21406286 | 0.00152087 |
| ENSG00000102189 | EEA1 | 2.21309813 | 0.00046523 |
| ENSG00000180357 | ZNF609 | 2.21122874 | 6.6556E-05 |
| ENSG00000138646 | HERC5 | 2.21122579 | 0.00014889 |
| ENSG00000103313 | MEFV | 2.20838651 | 0.00024218 |
| ENSG00000139687 | RB1 | 2.20778929 | 9.703E-05 |
| ENSG00000115232 | ITGA4 | 2.1977504 | 0.00066046 |
| ENSG00000081087 | OSTM1 | 2.18697561 | 5.2951E-05 |

| | | | |
|---|---|---|---|
| ENSG00000115155 | OTOF | 2.17916828 | 0.04578118 |
| ENSG00000110077 | MS4A6A | 2.17144347 | 0.00035769 |
| ENSG00000173821 | RNF213 | 2.16238113 | 0.00043225 |
| ENSG00000198734 | F5 | 2.15850947 | 0.00025416 |
| ENSG00000198951 | NAGA | 2.1583494 | 0.00023076 |
| ENSG00000150867 | PIP4K2A | 2.15794943 | 0.00017634 |
| ENSG00000071967 | CYBRD1 | 2.15624005 | 0.00158569 |
| ENSG00000074706 | IPCEF1 | 2.15391304 | 0.00079352 |
| ENSG00000166326 | TRIM44 | 2.15103466 | 0.00011131 |
| ENSG00000121210 | TMEM131L | 2.15055037 | 0.00080531 |
| ENSG00000108771 | DHX58 | 2.12712832 | 0.00088319 |
| ENSG00000112367 | FIG4 | 2.11750045 | 2.6941E-05 |
| ENSG00000165672 | PRDX3 | 2.11413223 | 0.00011131 |
| ENSG00000138756 | BMP2K | 2.10876117 | 0.00014473 |
| ENSG00000183023 | SLC8A1 | 2.10770505 | 0.00140107 |
| ENSG00000161929 | SCIMP | 2.10518218 | 0.00054607 |
| ENSG00000082996 | RNF13 | 2.1003053 | 0.00042489 |
| ENSG00000107099 | DOCK8 | 2.09599002 | 0.00042323 |
| ENSG00000025708 | TYMP | 2.09573646 | 0.00353048 |
| ENSG00000160551 | TAOK1 | 2.0948519 | 0.00031447 |
| ENSG00000163946 | FAM208A | 2.08670497 | 0.00025562 |
| ENSG00000179144 | GIMAP7 | 2.08182365 | 0.01499835 |
| ENSG00000138496 | PARP9 | 2.07771882 | 0.00058507 |
| ENSG00000135838 | NPL | 2.07449096 | 0.00096657 |
| ENSG00000173083 | HPSE | 2.06771702 | 0.00112582 |
| ENSG00000119321 | FKBP15 | 2.06242754 | 0.00027781 |
| ENSG00000166927 | MS4A7 | 2.06134147 | 0.00363434 |
| ENSG00000143970 | ASXL2 | 2.05713821 | 0.0002004 |
| ENSG00000124942 | AHNAK | 2.05330381 | 0.00037122 |
| ENSG00000145246 | ATP10D | 2.04835108 | 0.00039682 |
| ENSG00000149311 | ATM | 2.04810493 | 3.69E-05 |
| ENSG00000165476 | REEP3 | 2.04794403 | 0.0002967 |
| ENSG00000133561 | GIMAP6 | 2.04177072 | 0.00071202 |
| ENSG00000163220 | S100A9 | 2.03585241 | 0.00044987 |
| ENSG00000084733 | RAB10 | 2.03363615 | 0.0001934 |
| ENSG00000197043 | ANXA6 | 2.02891787 | 0.0074927 |
| ENSG00000100342 | APOL1 | 2.02806304 | 0.00710375 |
| ENSG00000088986 | DYNLL1 | 2.02678083 | 3.69E-05 |
| ENSG00000180370 | PAK2 | 2.02592389 | 0.00021444 |
| ENSG00000073969 | NSF | 2.02382863 | 0.00021098 |
| ENSG00000133313 | CNDP2 | 2.02318759 | 0.00070469 |
| ENSG00000129566 | TEP1 | 2.02316324 | 0.00094714 |

| | | | |
|---|---|---|---|
| ENSG00000170006 | TMEM154 | 2.02228185 | 0.00033278 |
| ENSG00000128815 | WDFY4 | 2.01621146 | 0.0007939 |
| ENSG00000280153 | AC133065.6 | 2.01591236 | 0.00098903 |
| ENSG00000130589 | HELZ2 | 2.01531518 | 0.00059211 |
| ENSG00000203710 | CR1 | 2.00968798 | 0.00167253 |
| ENSG00000111269 | CREBL2 | 1.99637291 | 0.0004298 |
| ENSG00000160216 | AGPAT3 | 1.9931901 | 0.00080171 |
| ENSG00000188906 | LRRK2 | 1.99158214 | 0.00021321 |
| ENSG00000168010 | ATG16L2 | 1.99121182 | 0.00011877 |
| ENSG00000040933 | INPP4A | 1.99048334 | 8.82E-05 |
| ENSG00000186088 | GSAP | 1.98529176 | 0.00022632 |
| ENSG00000123213 | NLN | 1.98353142 | 0.01069252 |
| ENSG00000122986 | HVCN1 | 1.98183554 | 0.00017899 |
| ENSG00000135457 | TFCP2 | 1.97973391 | 5.9604E-05 |
| ENSG00000136485 | DCAF7 | 1.97954393 | 0.0003913 |
| ENSG00000065413 | ANKRD44 | 1.97584367 | 3.4895E-05 |
| ENSG00000131374 | TBC1D5 | 1.97247733 | 0.00033575 |
| ENSG00000084070 | SMAP2 | 1.97215259 | 0.00518494 |
| ENSG00000155957 | TMBIM4 | 1.97188602 | 4.9812E-05 |
| ENSG00000169220 | RGS14 | 1.9697004 | 5.9265E-05 |
| ENSG00000104133 | SPG11 | 1.96773469 | 8.698E-05 |
| ENSG00000116237 | ICMT | 1.96760299 | 5.4121E-05 |
| ENSG00000145416 | MARCH1 | 1.96445135 | 0.00108021 |
| ENSG00000198585 | NUDT16 | 1.96325971 | 0.00025638 |
| ENSG00000153936 | HS2ST1 | 1.96273796 | 5.4121E-05 |
| ENSG00000137752 | CASP1 | 1.95801983 | 0.00041379 |
| ENSG00000133216 | EPHB2 | 1.94999142 | 0.00037151 |
| ENSG00000158517 | NCF1 | 1.9496836 | 0.00084495 |
| ENSG00000197548 | ATG7 | 1.94397589 | 0.00018547 |
| ENSG00000121281 | ADCY7 | 1.93336445 | 0.00041634 |
| ENSG00000170581 | STAT2 | 1.93316979 | 0.00175064 |
| ENSG00000151466 | SCLT1 | 1.93308209 | 0.00075271 |
| ENSG00000145287 | PLAC8 | 1.93174465 | 0.00184434 |
| ENSG00000166888 | STAT6 | 1.93017847 | 0.00051901 |
| ENSG00000136040 | PLXNC1 | 1.92853493 | 0.00025416 |
| ENSG00000156587 | UBE2L6 | 1.92681789 | 0.00130163 |
| ENSG00000122694 | GLIPR2 | 1.92658528 | 0.00020061 |
| ENSG00000226137 | BAIAP2-AS1 | 1.92619115 | 0.00170844 |
| ENSG00000136631 | VPS45 | 1.92511228 | 0.00010269 |
| ENSG00000166272 | WBP1L | 1.92179463 | 0.00031743 |
| ENSG00000107929 | LARP4B | 1.91918736 | 0.00013762 |
| ENSG00000090863 | GLG1 | 1.90303374 | 3.69E-05 |

| | | | |
|---|---|---|---|
| ENSG00000010810 | FYN | 1.90139286 | 0.00012627 |
| ENSG00000072501 | SMC1A | 1.90091004 | 0.00024409 |
| ENSG00000093144 | ECHDC1 | 1.90087661 | 0.00020061 |
| ENSG00000155660 | PDIA4 | 1.89991655 | 5.8839E-05 |
| ENSG00000150681 | RGS18 | 1.89711599 | 0.00022322 |
| ENSG00000104974 | LILRA1 | 1.89627793 | 0.00225028 |
| ENSG00000198771 | RCSD1 | 1.89415863 | 0.00013674 |
| ENSG00000105501 | SIGLEC5 | 1.89344721 | 0.0002242 |
| ENSG00000103479 | RBL2 | 1.88225062 | 0.00011163 |
| ENSG00000091106 | NLRC4 | 1.88025166 | 0.00037654 |
| ENSG00000103335 | PIEZO1 | 1.87511955 | 0.00136249 |
| ENSG00000010292 | NCAPD2 | 1.869991 | 0.00013089 |
| ENSG00000008130 | NADK | 1.86881805 | 0.000305 |
| ENSG00000162946 | DISC1 | 1.86805719 | 0.00253017 |
| ENSG00000138640 | FAM13A | 1.86643031 | 0.00278912 |
| ENSG00000140090 | SLC24A4 | 1.86540488 | 8.2108E-05 |
| ENSG00000129675 | ARHGEF6 | 1.85487557 | 0.00021321 |
| ENSG00000157827 | FMNL2 | 1.85417167 | 0.00017617 |
| ENSG00000110031 | LPXN | 1.84675535 | 0.00058189 |
| ENSG00000112419 | PHACTR2 | 1.83062034 | 0.0002053 |
| ENSG00000137478 | FCHSD2 | 1.82957906 | 0.00059545 |
| ENSG00000020577 | SAMD4A | 1.82723663 | 6.8112E-05 |
| ENSG00000166197 | NOLC1 | 1.82686298 | 0.00019633 |
| ENSG00000181192 | DHTKD1 | 1.82658412 | 0.00095394 |
| ENSG00000164054 | SHISA5 | 1.82575246 | 0.00014848 |
| ENSG00000171853 | TRAPPC12 | 1.82407323 | 0.00038272 |
| ENSG00000134996 | OSTF1 | 1.82067029 | 0.0001277 |
| ENSG00000132182 | NUP210 | 1.81876085 | 0.0006356 |
| ENSG00000184979 | USP18 | 1.81821607 | 0.00018668 |
| ENSG00000154930 | ACSS1 | 1.81709923 | 0.00043615 |
| ENSG00000167220 | HDHD2 | 1.81576718 | 0.0001277 |
| ENSG00000159339 | PADI4 | 1.8129343 | 0.00127516 |
| ENSG00000151702 | FLI1 | 1.81161047 | 0.00062992 |
| ENSG00000250510 | GPR162 | 1.81124772 | 0.00281025 |
| ENSG00000166928 | MS4A14 | 1.80676496 | 0.00387978 |
| ENSG00000213648 | SULT1A4 | 1.80594935 | 0.00220992 |
| ENSG00000177853 | ZNF518A | 1.80539933 | 0.00031839 |
| ENSG00000145703 | IQGAP2 | 1.80357672 | 0.00900436 |
| ENSG00000120594 | PLXDC2 | 1.79914964 | 0.00207841 |
| ENSG00000121691 | CAT | 1.79908906 | 0.00030322 |
| ENSG00000132256 | TRIM5 | 1.79834867 | 0.00041435 |
| ENSG00000064763 | FAR2 | 1.79614448 | 0.0001848 |

| | | | |
|---|---|---|---|
| ENSG00000197629 | MPEG1 | 1.79410737 | 0.00348206 |
| ENSG00000108389 | MTMR4 | 1.79292546 | 4.0167E-05 |
| ENSG00000258659 | TRIM34 | 1.78714573 | 1.5464E-05 |
| ENSG00000101346 | POFUT1 | 1.78666583 | 0.00019592 |
| ENSG00000263528 | IKBKE | 1.78215843 | 0.00013089 |
| ENSG00000133816 | MICAL2 | 1.77999999 | 0.00965276 |
| ENSG00000105281 | SLC1A5 | 1.77845426 | 0.00096657 |
| ENSG00000138449 | SLC40A1 | 1.77795244 | 0.00044978 |
| ENSG00000185885 | IFITM1 | 1.77549121 | 0.00195944 |
| ENSG00000164808 | SPIDR | 1.77402505 | 4.6204E-05 |
| ENSG00000164414 | SLC35A1 | 1.76140878 | 6.077E-05 |
| ENSG00000214078 | CPNE1 | 1.75638232 | 0.0061442 |
| ENSG00000155097 | ATP6V1C1 | 1.75552976 | 6.8996E-05 |
| ENSG00000130150 | MOSPD2 | 1.75525532 | 0.02616316 |
| ENSG00000164062 | APEH | 1.75516083 | 0.00042376 |
| ENSG00000129993 | CBFA2T3 | 1.75416807 | 0.00034175 |
| ENSG00000196664 | TLR7 | 1.75117674 | 0.03127928 |
| ENSG00000196730 | DAPK1 | 1.74965437 | 0.00305556 |
| ENSG00000145715 | RASA1 | 1.74920879 | 6.4871E-05 |
| ENSG00000250264 | AL669918.1 | 1.74666005 | 0.04869143 |
| ENSG00000001629 | ANKIB1 | 1.7450679 | 4.0167E-05 |
| ENSG00000143669 | LYST | 1.74377161 | 0.0001798 |
| ENSG00000177119 | ANO6 | 1.73797693 | 4.0167E-05 |
| ENSG00000113845 | TIMMDC1 | 1.73688181 | 0.00023416 |
| ENSG00000108798 | ABI3 | 1.73475905 | 0.0010977 |
| ENSG00000076770 | MBNL3 | 1.73193071 | 7.5228E-05 |
| ENSG00000134955 | SLC37A2 | 1.72948671 | 0.00224493 |
| ENSG00000112079 | STK38 | 1.72691606 | 0.00026307 |
| ENSG00000168995 | SIGLEC7 | 1.72316115 | 0.00234137 |
| ENSG00000174600 | CMKLR1 | 1.72163844 | 0.00172775 |
| ENSG00000142784 | WDTC1 | 1.72084308 | 4.0167E-05 |
| ENSG00000118596 | SLC16A7 | 1.71739468 | 0.00224493 |
| ENSG00000159228 | CBR1 | 1.71678557 | 0.00016348 |
| ENSG00000164308 | ERAP2 | 1.71653322 | 0.03650502 |
| ENSG00000117115 | PADI2 | 1.71315931 | 0.00446116 |
| ENSG00000103196 | CRISPLD2 | 1.71032628 | 0.00540432 |
| ENSG00000198814 | GK | 1.708838 | 0.0100024 |
| ENSG00000005844 | ITGAL | 1.70855868 | 0.00248508 |
| ENSG00000136161 | RCBTB2 | 1.70732997 | 0.00050507 |
| ENSG00000165675 | ENOX2 | 1.70371623 | 0.00034874 |
| ENSG00000254838 | GVINP1 | 1.70358337 | 0.00371368 |
| ENSG00000154822 | PLCL2 | 1.70345847 | 0.00242728 |

| | | | |
|---|---|---|---|
| ENSG00000280071 | FP565260.6 | 1.6991183 | 0.0033487 |
| ENSG00000138459 | SLC35A5 | 1.69403765 | 0.00018322 |
| ENSG00000198736 | MSRB1 | 1.68859047 | 0.00042489 |
| ENSG00000114770 | ABCC5 | 1.68856034 | 0.00092437 |
| ENSG00000135842 | FAM129A | 1.68719891 | 0.00173207 |
| ENSG00000178695 | KCTD12 | 1.68715468 | 0.00048075 |
| ENSG00000103544 | C16orf62 | 1.68559026 | 0.00013674 |
| ENSG00000182511 | FES | 1.68541817 | 0.0003573 |
| ENSG00000165819 | METTL3 | 1.67918653 | 0.00013429 |
| ENSG00000143110 | C1orf162 | 1.67827229 | 0.00036405 |
| ENSG00000136816 | TOR1B | 1.67818427 | 0.00021444 |
| ENSG00000184992 | BRI3BP | 1.6780333 | 0.0004298 |
| ENSG00000175538 | KCNE3 | 1.67614598 | 4.3201E-05 |
| ENSG00000101695 | RNF125 | 1.67437375 | 0.00086119 |
| ENSG00000088888 | MAVS | 1.67434021 | 0.00090564 |
| ENSG00000124357 | NAGK | 1.67011388 | 0.00078899 |
| ENSG00000077420 | APBB1IP | 1.66646142 | 0.00011229 |
| ENSG00000006715 | VPS41 | 1.6644439 | 0.00014019 |
| ENSG00000089597 | GANAB | 1.66262829 | 0.00017634 |
| ENSG00000138642 | HERC6 | 1.65949223 | 0.00015833 |
| ENSG00000101307 | SIRPB1 | 1.65722086 | 0.01664438 |
| ENSG00000104365 | IKBKB | 1.65533587 | 0.00016701 |
| ENSG00000175216 | CKAP5 | 1.65500781 | 0.00116057 |
| ENSG00000166002 | SMCO4 | 1.65376135 | 0.00130388 |
| ENSG00000119457 | SLC46A2 | 1.65167071 | 0.0023381 |
| ENSG00000135899 | SP110 | 1.65130811 | 0.00013089 |
| ENSG00000090861 | AARS | 1.64939027 | 0.00110912 |
| ENSG00000026652 | AGPAT4 | 1.64716441 | 0.0008941 |
| ENSG00000204261 | PSMB8-AS1 | 1.64527738 | 0.00138063 |
| ENSG00000103047 | TANGO6 | 1.64454838 | 0.00187327 |
| ENSG00000196782 | MAML3 | 1.64105639 | 0.00074128 |
| ENSG00000142185 | TRPM2 | 1.64064032 | 0.00191016 |
| ENSG00000172164 | SNTB1 | 1.63923717 | 0.0017562 |
| ENSG00000087274 | ADD1 | 1.63577739 | 0.00053834 |
| ENSG00000110934 | BIN2 | 1.63492943 | 0.0018027 |
| ENSG00000075303 | SLC25A40 | 1.63413334 | 0.0001277 |
| ENSG00000175309 | PHYKPL | 1.63396822 | 0.00275355 |
| ENSG00000177628 | GBA | 1.63228568 | 0.00179412 |
| ENSG00000198133 | TMEM229B | 1.6321927 | 0.01145874 |
| ENSG00000175471 | MCTP1 | 1.63149672 | 0.00014889 |
| ENSG00000167085 | PHB | 1.6309904 | 0.00072824 |
| ENSG00000168016 | TRANK1 | 1.63095088 | 0.00080171 |

| | | | |
|---|---|---|---|
| ENSG00000119397 | CNTRL | 1.63015137 | 0.00400993 |
| ENSG00000162704 | ARPC5 | 1.62932633 | 0.00023416 |
| ENSG00000020129 | NCDN | 1.62621196 | 0.00081608 |
| ENSG00000146776 | ATXN7L1 | 1.62399274 | 0.00026334 |
| ENSG00000197142 | ACSL5 | 1.62105862 | 0.00051346 |
| ENSG00000214013 | GANC | 1.62065399 | 0.00014848 |
| ENSG00000115415 | STAT1 | 1.62023511 | 0.00431728 |
| ENSG00000099326 | MZF1 | 1.61914023 | 0.00012317 |
| ENSG00000176783 | RUFY1 | 1.61894132 | 0.00059298 |
| ENSG00000163328 | GPR155 | 1.61543313 | 0.00032578 |
| ENSG00000108468 | CBX1 | 1.61471009 | 0.00010905 |
| ENSG00000205133 | TRIQK | 1.61434355 | 0.00085331 |
| ENSG00000004455 | AK2 | 1.61380087 | 0.00025545 |
| ENSG00000152818 | UTRN | 1.61190641 | 0.00268679 |
| ENSG00000130021 | PUDP | 1.6114027 | 3.69E-05 |
| ENSG00000113522 | RAD50 | 1.60816637 | 0.00048893 |
| ENSG00000148110 | MFSD14B | 1.60660809 | 0.00324543 |
| ENSG00000179921 | GPBAR1 | 1.6060237 | 0.00011774 |
| ENSG00000101337 | TM9SF4 | 1.60519099 | 0.00011131 |
| ENSG00000185482 | STAC3 | 1.60443502 | 0.00015833 |
| ENSG00000130303 | BST2 | 1.60174167 | 0.00075271 |
| ENSG00000112297 | CRYBG1 | 1.59901085 | 0.00026047 |
| ENSG00000086300 | SNX10 | 1.5940843 | 7.5228E-05 |
| ENSG00000179978 | AC140134.1 | 1.59225393 | 0.01034033 |
| ENSG00000134452 | FBXO18 | 1.58838748 | 0.00030322 |
| ENSG00000162869 | PPP1R21 | 1.58588347 | 0.00048075 |
| ENSG00000109436 | TBC1D9 | 1.58441502 | 0.00019192 |
| ENSG00000101974 | ATP11C | 1.5839431 | 8.4055E-05 |
| ENSG00000186111 | PIP5K1C | 1.5818681 | 0.00026203 |
| ENSG00000161955 | TNFSF13 | 1.58119351 | 0.00020436 |
| ENSG00000158467 | AHCYL2 | 1.57606659 | 0.00011131 |
| ENSG00000186063 | AIDA | 1.57409347 | 0.00046529 |
| ENSG00000178537 | SLC25A20 | 1.57353802 | 0.00081597 |
| ENSG00000111667 | USP5 | 1.57295741 | 0.00031694 |
| ENSG00000131844 | MCCC2 | 1.57282243 | 0.0002418 |
| ENSG00000143119 | CD53 | 1.57109935 | 0.00021484 |
| ENSG00000160211 | G6PD | 1.56978257 | 0.000601 |
| ENSG00000143382 | ADAMTSL4 | 1.56948514 | 0.00264256 |
| ENSG00000070814 | TCOF1 | 1.56868052 | 0.00047455 |
| ENSG00000173193 | PARP14 | 1.56740882 | 0.00445351 |
| ENSG00000180353 | HCLS1 | 1.56664584 | 0.00021343 |
| ENSG00000115904 | SOS1 | 1.56418694 | 0.0009446 |

| | | | |
|---|---|---|---|
| ENSG00000049239 | H6PD | 1.56086241 | 0.0023701 |
| ENSG00000166145 | SPINT1 | 1.56056375 | 0.00019592 |
| ENSG00000175662 | TOM1L2 | 1.56043927 | 0.00019192 |
| ENSG00000108679 | LGALS3BP | 1.56029735 | 0.01089084 |
| ENSG00000107736 | CDH23 | 1.5591635 | 0.00195481 |
| ENSG00000058866 | DGKG | 1.55747451 | 0.00087231 |
| ENSG00000138814 | PPP3CA | 1.55720597 | 0.00079219 |
| ENSG00000134851 | TMEM165 | 1.55709643 | 3.69E-05 |
| ENSG00000137509 | PRCP | 1.55445473 | 5.2936E-05 |
| ENSG00000164023 | SGMS2 | 1.54486665 | 0.00023497 |
| ENSG00000070190 | DAPP1 | 1.54042513 | 0.00052724 |
| ENSG00000136305 | CIDEB | 1.53982943 | 0.00035831 |
| ENSG00000162736 | NCSTN | 1.53913214 | 0.00014247 |
| ENSG00000160883 | HK3 | 1.53712863 | 0.00139367 |
| ENSG00000146070 | PLA2G7 | 1.53676805 | 0.0003987 |
| ENSG00000100030 | MAPK1 | 1.53661952 | 0.00038097 |
| ENSG00000143933 | CALM2 | 1.53556489 | 0.00061541 |
| ENSG00000173786 | CNP | 1.53534972 | 0.00189816 |
| ENSG00000117228 | GBP1 | 1.53441652 | 0.02972753 |
| ENSG00000172269 | DPAGT1 | 1.53404302 | 0.00017204 |
| ENSG00000151151 | IPMK | 1.53347652 | 0.01739805 |
| ENSG00000077380 | DYNC1I2 | 1.53322432 | 0.00032015 |
| ENSG00000100201 | DDX17 | 1.53199949 | 0.00146578 |
| ENSG00000164068 | RNF123 | 1.52989919 | 0.01662566 |
| ENSG00000130429 | ARPC1B | 1.52804571 | 0.00173207 |
| ENSG00000135905 | DOCK10 | 1.52795493 | 0.00066095 |
| ENSG00000138639 | ARHGAP24 | 1.52793953 | 0.00030893 |
| ENSG00000111540 | RAB5B | 1.52701183 | 0.00026203 |
| ENSG00000163840 | DTX3L | 1.52401629 | 0.00057399 |
| ENSG00000164125 | FAM198B | 1.51862512 | 0.00842223 |
| ENSG00000238227 | TMEM250 | 1.51500264 | 0.00056375 |
| ENSG00000196684 | HSH2D | 1.51431852 | 5.6748E-05 |
| ENSG00000175857 | GAPT | 1.5109292 | 0.00103798 |
| ENSG00000160310 | PRMT2 | 1.51058035 | 0.00063722 |
| ENSG00000148484 | RSU1 | 1.50881568 | 0.00104815 |
| ENSG00000131018 | SYNE1 | 1.50740295 | 0.00289354 |
| ENSG00000163154 | TNFAIP8L2 | 1.5067246 | 0.00059557 |
| ENSG00000007923 | DNAJC11 | 1.50596034 | 0.00047624 |
| ENSG00000147471 | PLPBP | 1.50432034 | 8.2108E-05 |
| ENSG00000146376 | ARHGAP18 | 1.5025789 | 0.00011131 |
| ENSG00000067704 | IARS2 | 1.50016081 | 0.0001848 |
| ENSG00000169429 | CXCL8 | -5.6337326 | 0.00464143 |

| | | | |
|---|---|---|---|
| ENSG00000123689 | G0S2 | -5.3324527 | 0.01637209 |
| ENSG00000137801 | THBS1 | -5.2549864 | 0.01454249 |
| ENSG00000148344 | PTGES | -4.7753901 | 0.00509201 |
| ENSG00000162496 | DHRS3 | -4.498196 | 0.00892293 |
| ENSG00000112137 | PHACTR1 | -4.4357644 | 0.02363157 |
| ENSG00000184205 | TSPYL2 | -4.2134831 | 0.00390017 |
| ENSG00000081041 | CXCL2 | -4.1267357 | 0.01287257 |
| ENSG00000102760 | RGCC | -4.0251224 | 0.020498 |
| ENSG00000176597 | B3GNT5 | -3.9922784 | 0.00753884 |
| ENSG00000118971 | CCND2 | -3.9763733 | 0.02016236 |
| ENSG00000178726 | THBD | -3.9694314 | 0.03090869 |
| ENSG00000139112 | GABARAPL1 | -3.9174867 | 0.00344943 |
| ENSG00000163376 | KBTBD8 | -3.7481036 | 0.01002672 |
| ENSG00000160789 | LMNA | -3.59249 | 0.03524042 |
| ENSG00000100644 | HIF1A | -3.5458608 | 0.00057401 |
| ENSG00000153207 | AHCTF1 | -3.5254676 | 0.0084494 |
| ENSG00000008083 | JARID2 | -3.3847124 | 0.00115696 |
| ENSG00000182782 | HCAR2 | -3.3290246 | 0.02444392 |
| ENSG00000188042 | ARL4C | -3.3023161 | 0.03434752 |
| ENSG00000125740 | FOSB | -3.2798454 | 0.01611739 |
| ENSG00000126351 | THRA | -3.2650285 | 0.00891857 |
| ENSG00000124145 | SDC4 | -3.261413 | 0.00214841 |
| ENSG00000118985 | ELL2 | -3.2453856 | 0.03585554 |
| ENSG00000113369 | ARRDC3 | -3.2098649 | 0.0063002 |
| ENSG00000171604 | CXXC5 | -3.2020789 | 0.0162197 |
| ENSG00000113448 | PDE4D | -3.1957399 | 0.02085715 |
| ENSG00000169508 | GPR183 | -3.1681931 | 0.00722174 |
| ENSG00000277117 | FP565260.3 | -3.1326512 | 0.00510494 |
| ENSG00000166920 | C15orf48 | -3.1238849 | 0.01391301 |
| ENSG00000153234 | NR4A2 | -3.1148471 | 0.00604553 |
| ENSG00000078804 | TP53INP2 | -3.1113741 | 0.00575271 |
| ENSG00000168036 | CTNNB1 | -3.1044079 | 0.00037088 |
| ENSG00000130340 | SNX9 | -3.0836264 | 0.00821045 |
| ENSG00000110046 | ATG2A | -3.082018 | 0.00672133 |
| ENSG00000255398 | HCAR3 | -3.0667303 | 0.00655629 |
| ENSG00000088826 | SMOX | -3.0513615 | 0.00168846 |
| ENSG00000008056 | SYN1 | -3.017608 | 0.00202534 |
| ENSG00000151014 | NOCT | -3.0104672 | 0.0001848 |
| ENSG00000196878 | LAMB3 | -2.9950369 | 0.03525671 |
| ENSG00000120063 | GNA13 | -2.9950306 | 0.00263842 |
| ENSG00000165029 | ABCA1 | -2.9948132 | 0.00923629 |
| ENSG00000231721 | LINC-PINT | -2.9796939 | 0.00123023 |

| | | | |
|---|---|---|---|
| ENSG00000112715 | VEGFA | -2.9746894 | 0.00734878 |
| ENSG00000173575 | CHD2 | -2.9705471 | 0.0001277 |
| ENSG00000113070 | HBEGF | -2.9627795 | 0.00106234 |
| ENSG00000169155 | ZBTB43 | -2.9596239 | 0.00175002 |
| ENSG00000137507 | LRRC32 | -2.9517331 | 0.00049759 |
| ENSG00000106089 | STX1A | -2.9492251 | 0.01038173 |
| ENSG00000104312 | RIPK2 | -2.9480134 | 5.2951E-05 |
| ENSG00000075426 | FOSL2 | -2.9416515 | 0.0194375 |
| ENSG00000171174 | RBKS | -2.9359671 | 0.0252909 |
| ENSG00000271614 | ATP2B1-AS1 | -2.9328893 | 0.0007034 |
| ENSG00000173281 | PPP1R3B | -2.9287612 | 0.03212843 |
| ENSG00000185477 | GPRIN3 | -2.9252603 | 0.0011287 |
| ENSG00000185022 | MAFF | -2.9154679 | 0.03122127 |
| ENSG00000128271 | ADORA2A | -2.9125523 | 0.00079208 |
| ENSG00000155252 | PI4K2A | -2.9113472 | 0.00361432 |
| ENSG00000070961 | ATP2B1 | -2.8902578 | 0.03972003 |
| ENSG00000113742 | CPEB4 | -2.8816276 | 0.01257684 |
| ENSG00000160223 | ICOSLG | -2.8795385 | 0.00645006 |
| ENSG00000184588 | PDE4B | -2.8711138 | 0.00128813 |
| ENSG00000244242 | IFITM10 | -2.870418 | 0.00188837 |
| ENSG00000180530 | NRIP1 | -2.8643287 | 0.00170163 |
| ENSG00000166949 | SMAD3 | -2.8640214 | 0.00335679 |
| ENSG00000134070 | IRAK2 | -2.8624185 | 0.01250651 |
| ENSG00000099985 | OSM | -2.8597775 | 0.00029425 |
| ENSG00000144476 | ACKR3 | -2.8573954 | 0.00741895 |
| ENSG00000086062 | B4GALT1 | -2.8457697 | 0.00096056 |
| ENSG00000160271 | RALGDS | -2.8352271 | 0.00840216 |
| ENSG00000161921 | CXCL16 | -2.8324701 | 0.00334081 |
| ENSG00000108828 | VAT1 | -2.8268836 | 0.00453696 |
| ENSG00000134531 | EMP1 | -2.7956852 | 0.0093593 |
| ENSG00000112149 | CD83 | -2.7844651 | 0.00630094 |
| ENSG00000132819 | RBM38 | -2.7792212 | 0.00700209 |
| ENSG00000108179 | PPIF | -2.7523932 | 0.01740293 |
| ENSG00000133657 | ATP13A3 | -2.7405907 | 0.00591523 |
| ENSG00000128342 | LIF | -2.725365 | 0.02156661 |
| ENSG00000183696 | UPP1 | -2.7206894 | 3.816E-06 |
| ENSG00000011422 | PLAUR | -2.7006523 | 0.01328449 |
| ENSG00000198742 | SMURF1 | -2.6925219 | 0.00011131 |
| ENSG00000115548 | KDM3A | -2.6723105 | 0.00034172 |
| ENSG00000122644 | ARL4A | -2.667149 | 0.00045749 |
| ENSG00000116604 | MEF2D | -2.6511897 | 0.01193626 |
| ENSG00000143622 | RIT1 | -2.6418557 | 0.01017276 |

| | | | |
|---|---|---|---|
| ENSG00000133639 | BTG1 | -2.6096684 | 0.00691552 |
| ENSG00000057657 | PRDM1 | -2.5991526 | 0.00025626 |
| ENSG00000105835 | NAMPT | -2.5885866 | 0.01033927 |
| ENSG00000161835 | GRASP | -2.5845808 | 0.0444488 |
| ENSG00000139572 | GPR84 | -2.5804233 | 0.03542708 |
| ENSG00000060558 | GNA15 | -2.5766184 | 0.014623 |
| ENSG00000059804 | SLC2A3 | -2.5444028 | 0.00359548 |
| ENSG00000140332 | TLE3 | -2.5403499 | 0.00424621 |
| ENSG00000137309 | HMGA1 | -2.5322661 | 0.00031626 |
| ENSG00000234290 | AC116366.1 | -2.5197718 | 0.03965652 |
| ENSG00000055483 | USP36 | -2.5173927 | 0.00376868 |
| ENSG00000095794 | CREM | -2.5126091 | 0.02652157 |
| ENSG00000162711 | NLRP3 | -2.5091521 | 0.00029457 |
| ENSG00000025156 | HSF2 | -2.4950352 | 0.00038666 |
| ENSG00000170525 | PFKFB3 | -2.4903444 | 4.0167E-05 |
| ENSG00000132952 | USPL1 | -2.4856584 | 0.00113974 |
| ENSG00000189067 | LITAF | -2.4772791 | 0.02578617 |
| ENSG00000153094 | BCL2L11 | -2.4749415 | 0.00351515 |
| ENSG00000162889 | MAPKAPK2 | -2.4565655 | 0.00581231 |
| ENSG00000130202 | NECTIN2 | -2.4542862 | 0.03099256 |
| ENSG00000153922 | CHD1 | -2.4384453 | 0.000185 |
| ENSG00000162772 | ATF3 | -2.4341496 | 0.00106477 |
| ENSG00000137409 | MTCH1 | -2.4266198 | 0.00096657 |
| ENSG00000004660 | CAMKK1 | -2.4208654 | 0.00010145 |
| ENSG00000211445 | GPX3 | -2.4189819 | 0.02016236 |
| ENSG00000239305 | RNF103 | -2.4170593 | 1.2095E-05 |
| ENSG00000121966 | CXCR4 | -2.4156745 | 0.00711902 |
| ENSG00000141503 | MINK1 | -2.4153969 | 0.00443835 |
| ENSG00000177426 | TGIF1 | -2.4146208 | 0.00170844 |
| ENSG00000085117 | CD82 | -2.4142602 | 0.00082171 |
| ENSG00000120705 | ETF1 | -2.4137603 | 0.00115407 |
| ENSG00000143862 | ARL8A | -2.4017085 | 0.00690308 |
| ENSG00000154640 | BTG3 | -2.3777685 | 0.00037088 |
| ENSG00000110721 | CHKA | -2.3690688 | 0.00280401 |
| ENSG00000175352 | NRIP3 | -2.3658043 | 0.00702715 |
| ENSG00000104419 | NDRG1 | -2.3649559 | 0.00786721 |
| ENSG00000166579 | NDEL1 | -2.3578224 | 0.00828844 |
| ENSG00000105851 | PIK3CG | -2.3529044 | 0.00109512 |
| ENSG00000144597 | EAF1 | -2.3476008 | 0.00638678 |
| ENSG00000158470 | B4GALT5 | -2.3460849 | 0.00018367 |
| ENSG00000153201 | RANBP2 | -2.3460562 | 0.00176715 |
| ENSG00000116574 | RHOU | -2.3456583 | 0.00530599 |

| | | | |
|---|---|---|---|
| ENSG00000143153 | ATP1B1 | -2.3436801 | 0.00077849 |
| ENSG00000136603 | SKIL | -2.3399632 | 0.01964359 |
| ENSG00000112511 | PHF1 | -2.3395331 | 0.00017634 |
| ENSG00000177374 | HIC1 | -2.3365351 | 0.04264038 |
| ENSG00000130024 | PHF10 | -2.3349711 | 0.00376857 |
| ENSG00000179361 | ARID3B | -2.3322307 | 0.00836551 |
| ENSG00000143851 | PTPN7 | -2.3311657 | 0.00019919 |
| ENSG00000272888 | LINC01578 | -2.3298954 | 0.00039024 |
| ENSG00000146457 | WTAP | -2.329158 | 0.00024688 |
| ENSG00000204178 | TMEM57 | -2.3207445 | 0.00024688 |
| ENSG00000182831 | C16orf72 | -2.3189323 | 0.00687232 |
| ENSG00000165233 | CARD19 | -2.3086297 | 0.00512586 |
| ENSG00000070808 | CAMK2A | -2.3083899 | 0.02065956 |
| ENSG00000166747 | AP1G1 | -2.3035702 | 3.2365E-05 |
| ENSG00000158882 | TOMM40L | -2.3034862 | 0.00333586 |
| ENSG00000173812 | EIF1 | -2.2977993 | 5.2951E-05 |
| ENSG00000173334 | TRIB1 | -2.289118 | 0.00037874 |
| ENSG00000161714 | PLCD3 | -2.2762185 | 0.00120954 |
| ENSG00000110042 | DTX4 | -2.2758733 | 0.01231379 |
| ENSG00000103966 | EHD4 | -2.2706792 | 0.00220767 |
| ENSG00000256235 | SMIM3 | -2.2670807 | 0.00633024 |
| ENSG00000137331 | IER3 | -2.2625199 | 9.7018E-05 |
| ENSG00000186594 | MIR22HG | -2.2617682 | 0.01271882 |
| ENSG00000147872 | PLIN2 | -2.2604177 | 0.01529729 |
| ENSG00000135094 | SDS | -2.2587265 | 0.00393322 |
| ENSG00000154710 | RABGEF1 | -2.2579161 | 0.00573577 |
| ENSG00000147454 | SLC25A37 | -2.2524847 | 0.00808829 |
| ENSG00000054967 | RELT | -2.2509272 | 0.00401152 |
| ENSG00000131669 | NINJ1 | -2.2499543 | 0.00217054 |
| ENSG00000182149 | IST1 | -2.2486295 | 0.00016228 |
| ENSG00000015568 | RGPD5 | -2.2473691 | 0.02185054 |
| ENSG00000136717 | BIN1 | -2.2422262 | 0.00011131 |
| ENSG00000213923 | CSNK1E | -2.2412193 | 0.00076375 |
| ENSG00000005379 | TSPOAP1 | -2.2345228 | 0.02347435 |
| ENSG00000164211 | STARD4 | -2.2336266 | 0.01070554 |
| ENSG00000165804 | ZNF219 | -2.2328169 | 0.00045548 |
| ENSG00000175606 | TMEM70 | -2.2287584 | 0.04537919 |
| ENSG00000182957 | SPATA13 | -2.2211987 | 0.00342977 |
| ENSG00000164056 | SPRY1 | -2.2195278 | 0.01833939 |
| ENSG00000164715 | LMTK2 | -2.2135726 | 0.00079208 |
| ENSG00000090339 | ICAM1 | -2.2057344 | 0.00461745 |
| ENSG00000174437 | ATP2A2 | -2.2038848 | 0.00111639 |

| | | | |
|---|---|---|---|
| ENSG00000068878 | PSME4 | -2.2033136 | 0.00114568 |
| ENSG00000109066 | TMEM104 | -2.1988191 | 0.00428046 |
| ENSG00000141506 | PIK3R5 | -2.1952702 | 0.00129345 |
| ENSG00000121671 | CRY2 | -2.1947757 | 0.03182714 |
| ENSG00000165006 | UBAP1 | -2.1913766 | 4.6415E-05 |
| ENSG00000086666 | ZFAND6 | -2.1889773 | 0.00018529 |
| ENSG00000136732 | GYPC | -2.1856152 | 0.00024683 |
| ENSG00000213639 | PPP1CB | -2.1835705 | 0.00348599 |
| ENSG00000152102 | FAM168B | -2.1803401 | 0.00888977 |
| ENSG00000116679 | IVNS1ABP | -2.1759616 | 0.03850762 |
| ENSG00000170989 | S1PR1 | -2.1745131 | 0.01271455 |
| ENSG00000106546 | AHR | -2.1643124 | 0.00319666 |
| ENSG00000105722 | ERF | -2.1626278 | 0.00134062 |
| ENSG00000124198 | ARFGEF2 | -2.1572501 | 0.00049702 |
| ENSG00000166128 | RAB8B | -2.1568523 | 0.00023416 |
| ENSG00000270681 | AC095055.1 | -2.1564927 | 0.00334425 |
| ENSG00000057704 | TMCC3 | -2.155854 | 0.03222085 |
| ENSG00000204977 | TRIM13 | -2.1521171 | 0.00013559 |
| ENSG00000169629 | RGPD8 | -2.1506083 | 0.02394401 |
| ENSG00000169251 | NMD3 | -2.1366503 | 0.0000369 |
| ENSG00000107968 | MAP3K8 | -2.1358555 | 0.01085837 |
| ENSG00000101782 | RIOK3 | -2.1322235 | 0.00026591 |
| ENSG00000155307 | SAMSN1 | -2.1298181 | 0.03896169 |
| ENSG00000106635 | BCL7B | -2.1270893 | 0.00016228 |
| ENSG00000173848 | NET1 | -2.1189954 | 0.00079537 |
| ENSG00000197170 | PSMD12 | -2.1151412 | 0.00032825 |
| ENSG00000119950 | MXI1 | -2.1135775 | 0.02516512 |
| ENSG00000114784 | EIF1B | -2.1078115 | 0.00079091 |
| ENSG00000110047 | EHD1 | -2.1013485 | 0.00697519 |
| ENSG00000152484 | USP12 | -2.1003625 | 0.00089264 |
| ENSG00000183876 | ARSI | -2.099386 | 0.0013479 |
| ENSG00000166839 | ANKDD1A | -2.099097 | 0.01754524 |
| ENSG00000117000 | RLF | -2.0980237 | 0.00075679 |
| ENSG00000175130 | MARCKSL1 | -2.0916263 | 0.00105513 |
| ENSG00000103365 | GGA2 | -2.0901133 | 0.00121006 |
| ENSG00000102034 | ELF4 | -2.0898317 | 0.00326607 |
| ENSG00000173846 | PLK3 | -2.0895813 | 0.00113894 |
| ENSG00000110848 | CD69 | -2.0733829 | 0.01675735 |
| ENSG00000168264 | IRF2BP2 | -2.071314 | 0.0026279 |
| ENSG00000122877 | EGR2 | -2.0695338 | 0.00800203 |
| ENSG00000110367 | DDX6 | -2.0645479 | 0.00088717 |
| ENSG00000078269 | SYNJ2 | -2.0629506 | 0.02118512 |

| | | | |
|---|---|---|---|
| ENSG00000263020 | AL662899.2 | -2.0572586 | 0.04147729 |
| ENSG00000102265 | TIMP1 | -2.0567885 | 0.00349848 |
| ENSG00000100614 | PPM1A | -2.0557918 | 8.4055E-05 |
| ENSG00000177706 | FAM20C | -2.0534868 | 0.028623 |
| ENSG00000126705 | AHDC1 | -2.0525454 | 0.01052401 |
| ENSG00000125733 | TRIP10 | -2.0485752 | 0.01502992 |
| ENSG00000198805 | PNP | -2.0454355 | 0.03460549 |
| ENSG00000186431 | FCAR | -2.0449644 | 0.03493191 |
| ENSG00000145675 | PIK3R1 | -2.0436209 | 0.0444155 |
| ENSG00000145632 | PLK2 | -2.0431337 | 0.03173833 |
| ENSG00000149115 | TNKS1BP1 | -2.0344923 | 0.00977956 |
| ENSG00000020633 | RUNX3 | -2.0304191 | 0.00303617 |
| ENSG00000132326 | PER2 | -2.0278655 | 0.00143466 |
| ENSG00000135404 | CD63 | -2.0257583 | 0.00303568 |
| ENSG00000155304 | HSPA13 | -2.0245682 | 0.00124066 |
| ENSG00000171316 | CHD7 | -2.0198256 | 0.0001798 |
| ENSG00000116786 | PLEKHM2 | -2.0185337 | 0.00365448 |
| ENSG00000101413 | RPRD1B | -2.0152206 | 0.00032015 |
| ENSG00000173744 | AGFG1 | -2.0128491 | 0.00367787 |
| ENSG00000171867 | PRNP | -2.008113 | 0.00149954 |
| ENSG00000153914 | SREK1 | -2.0060482 | 0.00187327 |
| ENSG00000185650 | ZFP36L1 | -2.0056671 | 0.01145874 |
| ENSG00000186174 | BCL9L | -2.002974 | 0.01942106 |
| ENSG00000168214 | RBPJ | -2.0013176 | 0.00293188 |
| ENSG00000023287 | RB1CC1 | -1.9931023 | 0.00702366 |
| ENSG00000132661 | NXT1 | -1.9919591 | 0.04292691 |
| ENSG00000204160 | ZDHHC18 | -1.9917883 | 0.00149954 |
| ENSG00000172071 | EIF2AK3 | -1.9917504 | 0.00253835 |
| ENSG00000153066 | TXNDC11 | -1.9897945 | 0.00119443 |
| ENSG00000176845 | METRNL | -1.9892546 | 0.0068429 |
| ENSG00000186660 | ZFP91 | -1.9889859 | 0.0000369 |
| ENSG00000069956 | MAPK6 | -1.9881321 | 0.00203372 |
| ENSG00000107816 | LZTS2 | -1.9862474 | 0.00036847 |
| ENSG00000123091 | RNF11 | -1.9860266 | 2.4517E-05 |
| ENSG00000196923 | PDLIM7 | -1.9855536 | 0.00080795 |
| ENSG00000033327 | GAB2 | -1.9828257 | 0.00425917 |
| ENSG00000139832 | RAB20 | -1.9828184 | 0.00775912 |
| ENSG00000174738 | NR1D2 | -1.9794011 | 0.00432981 |
| ENSG00000186642 | PDE2A | -1.9782233 | 0.02931326 |
| ENSG00000198900 | TOP1 | -1.9740894 | 0.0079006 |
| ENSG00000145241 | CENPC | -1.9740236 | 0.00061876 |
| ENSG00000107554 | DNMBP | -1.9723214 | 0.01715119 |

| | | | |
|---|---|---|---|
| ENSG00000183735 | TBK1 | -1.9713113 | 1.8312E-05 |
| ENSG00000100083 | GGA1 | -1.9684336 | 0.0001054 |
| ENSG00000165030 | NFIL3 | -1.9683506 | 0.0001798 |
| ENSG00000163877 | SNIP1 | -1.9677867 | 0.00134844 |
| ENSG00000110713 | NUP98 | -1.9636771 | 0.00253039 |
| ENSG00000125812 | GZF1 | -1.9620554 | 0.02372848 |
| ENSG00000110852 | CLEC2B | -1.9618504 | 0.00497045 |
| ENSG00000165424 | ZCCHC24 | -1.9606687 | 0.00861902 |
| ENSG00000118515 | SGK1 | -1.9587755 | 0.03928864 |
| ENSG00000111641 | NOP2 | -1.9578295 | 0.030491 |
| ENSG00000126368 | NR1D1 | -1.9572148 | 0.01145314 |
| ENSG00000117139 | KDM5B | -1.9569944 | 0.00217905 |
| ENSG00000134107 | BHLHE40 | -1.9566723 | 0.01264053 |
| ENSG00000111615 | KRR1 | -1.954116 | 0.0008326 |
| ENSG00000280138 | AC027290.2 | -1.9507009 | 0.00039116 |
| ENSG00000083799 | CYLD | -1.950076 | 0.00022505 |
| ENSG00000137166 | FOXP4 | -1.949993 | 0.00140031 |
| ENSG00000145780 | FEM1C | -1.9471062 | 0.00023497 |
| ENSG00000082153 | BZW1 | -1.9431389 | 0.01711668 |
| ENSG00000107863 | ARHGAP21 | -1.9402569 | 6.1348E-05 |
| ENSG00000036054 | TBC1D23 | -1.9309444 | 0.00022218 |
| ENSG00000133789 | SWAP70 | -1.9304165 | 0.00333076 |
| ENSG00000123360 | PDE1B | -1.9295376 | 0.00186871 |
| ENSG00000186918 | ZNF395 | -1.9273649 | 0.00392283 |
| ENSG00000125772 | GPCPD1 | -1.9239971 | 0.00387853 |
| ENSG00000170638 | TRABD | -1.9202134 | 0.00016159 |
| ENSG00000083896 | YTHDC1 | -1.9190759 | 8.2108E-05 |
| ENSG00000064012 | CASP8 | -1.912928 | 0.00168434 |
| ENSG00000184216 | IRAK1 | -1.9121738 | 0.01178924 |
| ENSG00000253276 | CCDC71L | -1.911114 | 0.00254656 |
| ENSG00000099381 | SETD1A | -1.9071814 | 0.02676969 |
| ENSG00000162924 | REL | -1.9040934 | 3.2365E-05 |
| ENSG00000154978 | VOPP1 | -1.9028077 | 0.01537657 |
| ENSG00000055609 | KMT2C | -1.8989444 | 0.02105759 |
| ENSG00000160741 | CRTC2 | -1.8987936 | 0.0002735 |
| ENSG00000240849 | TMEM189 | -1.8954654 | 0.01891436 |
| ENSG00000196233 | LCOR | -1.8941411 | 0.00006077 |
| ENSG00000079332 | SAR1A | -1.8930478 | 0.00037122 |
| ENSG00000019995 | ZRANB1 | -1.8898784 | 0.00046556 |
| ENSG00000109113 | RAB34 | -1.8894695 | 0.0012744 |
| ENSG00000101558 | VAPA | -1.8860457 | 0.04326385 |
| ENSG00000101421 | CHMP4B | -1.8811711 | 0.00179355 |

| | | | |
|---|---|---|---|
| ENSG00000163785 | RYK | -1.8750924 | 0.00092722 |
| ENSG00000272886 | DCP1A | -1.8731265 | 0.00157429 |
| ENSG00000118689 | FOXO3 | -1.8706899 | 0.00959379 |
| ENSG00000121749 | TBC1D15 | -1.8704506 | 0.00049264 |
| ENSG00000196843 | ARID5A | -1.8689803 | 0.00450279 |
| ENSG00000211459 | MT-RNR1 | -1.8644215 | 0.00135837 |
| ENSG00000173276 | ZBTB21 | -1.8617309 | 0.00147052 |
| ENSG00000181045 | SLC26A11 | -1.8598453 | 0.01978584 |
| ENSG00000112406 | HECA | -1.8598173 | 0.00092214 |
| ENSG00000105656 | ELL | -1.8583687 | 0.00732956 |
| ENSG00000112282 | MED23 | -1.8571229 | 0.00162936 |
| ENSG00000138166 | DUSP5 | -1.8569247 | 0.00086712 |
| ENSG00000169762 | TAPT1 | -1.8516965 | 0.0003077 |
| ENSG00000163050 | COQ8A | -1.8501052 | 0.00675146 |
| ENSG00000126775 | ATG14 | -1.8468722 | 0.00024848 |
| ENSG00000132912 | DCTN4 | -1.8468371 | 0.00010145 |
| ENSG00000100284 | TOM1 | -1.846443 | 0.00153216 |
| ENSG00000167996 | FTH1 | -1.8459891 | 0.00235438 |
| ENSG00000023734 | STRAP | -1.8458261 | 0.0000369 |
| ENSG00000183955 | KMT5A | -1.8436599 | 0.00498639 |
| ENSG00000186834 | HEXIM1 | -1.8421121 | 0.00571373 |
| ENSG00000166165 | CKB | -1.8411489 | 0.00802101 |
| ENSG00000065809 | FAM107B | -1.8409382 | 0.00187327 |
| ENSG00000198853 | RUSC2 | -1.8374819 | 0.00601804 |
| ENSG00000160570 | DEDD2 | -1.8355286 | 0.00329172 |
| ENSG00000181220 | ZNF746 | -1.8354554 | 0.00622101 |
| ENSG00000008952 | SEC62 | -1.8310351 | 0.00041379 |
| ENSG00000169967 | MAP3K2 | -1.8276966 | 0.01488026 |
| ENSG00000258890 | CEP95 | -1.8171787 | 0.00297137 |
| ENSG00000167491 | GATAD2A | -1.8153259 | 0.00293188 |
| ENSG00000122068 | FYTTD1 | -1.8128755 | 0.0002456 |
| ENSG00000132155 | RAF1 | -1.8089564 | 0.00063722 |
| ENSG00000168438 | CDC40 | -1.8080239 | 0.00339332 |
| ENSG00000255112 | CHMP1B | -1.8052809 | 0.00601804 |
| ENSG00000140564 | FURIN | -1.7987076 | 0.01662645 |
| ENSG00000140941 | MAP1LC3B | -1.7985835 | 0.00108044 |
| ENSG00000156650 | KAT6B | -1.7985556 | 0.00043898 |
| ENSG00000103657 | HERC1 | -1.7963983 | 0.00025626 |
| ENSG00000132510 | KDM6B | -1.795885 | 0.0001275 |
| ENSG00000176624 | MEX3C | -1.7956592 | 0.00010911 |
| ENSG00000104081 | BMF | -1.7933229 | 0.0037225 |
| ENSG00000101544 | ADNP2 | -1.7924296 | 0.00071567 |

| | | | |
|---|---|---|---|
| ENSG00000047634 | SCML1 | -1.7920938 | 0.00018529 |
| ENSG00000196396 | PTPN1 | -1.7911115 | 0.00848506 |
| ENSG00000101216 | GMEB2 | -1.7898481 | 0.00651799 |
| ENSG00000170677 | SOCS6 | -1.7869862 | 0.01133185 |
| ENSG00000106723 | SPIN1 | -1.786127 | 0.00392897 |
| ENSG00000138069 | RAB1A | -1.7825402 | 0.0046341 |
| ENSG00000141580 | WDR45B | -1.7784459 | 0.00614734 |
| ENSG00000139725 | RHOF | -1.7777834 | 0.00807752 |
| ENSG00000126524 | SBDS | -1.7750369 | 0.00087231 |
| ENSG00000183431 | SF3A3 | -1.7741834 | 0.0004242 |
| ENSG00000164687 | FABP5 | -1.772035 | 0.00229042 |
| ENSG00000162664 | ZNF326 | -1.7703503 | 0.00016166 |
| ENSG00000164442 | CITED2 | -1.7700743 | 0.00254922 |
| ENSG00000144655 | CSRNP1 | -1.7700244 | 0.01231379 |
| ENSG00000160408 | ST6GALNAC6 | -1.7689971 | 0.00179958 |
| ENSG00000109118 | PHF12 | -1.7661587 | 0.00072397 |
| ENSG00000106346 | USP42 | -1.7657116 | 0.00231527 |
| ENSG00000102984 | ZNF821 | -1.7654559 | 0.00205786 |
| ENSG00000101384 | JAG1 | -1.7652708 | 0.00023497 |
| ENSG00000132334 | PTPRE | -1.7617087 | 0.00829848 |
| ENSG00000100425 | BRD1 | -1.7605867 | 0.0001505 |
| ENSG00000127838 | PNKD | -1.7604756 | 0.03950871 |
| ENSG00000178127 | NDUFV2 | -1.7603756 | 0.00073001 |
| ENSG00000164823 | OSGIN2 | -1.753502 | 0.00033773 |
| ENSG00000124762 | CDKN1A | -1.7524954 | 0.00250115 |
| ENSG00000197183 | NOL4L | -1.749076 | 0.02292124 |
| ENSG00000104885 | DOT1L | -1.7490404 | 0.00170844 |
| ENSG00000156860 | FBRS | -1.749037 | 0.0011299 |
| ENSG00000178789 | CD300LB | -1.7414884 | 0.02436108 |
| ENSG00000242125 | SNHG3 | -1.7411488 | 0.00077068 |
| ENSG00000072274 | TFRC | -1.7400163 | 0.04603645 |
| ENSG00000027697 | IFNGR1 | -1.7393537 | 0.0136665 |
| ENSG00000183484 | GPR132 | -1.7384703 | 0.00125237 |
| ENSG00000146872 | TLK2 | -1.738412 | 0.00011229 |
| ENSG00000139697 | SBNO1 | -1.7374031 | 0.00493368 |
| ENSG00000175556 | LONRF3 | -1.7374001 | 7.6536E-05 |
| ENSG00000131051 | RBM39 | -1.7358744 | 0.00069566 |
| ENSG00000090924 | PLEKHG2 | -1.7317967 | 0.00258906 |
| ENSG00000114796 | KLHL24 | -1.7293225 | 0.0001253 |
| ENSG00000268734 | AC245128.3 | -1.7288244 | 0.03527874 |
| ENSG00000159461 | AMFR | -1.7251031 | 0.00459037 |
| ENSG00000134815 | DHX34 | -1.7200533 | 0.00094463 |

| | | | |
|---|---|---|---|
| ENSG00000106004 | HOXA5 | -1.7157683 | 0.01131822 |
| ENSG00000174574 | AKIRIN1 | -1.7154807 | 0.00059211 |
| ENSG00000100100 | PIK3IP1 | -1.7144539 | 0.00698448 |
| ENSG00000119801 | YPEL5 | -1.7132154 | 0.0001146 |
| ENSG00000138593 | SECISBP2L | -1.7125599 | 0.00014906 |
| ENSG00000183337 | BCOR | -1.7104719 | 0.0451965 |
| ENSG00000080371 | RAB21 | -1.7089091 | 0.00018222 |
| ENSG00000008294 | SPAG9 | -1.7086175 | 0.00105353 |
| ENSG00000076924 | XAB2 | -1.7069058 | 0.00044978 |
| ENSG00000145495 | 38777 | -1.7043489 | 0.0003874 |
| ENSG00000140367 | UBE2Q2 | -1.7043335 | 0.00029716 |
| ENSG00000177169 | ULK1 | -1.701854 | 6.3157E-05 |
| ENSG00000164327 | RICTOR | -1.7017626 | 0.00133432 |
| ENSG00000163697 | APBB2 | -1.7010362 | 0.02373579 |
| ENSG00000150593 | PDCD4 | -1.7007338 | 0.00040152 |
| ENSG00000036257 | CUL3 | -1.7001997 | 0.00072261 |
| ENSG00000196428 | TSC22D2 | -1.6986638 | 0.00068123 |
| ENSG00000143217 | NECTIN4 | -1.698404 | 0.00022322 |
| ENSG00000213722 | DDAH2 | -1.6968848 | 0.0022176 |
| ENSG00000155926 | SLA | -1.6968623 | 0.03257164 |
| ENSG00000163605 | PPP4R2 | -1.695595 | 0.00032825 |
| ENSG00000169895 | SYAP1 | -1.6955365 | 0.01716858 |
| ENSG00000087206 | UIMC1 | -1.6949801 | 0.00035769 |
| ENSG00000013441 | CLK1 | -1.694474 | 0.00031514 |
| ENSG00000171988 | JMJD1C | -1.6918273 | 0.00078546 |
| ENSG00000075415 | SLC25A3 | -1.6912662 | 0.00151518 |
| ENSG00000124486 | USP9X | -1.6910736 | 0.00016587 |
| ENSG00000259330 | INAFM2 | -1.6891145 | 0.00217509 |
| ENSG00000188215 | DCUN1D3 | -1.6882633 | 0.00039781 |
| ENSG00000231259 | AC125232.1 | -1.6869602 | 0.01216003 |
| ENSG00000197818 | SLC9A8 | -1.6864555 | 0.01216003 |
| ENSG00000114120 | SLC25A36 | -1.6853343 | 0.00396268 |
| ENSG00000008256 | CYTH3 | -1.6823948 | 0.00671091 |
| ENSG00000102225 | CDK16 | -1.68182 | 0.0067314 |
| ENSG00000122862 | SRGN | -1.6814337 | 0.00363392 |
| ENSG00000150907 | FOXO1 | -1.6803792 | 0.0011299 |
| ENSG00000197063 | MAFG | -1.6756863 | 0.01274117 |
| ENSG00000067182 | TNFRSF1A | -1.6720402 | 1.2095E-05 |
| ENSG00000150457 | LATS2 | -1.6711013 | 0.00114234 |
| ENSG00000165813 | CCDC186 | -1.6705424 | 0.00225444 |
| ENSG00000166484 | MAPK7 | -1.6667525 | 0.0052913 |
| ENSG00000143514 | TP53BP2 | -1.6645397 | 0.00098261 |

| | | | |
|---|---|---|---|
| ENSG00000168615 | ADAM9 | -1.6638168 | 0.0008965 |
| ENSG00000135093 | USP30 | -1.6631448 | 0.00340911 |
| ENSG00000072364 | AFF4 | -1.6623323 | 3.816E-06 |
| ENSG00000102119 | EMD | -1.6621668 | 0.00070482 |
| ENSG00000214174 | AMZ2P1 | -1.6618303 | 0.00776543 |
| ENSG00000188994 | ZNF292 | -1.6564893 | 0.01019047 |
| ENSG00000126767 | ELK1 | -1.6562438 | 7.4197E-05 |
| ENSG00000131408 | NR1H2 | -1.6550241 | 0.00025405 |
| ENSG00000167703 | SLC43A2 | -1.6507063 | 0.01025104 |
| ENSG00000105325 | FZR1 | -1.6463611 | 0.00941259 |
| ENSG00000117036 | ETV3 | -1.6441059 | 0.0422585 |
| ENSG00000130517 | PGPEP1 | -1.640259 | 0.00108282 |
| ENSG00000159346 | ADIPOR1 | -1.6401029 | 0.00077849 |
| ENSG00000117569 | PTBP2 | -1.6397044 | 0.00083979 |
| ENSG00000118707 | TGIF2 | -1.6382022 | 0.00171347 |
| ENSG00000123066 | MED13L | -1.636608 | 0.00066794 |
| ENSG00000176170 | SPHK1 | -1.6363195 | 0.0084539 |
| ENSG00000130695 | CEP85 | -1.6359461 | 0.00590266 |
| ENSG00000102908 | NFAT5 | -1.6359083 | 0.00269425 |
| ENSG00000140044 | JDP2 | -1.6348555 | 0.01099759 |
| ENSG00000109332 | UBE2D3 | -1.6330588 | 0.00072397 |
| ENSG00000175197 | DDIT3 | -1.632968 | 0.033075 |
| ENSG00000135503 | ACVR1B | -1.6323848 | 0.0008674 |
| ENSG00000087087 | SRRT | -1.629347 | 0.01770152 |
| ENSG00000150403 | TMCO3 | -1.6286247 | 0.00081798 |
| ENSG00000178607 | ERN1 | -1.6281895 | 0.00018294 |
| ENSG00000105856 | HBP1 | -1.6271893 | 0.00005434 |
| ENSG00000267520 | AC010733.2 | -1.6271018 | 0.00235373 |
| ENSG00000140379 | BCL2A1 | -1.6264128 | 0.00184798 |
| ENSG00000197386 | HTT | -1.6249569 | 0.03190124 |
| ENSG00000280987 | MATR3 | -1.6242489 | 0.03360204 |
| ENSG00000056972 | TRAF3IP2 | -1.6234054 | 0.00461753 |
| ENSG00000111011 | RSRC2 | -1.6228625 | 0.00059557 |
| ENSG00000135968 | GCC2 | -1.6219996 | 0.00934996 |
| ENSG00000147526 | TACC1 | -1.6202835 | 0.0013507 |
| ENSG00000005486 | RHBDD2 | -1.6201412 | 0.01186484 |
| ENSG00000115165 | CYTIP | -1.6199146 | 0.00976103 |
| ENSG00000172766 | NAA16 | -1.6173609 | 0.00064718 |
| ENSG00000145860 | RNF145 | -1.6171896 | 0.00100657 |
| ENSG00000156381 | ANKRD9 | -1.6163393 | 0.02888984 |
| ENSG00000069399 | BCL3 | -1.6159176 | 0.01288218 |
| ENSG00000143373 | ZNF687 | -1.6152912 | 0.03419193 |

| | | | |
|---|---|---|---|
| ENSG00000116584 | ARHGEF2 | -1.6134978 | 0.01483446 |
| ENSG00000179051 | RCC2 | -1.6127476 | 0.01160789 |
| ENSG00000124766 | SOX4 | -1.6114084 | 0.00223409 |
| ENSG00000044574 | HSPA5 | -1.6099202 | 0.00492287 |
| ENSG00000145979 | TBC1D7 | -1.6092439 | 0.00847841 |
| ENSG00000139433 | GLTP | -1.6066986 | 0.00417663 |
| ENSG00000138867 | GUCD1 | -1.6063594 | 0.00169689 |
| ENSG00000198925 | ATG9A | -1.6058816 | 0.00784433 |
| ENSG00000032219 | ARID4A | -1.6058472 | 0.00016587 |
| ENSG00000241839 | PLEKHO2 | -1.6046012 | 0.00403476 |
| ENSG00000100221 | JOSD1 | -1.6035386 | 0.01667953 |
| ENSG00000114098 | ARMC8 | -1.6017833 | 7.5228E-05 |
| ENSG00000178623 | GPR35 | -1.6007657 | 0.0312656 |
| ENSG00000136830 | FAM129B | -1.5973004 | 0.00921876 |
| ENSG00000005483 | KMT2E | -1.5948804 | 0.00019371 |
| ENSG00000136807 | CDK9 | -1.5938694 | 0.00702732 |
| ENSG00000105655 | ISYNA1 | -1.5926978 | 0.00582601 |
| ENSG00000138434 | SSFA2 | -1.5911581 | 0.00017899 |
| ENSG00000115956 | PLEK | -1.591103 | 0.0025654 |
| ENSG00000055208 | TAB2 | -1.5910037 | 0.00017634 |
| ENSG00000120690 | ELF1 | -1.589289 | 0.0006017 |
| ENSG00000107937 | GTPBP4 | -1.5890128 | 0.00428685 |
| ENSG00000023330 | ALAS1 | -1.5889304 | 0.04777731 |
| ENSG00000115339 | GALNT3 | -1.587047 | 0.01936306 |
| ENSG00000152409 | JMY | -1.586805 | 0.00037088 |
| ENSG00000270069 | MIR222HG | -1.5859323 | 0.00096657 |
| ENSG00000116260 | QSOX1 | -1.5833351 | 0.00220839 |
| ENSG00000043093 | DCUN1D1 | -1.5832245 | 0.00075096 |
| ENSG00000197122 | SRC | -1.5803507 | 0.00058719 |
| ENSG00000135241 | PNPLA8 | -1.5795546 | 0.0024376 |
| ENSG00000124782 | RREB1 | -1.5782724 | 0.00106505 |
| ENSG00000127824 | TUBA4A | -1.5779967 | 0.00708238 |
| ENSG00000198369 | SPRED2 | -1.5763408 | 0.0016113 |
| ENSG00000151247 | EIF4E | -1.5752877 | 0.0004212 |
| ENSG00000173960 | UBXN2A | -1.5744389 | 0.00037874 |
| ENSG00000184428 | TOP1MT | -1.5739683 | 0.01145314 |
| ENSG00000240053 | LY6G5B | -1.5730019 | 0.00483369 |
| ENSG00000140743 | CDR2 | -1.5723227 | 0.00416262 |
| ENSG00000168066 | SF1 | -1.5711265 | 0.00248889 |
| ENSG00000160179 | ABCG1 | -1.5694762 | 0.00069117 |
| ENSG00000064961 | HMG20B | -1.5673559 | 0.00040591 |
| ENSG00000154124 | OTULIN | -1.5673035 | 0.00123872 |

| | | | |
|---|---|---|---|
| ENSG00000181222 | POLR2A | -1.566575 | 0.00159885 |
| ENSG00000076108 | BAZ2A | -1.5661032 | 0.00076885 |
| ENSG00000076604 | TRAF4 | -1.5658535 | 0.01836102 |
| ENSG00000074416 | MGLL | -1.5650616 | 0.02306784 |
| ENSG00000159082 | SYNJ1 | -1.5626197 | 0.00604553 |
| ENSG00000153561 | RMND5A | -1.5626103 | 0.00119944 |
| ENSG00000186469 | GNG2 | -1.5612618 | 0.01606603 |
| ENSG00000152700 | SAR1B | -1.5608848 | 0.00029457 |
| ENSG00000239857 | GET4 | -1.559559 | 0.0017433 |
| ENSG00000113712 | CSNK1A1 | -1.5593144 | 0.00022731 |
| ENSG00000260708 | AL118516.1 | -1.5586566 | 0.00633623 |
| ENSG00000102007 | PLP2 | -1.5573676 | 0.00035647 |
| ENSG00000112033 | PPARD | -1.5559943 | 0.00175002 |
| ENSG00000147439 | BIN3 | -1.555449 | 0.00188738 |
| ENSG00000198833 | UBE2J1 | -1.5532614 | 0.00794965 |
| ENSG00000137817 | PARP6 | -1.5530293 | 0.00011008 |
| ENSG00000105968 | H2AFV | -1.5522882 | 0.00025717 |
| ENSG00000117614 | SYF2 | -1.5513175 | 8.5435E-05 |
| ENSG00000155508 | CNOT8 | -1.5510955 | 0.001272 |
| ENSG00000162923 | WDR26 | -1.5504078 | 0.00013674 |
| ENSG00000116285 | ERRFI1 | -1.5503939 | 0.00773351 |
| ENSG00000164169 | PRMT9 | -1.5500579 | 0.00051771 |
| ENSG00000113575 | PPP2CA | -1.5499641 | 0.00062165 |
| ENSG00000197780 | TAF13 | -1.5496012 | 0.00091009 |
| ENSG00000262246 | CORO7 | -1.5483429 | 0.00958444 |
| ENSG00000166233 | ARIH1 | -1.5479744 | 0.00015176 |
| ENSG00000244486 | SCARF2 | -1.5463953 | 0.0210235 |
| ENSG00000128989 | ARPP19 | -1.5457656 | 0.00077102 |
| ENSG00000164543 | STK17A | -1.5455914 | 0.0392918 |
| ENSG00000131759 | RARA | -1.5448456 | 0.01590993 |
| ENSG00000276107 | AC037198.2 | -1.5440499 | 0.00238135 |
| ENSG00000139636 | LMBR1L | -1.5439465 | 0.00464824 |
| ENSG00000167604 | NFKBID | -1.5436091 | 0.00256234 |
| ENSG00000197405 | C5AR1 | -1.542184 | 0.00477562 |
| ENSG00000070495 | JMJD6 | -1.5394051 | 0.0030835 |
| ENSG00000092847 | AGO1 | -1.5393011 | 0.00526619 |
| ENSG00000105821 | DNAJC2 | -1.5391475 | 0.00675342 |
| ENSG00000119048 | UBE2B | -1.5368156 | 0.00043714 |
| ENSG00000156671 | SAMD8 | -1.5360163 | 0.00349533 |
| ENSG00000139505 | MTMR6 | -1.5350177 | 0.00115962 |
| ENSG00000164548 | TRA2A | -1.5338747 | 0.00011163 |
| ENSG00000029993 | HMGB3 | -1.5316349 | 0.01029999 |

| | | | |
|---|---|---|---|
| ENSG00000185947 | ZNF267 | -1.5298207 | 0.00454279 |
| ENSG00000273356 | LINC02019 | -1.5292778 | 0.00119944 |
| ENSG00000141985 | SH3GL1 | -1.5280713 | 0.00588359 |
| ENSG00000128272 | ATF4 | -1.5280042 | 5.2951E-05 |
| ENSG00000172216 | CEBPB | -1.527772 | 0.02014986 |
| ENSG00000158669 | GPAT4 | -1.5268625 | 0.00166041 |
| ENSG00000188070 | C11orf95 | -1.5266466 | 0.02506593 |
| ENSG00000111252 | SH2B3 | -1.5263865 | 0.00860229 |
| ENSG00000065357 | DGKA | -1.5260897 | 0.0002973 |
| ENSG00000167671 | UBXN6 | -1.5259656 | 0.00109223 |
| ENSG00000169905 | TOR1AIP2 | -1.5249436 | 0.00597502 |
| ENSG00000147119 | CHST7 | -1.5236811 | 0.00789381 |
| ENSG00000167034 | NKX3-1 | -1.5236105 | 0.00062255 |
| ENSG00000161638 | ITGA5 | -1.5223195 | 4.6944E-05 |
| ENSG00000154359 | LONRF1 | -1.5210642 | 0.00063116 |
| ENSG00000143067 | ZNF697 | -1.5168708 | 7.6536E-05 |
| ENSG00000230551 | AC021078.1 | -1.5167031 | 0.00229423 |
| ENSG00000166974 | MAPRE2 | -1.5133908 | 0.00017899 |
| ENSG00000179119 | SPTY2D1 | -1.512944 | 0.00042489 |
| ENSG00000067596 | DHX8 | -1.512099 | 0.00033588 |
| ENSG00000137876 | RSL24D1 | -1.5114195 | 0.00105092 |
| ENSG00000137947 | GTF2B | -1.510406 | 0.00011985 |
| ENSG00000108669 | CYTH1 | -1.505954 | 0.00110115 |
| ENSG00000149782 | PLCB3 | -1.5047599 | 0.00626591 |
| ENSG00000144711 | IQSEC1 | -1.50364 | 0.0020674 |
| ENSG00000133794 | ARNTL | -1.5018533 | 0.00050819 |
| ENSG00000067900 | ROCK1 | -1.5006315 | 0.00019992 |

**Supplemental Table 3. Flow cytometry antibody and reagents.**

| Antibody | Fluorocrhome | Clone | Ref | Company |
|---|---|---|---|---|
| Anti-mouse MHCII | APC/Fire750 | M5/114.15.2 | 107652 | Biolegend |
| Anti-mouse CD3 | FITC | 145-2C11 | 35-0031 | Tonbo Biosciences |
| Anti-mouse CD11b | Violet Fluor 450 | M1/70 | 75-0112 | Tonbo Biosciences |
| Anti-mouse CD27 | PE/Cyanine7 | LG.3A10 | 124216 | Biolegend |
| Anti-mouse NK1.1 | PE/Cyanine7 | PK136 | 50-5941 | Tonbo Biosciences |
| Anti-mouse CD45.2 | APC/Fire750 | 104 | 17-0454-81 | eBioscience |
| Anti-mouse CD64 | Alexa Fluor 647 | X54-5/7.1 | 558539 | BD Bioscience |
| Anti-mouse MHCII | FITC | 2G9 | 553623 | BD Bioscience |
| Anti-mouse CD11c | Brilliant Violet 421 | N418 | 565452 | BD Bioscience |
| Anti-mouse CD11b | PE/Cyanine7 | M1/70 | 25-0112-81 | eBioscience |
| Anti-mouse XCR1 | APC/Cyanine7 | ZET | 148224 | Biolegend |
| Anti-mouse Ly6C | PerCP-Cy5.5 | AL-21 | 560525 | BD Bioscience |
| Anti-mouse CD86 | PE | GL-1 | 50-0862 | Tonbo Biosciences |
| Anti-mouse CD3 | BV711 | 145-2C11 | 563123 | BD Bioscience |
| Anti-mouse CD107a | Alexa Fluor 647 | 1D4B | 121610 | Biolegend |
| Anti-mouse IFNγ | FITC | XMG1.2 | 11-7311-82 | eBioscience |
| Anti-mouse CD45.2 | BV786 | 107 | 563686 | BD Bioscience |
| Anti-mouse NKG2D | PerCPeFluor710 | CX5 | 46-5882-82 | ThermoFisher |
| Anti-human CD3 | Percp/Cy5.5 | HIT3a | 300327 | Biolegend |
| Anti-human CD19 | Percp/Cy5.5 | SJ25C1 | 363015 | Biolegend |
| Anti-human CD20 | Percp/Cy5.5 | 2H7 | 302325 | Biolegend |
| Anti-human CD56 | Percp/Cy5.5 | 5.1H11 | 362505 | Biolegend |
| Anti-human CD14 | APC Cy7 | MϕP9 | 557831 | BD Bioscience |
| Anti-human CD16 | PB | 3G8 | 302021 | Biolegend |
| Anti-human CD56 | PE | 5.1H11 | 362524 | Biolegend |
| Anti-human HLA-DR | PB | L243 | 307623 | Biolegend |
| Anti-human CD11c | PB | 3.9 | 301625 | Biolegend |
| Anti-human MICA/B | Alexa Fluor 488 | 6D4 | 320912 | Biolegend |
| Anti-human CD40 | FITC | 5C3 | 334306 | Biolegend |
| Anti-human CD86 | PE/Cy7 | BU63 | 374210 | Biolegend |
| Anti-human IFNγ | FITC | 4S.B3 | 554551 | Biolegend |
| Anti-human CD141 | APC | M80 | 344105 | Biolegend |
| Anti-human TNFα | Percp | MAB11 | 502923 | Biolegend |
| Anti-human CD107a | APC | H4A3 | 328620 | Biolegend |
| Anti-human CD1c | PE/Cy7 | L161 | 331515 | Biolegend |
| Anti-human CD64 | PE/Dazzle 594 | 10.1 | 305131 | Biolegend |
| Anti-human ULBP1 | PE | 170818 | FAB1380P | RyD systems |
| Anti-human NKG2C | APC | 134591 | FAB138A-025 | RyD systems |
| Anti-human CD1c | PE | F10/21A3 | 564900 | BD Bioscience |

| | | | | |
|---|---|---|---|---|
| Anti-human PCNA | Biotin | PC10 | 307904 | Biolegend |
| Anti-human HLA-F | APC | 3D11 | 373207 | Biolegend |
| Anti-human SLAMF7 | PE/Cy7 | 162.1 | 331815 | Biolegend |
| Streptavidin | V500 | n.a. | 561419 | BD Bioscience |
| Anti-human HLADR | APC/Cy7 | L243 | 307618 | Biolegend |
| Anti-human CD3 | APC/Cy7 | HIT3a | 300317 | Biolegend |
| Anti-human CD19 | APC/Cy7 | HIB19 | 302217 | Biolegend |
| Anti-human NKG2A | Percp | 131411 | FAB1059C-025 | RyD Systems |
| Anti-human NKG2D | FITC | 1D11 | 320819 | Biolegend |
| Anti-human NKp30 | PE/Cy7 | P30-15 | 325213 | Biolegend |
| Anti-human PD-L1 | Brilliant Violet 711 | 29E2A3 | 329721 | Biolegend |
| Anti-human CD14 | Alexa 700 | 63D3 | 367113 | Biolegend |
| Anti-human CD123 | APC | 7G3 | 560087 | BD Bioscience |
| Anti-human CD86 | BV650 | IT2.2 | 305427 | Biolegend |
| Anti-human CD16 | Brilliant Violet 785 | 3G8 | 302045 | Biolegend |
| Ghost DyeTM | Red 780 | n/a | 13-0865-T100 | Tonbo Biosciences |
| Yellow fluorescent reactive dye | BV405 | n/a | L34959 A | Invitrogen |
